# Working together to control mutation: how collective peroxide detoxification determines microbial mutation rate plasticity

**DOI:** 10.1101/2023.09.27.557722

**Authors:** Rowan Green, Hejie Wang, Carol Botchey, Nancy Zhang, Charles Wadsworth, Andrew J McBain, Pawel Paszek, Rok Krašovec, Christopher G Knight

**Author notes:** Corresponding Author (CGK); (RK).

## Abstract

Mutagenesis is responsive to many environmental factors. Evolution therefore depends on the environment not only for selection but also in determining the variation available in a population. One such environmental dependency is the inverse relationship between mutation rates and population density in many microbial species. Here we determine the mechanism responsible for this mutation rate plasticity. Using dynamical computational modelling and *in vivo* mutation rate estimation we show that the negative relationship between mutation rate and population density arises from the collective ability of microbial populations to control concentrations of hydrogen peroxide. We demonstrate a loss of this density-associated mutation rate plasticity when *Escherichia coli* populations are deficient in the degradation of hydrogen peroxide. We further show that the reduction in mutation rate in denser populations is restored in peroxide degradation-deficient cells by the presence of wild-type cells in a mixed population. Together, these model-guided experiments provide a mechanistic explanation for density-associated mutation rate plasticity, applicable across all domains of life, and frames mutation rate as a dynamic trait shaped by microbial community composition.

## Introduction

Uncovering the mechanisms behind environmentally responsive mutagenesis informs our understanding of evolution, notably antimicrobial resistance, where mutation supply can be critical (1, 2). Microbial mutation rates are responsive to a wide variety of environmental factors including population density (3), temperature (4), growth rate (5, 6), stress (7, 8), growth phase (9) and nutritional state (10). Such mutation rate plasticity inspires the idea of “anti-evolution drugs”, able to slow the evolution of antimicrobial resistance during the treatment of an infection (2, 11–13). Even small reductions in the mutation rate (2-5-fold) can have dramatic effects on the capacity of bacterial populations to adapt to antibiotic treatment, particularly when evolution is limited by mutation supply, as is the case for small pathogen populations (2).

Microbial mutation rates have an inverse association with population density across all domains of life, we have previously shown that 93% of otherwise unexplained variation in published mutation rate estimates is explained by the final population density (3). This density-associated mutation rate plasticity (DAMP) is a distinct phenotype from stress-induced mutagenesis, which acts via independent genetic mechanisms (14). Population density alters not only the rate but also the spectrum of mutations, with significantly higher rates of AT>GC transitions seen in low density populations (15). Density effects are likely relevant to natural populations given that population sizes and densities vary greatly, for example, *Escherichia coli* populations in host faeces can range in density by 5 orders of magnitude (16), and infections can be established by populations as small as 6×10^3^ cells (17). We therefore aim to mechanistically describe the widespread phenotype of DAMP.

In order to test potential mechanisms generating DAMP, we developed and systematically assessed a computational model connecting metabolism and mutagenesis in a growing *E. coli* population. This model generates the hypothesis that the key determinants of DAMP are the production and degradation rates of reactive oxygen species (ROS). Though molecular oxygen is relatively stable it can be reduced to superoxide (^•^O_2−_), hydrogen peroxide (H_2_O_2_) and hydroxyl radicals (HO^•^). These “reactive oxygen species” are strong oxidants able to damage multiple biological molecules including nucleotides and DNA (18). We tested the role of ROS in controlling DAMP by estimating mutation rate plasticity under different conditions of environmental oxygen and with genetic manipulations known to alter ROS dynamics. We find that the reduction in mutation rate at increased population density results from the population’s increased ability to degrade H_2_O_2_, resulting in reduced ROS-associated mutagenesis. We show that this density effect is also experienced by cells deficient in H_2_O_2_ degradation when cocultured with wild-type cells able to detoxify the environment. Mutation rates therefore depend not only on the genotype of the individual but also on the community’s capacity to degrade H_2_O_2_.

## Results

### Initial computational model of nucleotide metabolism in a growing microbial population fails to reproduce mutation rate plasticity

To generate hypotheses for the mechanisms of density-associated mutation rate plasticity we constructed a system of ordinary differential equations (ODEs) that recapitulates the dynamics of metabolism, growth and mutagenesis in a 1mL batch culture of *E. coli* (Fig. 1). The enzyme MutT, responsible for degrading mutagenic oxidised GTP (19), is essential in DAMP (3); the ODE model is therefore focussed on guanine bases. In the model external glucose (***eGlc***) is taken up by a small initial *E. coli* population (***wtCell***). Internal glucose (***iGlc***) is then metabolised to produce ***ROS*, *dGTP*** and, largely, ‘other’ molecules (‘Sink’ in Fig. 1). ***dGTP*** is then either integrated into a newly synthesised DNA molecule (***DNA***) or it reacts with ***ROS*** to produce 8-oxo-2’-deoxyguanosine triphosphate (***odGTP***). In this model, non-oxidised ***dGTP*** always pairs correctly with cytosine, producing non-mutant DNA (***DNA***). In a second round of DNA replication the guanine base is now on the template strand, cytosine is correctly inserted opposite producing new chromosomes (***wtCell***). ***odGTP***, if it is not dephosphorylated by MutT into *dGMP* (Sink), can either pair correctly with cytosine (becoming ***DNA***) or mis-pair with adenine (becoming ***mDNA***). When ***odGTP*** is inserted opposite adenine into DNA (***mDNA***) it may be repaired by the MutS or MutY proteins, converting the ***mDNA*** back to ***DNA***. The key output of interest is the mutation rate, which is defined as the number of mutant base pairs (***mCell***) divided by the number of non-mutant base pairs (***wtCell***). The model comprises 10 ordinary differential equations (ODEs), one for each substance variable in Fig.1 (excluding ‘Sink’), plus ***cytVol***, the total population cytoplasmic volume within which all the reactions occur (Table 1, Eq. 1-10, Methods). These equations require 14 parameters (some of them composite, Table 2); the structure and parameter values are largely taken from the existing literature (for details see Methods). Un-measurable parameters (notably the rate of ***dGTP*** oxidation to ***odGTP*** by ***ROS***, ‘**O2’**) were set to give the observed mutation rate (2 × 10^-10^ mutations per base pair per generation, (20)) at a final population density of 3 × 10^8^ CFU ml^-1^, typical of 250 mg L^-1^ glucose in minimal media. As with most experiments demonstrating density-associated mutation rate plasticity (3, 21), final population density is controlled by varying initial external glucose. We initiated 28h simulations of 1ml cultures with 2175 cells (a small number, typical of fluctuation assays estimating mutation rate, Fig. S10), no internal metabolites and external glucose concentrations relevant to wet-lab experiments – across a log scale from 55 to 1100 mg L^-1^ (Table 1). The dynamics of external glucose, population size and mutation rate for these simulations are shown in Fig.1B-D.

**Figure 1:**
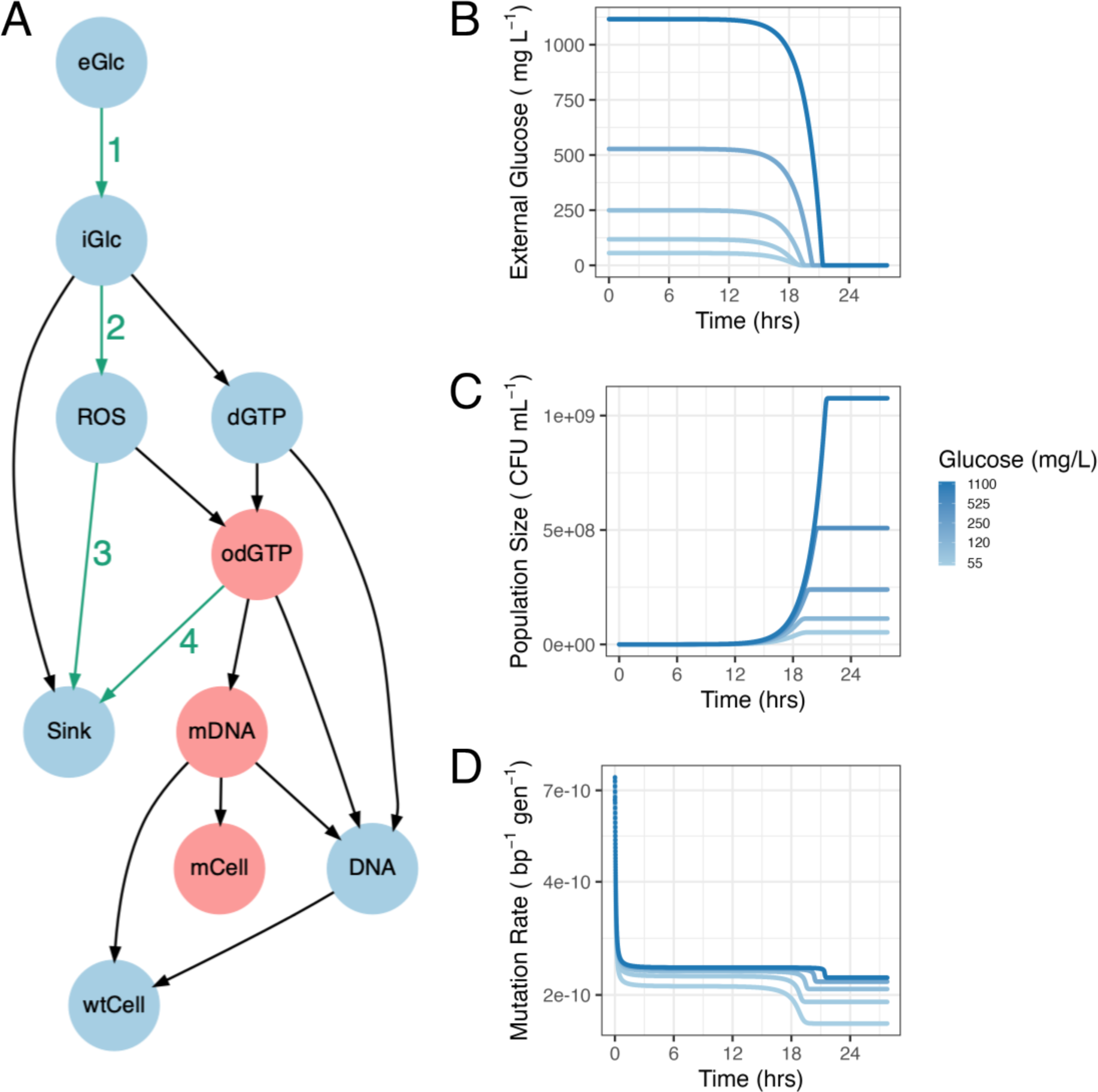
Dynamical computational model of growth, metabolism and mutagenesis in E. coli. A: Model structure connecting variables. Red variables indicate the pathway to mutagenesis; green numbered arrows indicate pathways targeted by model variants. This structure was represented in ODEs, parameterised from the literature (Methods), and simulated to give output shown in B-D. B: Kinetics of external glucose concentration (eGlc), C: population size (wtCell divided by G nucleotides in the E. coli genome) and D: mutation rate (mCell/wtCell). Note log scale on y-axis in panel D. Panels B-D are plotted for 5 initial glucose concentrations (range 55 – 1100 mg L_-1_ as shown in legend), initial glucose concentration indicated by line colour.

**Table 1:**
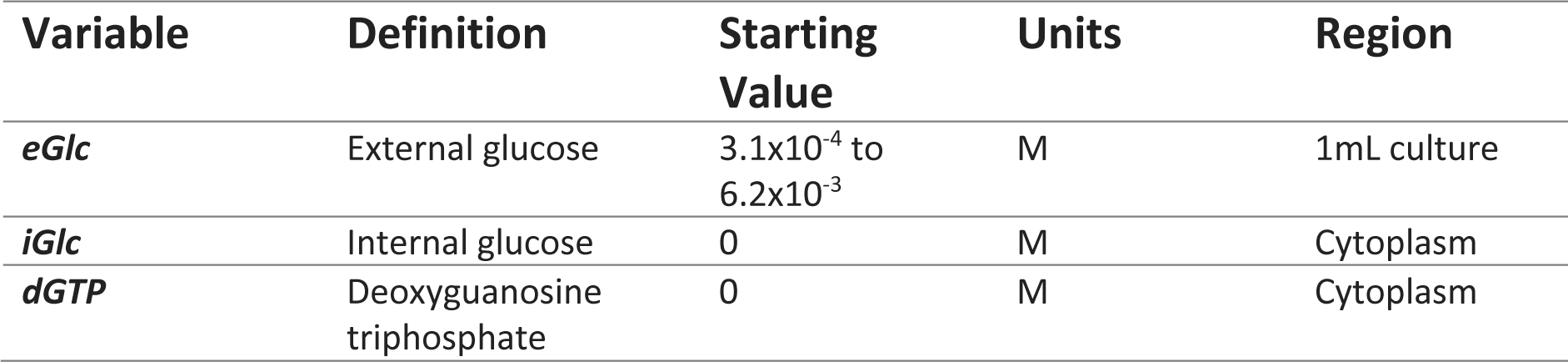
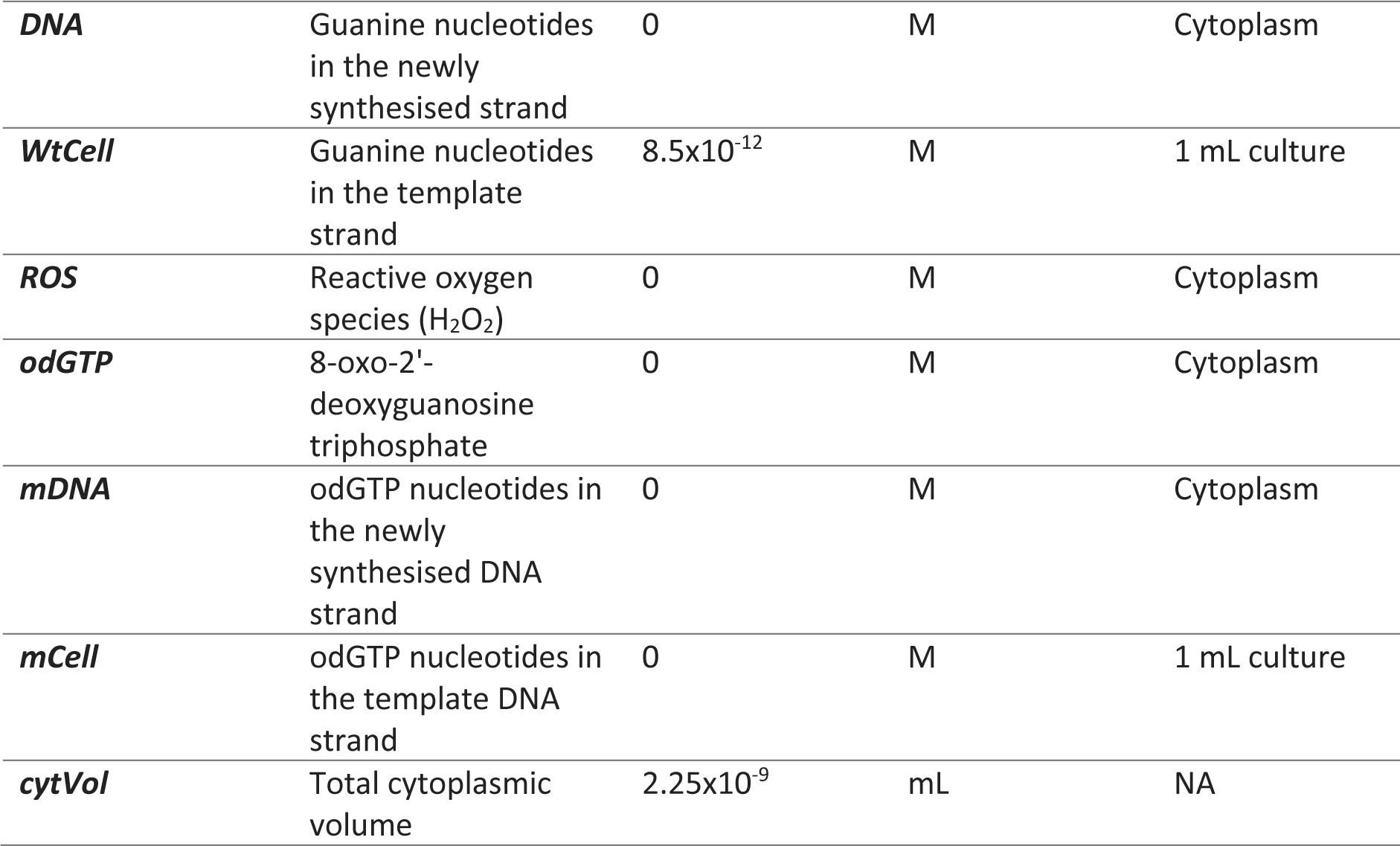
Definitions and starting values for the 10 variables in ODE model A. For variables measured as a concentration, the volume within which this is calculated is given in the ‘region’ column. wtCell and cytVol starting values equate to 2175 cells (assuming 2357528 GC bp in the E. coli genome (strain MG1655, EBI Accession U00096.3) and cell volume of 1.03×10_-12_ mL (72)).

**Table 2.** Parameter values and descriptions for all parameters used in model A.

| Parameter | Value | Units | Description | Source |
| --- | --- | --- | --- | --- |
| **U1** | 2.66E-01 | $M^{-1}s^{-1}$ | Maximum uptake rate ($V_{max}$) of eGlc | Fitted from a known value of $K_s$ (80) and data on *E. coli* growth dynamics (81) (Fig. S14). |
| **M1** | 2.69E-04 | $s^{-1}$ | Rate of dGTP synthesis from iGlc | This value was fitted to published data on *E. coli* growth dynamics (81) (Fig. S14). |
| **Ks** | 3.97E-05 | M | Michaelis<br>Menten<br>constant $K_s$:<br>Concentration<br>of glucose at<br>which glucose<br>uptake rate of<br>1/2 $V_{max}$ is<br>achieved | $K_s$ measured as 7.16 $\mu\text{g/ml}$ by (80). |
| **I1** | 6.90E-03 | $\text{s}^{-1}$ | Rate of dGTP<br>incorporation<br>into DNA<br>opposite C | Fitted to give known cytoplasmic<br>concentration dGTP in exponential<br>growth phase *E. coli* of 92 $\mu\text{M}$ in 0.4%<br>glucose (82). |
| **D1** | 6.90E-03 | $\text{s}^{-1}$ | Rate of C<br>pairing<br>opposite<br>incorporated<br>G | Given the same value as I1 as the<br>synthesis of new DNA (**DNA**) and new<br>genomes (**wtCell**) should generally<br>proceed at equal rates (this is violated<br>during rapid exponential growth (83)<br>but not included in this model). |
| **O2** | 12.0 | $\text{M}^{-1}\text{s}^{-1}$ | Rate of dGTP<br>oxidation to<br>odGTP by ROS | Selected to give a mutation rate of<br>$1.94 \times 10^{-10}$ base pair substitutions per<br>nucleotide in 0.2% glucose minimal<br>media (20). |
| **I2** | 2.53E-04 | $\text{s}^{-1}$ | Rate of odGTP<br>incorporation<br>into DNA<br>opposite<br>adenine | The relative efficiency of odGTP<br>binding to A (I2) compared to G<br>binding to C (I1) is $7.7 \times 10^{-8} : 2.1 \times 10^{-6}$<br>(19) therefore $I2 = I1 * (7.7 \times 10^{-8} / 2.1 \times 10^{-6})$. |
| **D2** | 2.00E-04 | $\text{s}^{-1}$ | Rate of C<br>pairing<br>opposite<br>incorporated<br>oG causing an<br>AT>CG<br>mutation | Rate of C pairing opposite an odGTP<br>(D2) relative to CG (I1 & D1) is $6 \times 10^{-8} : 2.1 \times 10^{-6}$ (19). |
| **C1** | 2.8 | $\text{s}^{-1}$ | Rate of odGTP<br>hydrolysis to<br>odGMP by<br>nudix<br>hydrolase<br>enzyme MutT<br>(NudA) | Value taken from Kcat of MutT<br>measured *in vitro* (84). |
| **C2** | 3.50E-04 | $\text{s}^{-1}$ | Rate of<br>removal of<br>adenine base<br>incorporated<br>opposite 8-oxodG in the<br>genome by<br>enzyme MutY | Value taken from Kcat of MutY<br>measured *in vitro* as $0.021 \text{ min}^{-1} = 3.5 \times 10^{-4} \text{ sec}^{-1}$ (85). |
| **R1** | 2.00E-04 | s<sup>-1</sup> | Rate of oG insertion into DNA opposite cytosine | Relative incorporation efficiency opposite C of odGTP:dGTP (R1) is 0.029 (19), therefore R1 = I1 * 0.029 . |
| **S** | 2.58E-02 | s<sup>-1</sup> | Rate of removal of adenine base incorporated opposite 8-oxodG in the genome by enzyme MutS | Fitted to known rate of mutation in *mutS* knockout of 40x wild-type (86). |
| **r** | 17.3 | s<sup>-1</sup> | Rate of ROS production from iGlc relative to dGTP production rate | Selected to a known H<sub>2</sub>O<sub>2</sub> production rate of 14 µM/second in 0.2% glucose minimal media normalised to cell volume (24). |
| **O3** | 5.60E+01 | s<sup>-1</sup> | Rate of ROS degradation through reactions other than dGTP oxidation | Fitted to give a standing ROS concentration of 1.9x10<sup>-7</sup> M, midpoint of known 1.3x10<sup>-7</sup> - 2.5x10<sup>-7</sup> M in LB (87). |
| **R2** | 2.53E-04 | s<sup>-1</sup> | Rate of adenine pairing opposite incorporated oG | The relative efficiency of odGTP binding to A (R2) compared to G binding to C (I1) is 7.7x10<sup>-8</sup> : 2.1x10<sup>-6</sup> (19) therefore R2 = I1 * (7.7x10<sup>-8</sup>/2.1x10<sup>-6</sup>). |
| **Met1** | 1545 | s<sup>-1</sup> | Stoichiometry of glucose conversion to dGTP for genome building (i.e. how many molecules of glucose are needed to produce 1 molecule of dGTP) | Fitted to published data from (3) of cell density as a product of glucose concentration (Fig. S15). |
| **CellVol** | 1.03E-12 | mL | Volume of one *E. coli* cell in minimal media growing in exponential<br>phase | Mean value of 4 estimates of cell volume in exponential phase cells grown in minimal M9 media (72). |
| **molML** | 6.02E+20 | molecules | Number of molecules per mL in a 1M solution. | One thousandth of Avogadro's constant ($N_A$). |
| **GCperGen** | 2357528 | GC basepairs | Number of GC basepairs per genome. | *E. coli* Strain MG1655, EBI Accession U00096.3 |

This initial model (Fig 1, referred to as model A) creates an approximately linear log-log slope of 0.09 ± 0.06 (95% CI) between final population and mutation rate (Red line, Fig.2A, Regression 1 (SI)). We can compare the slope directly to *in vivo* estimates of mutation rates in *E. coli*, which show strong DAMP, with a slope of -0.84 ± 0.13 (95% CI, grey dots and dashed line, Fig. 2A, Regression 2 (SI)). Model A is therefore not describing the processes causing DAMP – the structure and/or the parameters used are either incomplete or fail to replicate biology for some other reason. To test whether inappropriate parameter values could be responsible for the lack of DAMP in model A, we simulated 50,000 parameter sets simultaneously varying all parameters randomly across 10% – 1,000% of their original value. These results were filtered as described in Methods and are plotted in Fig. 2B. This global sensitivity analysis showed the mutation rate plasticity, i.e., slope of model A to be very robust, with an interquartile range of 0.02-0.13 as shown by error bars in Fig.2B. All tested parameter sets gave a log-log linear slope of > - 0.05, suggesting that DAMP requires processes not represented in this initial model.

**Figure 2:**
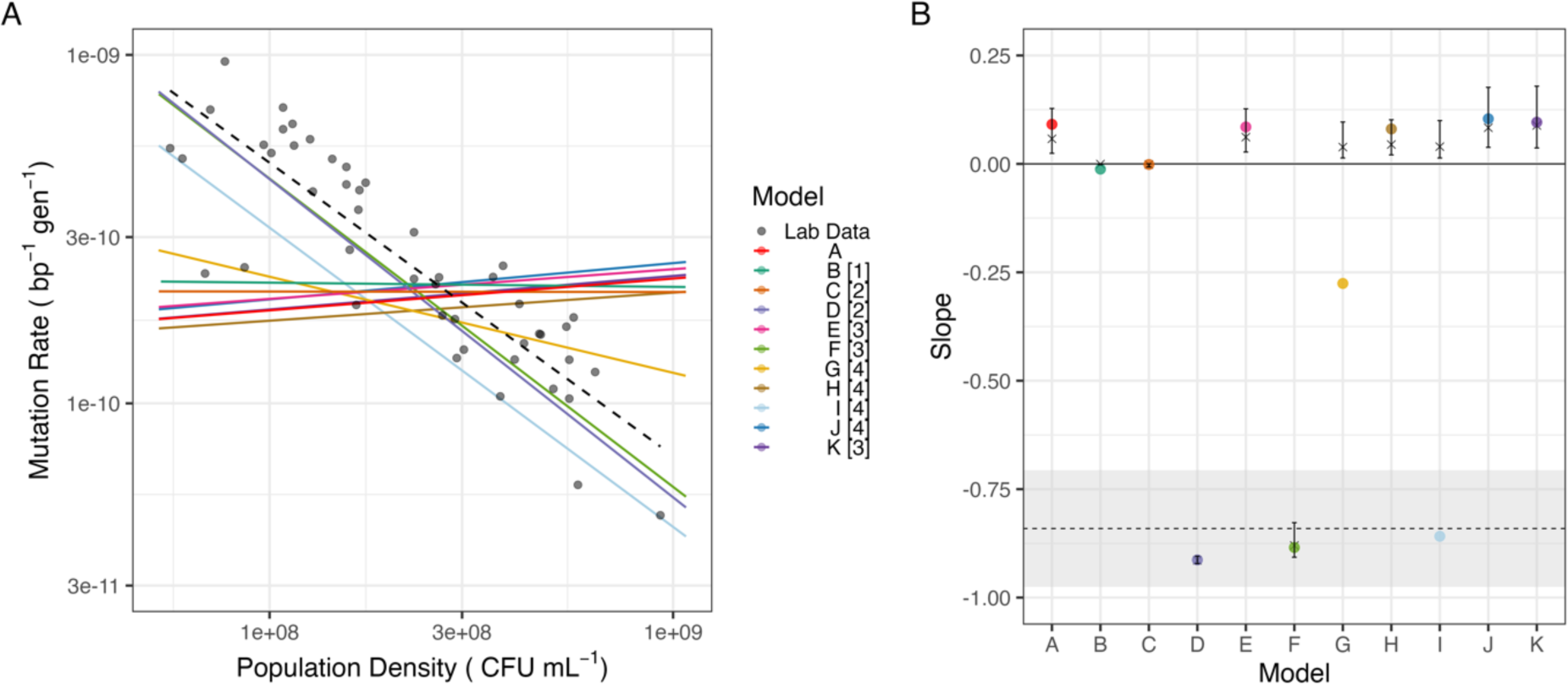
Mutation rates in model variants. A) Solid coloured lines show fitted log-log linear relationship between final population density and mutation rate for models A to K (Regression 1 (SI)); numbers [1] – [4] in legend indicate the pathway targeted from Fig. 1A. Black points and dashed line show lab data for E. coli wild-type BW25113 in glucose minimal media with a log-log linear regression fitted (Regression 2 (SI)). B) Global sensitivity analysis; coloured points show slopes from baseline parameters (as in 2A), and crosses and error bars show median and interquartile range of slope across 5×10_5_ randomly perturbed parameter sets, models are coloured as in Fig. 2A. Dashed line shows slope of lab data in Fig. 2A (Regression 2), and grey area shows 95% CI on this slope.

### ROS production and degradation are central to density-associated mutation rate (DAMP) plasticity *in silico*

While there was only limited variation in the relationship between mutation rate and population density, defining the slope of DAMP, in model A (Fig. 2B) we can ask which model parameters are associated either with this variation or with variation in mutation rate itself (Fig. S1). The affinity of importers for glucose (Ks, part of reaction 1 in Fig. 1A) had by far the closest association with the slope (Spearman’s Rho _(*DF* = 3583)_ = 0.91, *P* < 2.2 × 10^-16^, Fig. S1), whereas a group of parameters, including parameters controlling the rates of both ***ROS*** production (r, reaction 2 in Fig. 1A) and ***ROS*** degradation (parameters O2 and O3, corresponding to reaction 3 in Fig. 1A) had the closest association with the mutation rate (Spearman’s Rho _(*DF* = 3583)_ = 0.22, 0.21 and -0.21 respectively, all *P* < 2.2 × 10^-16^ Fig. S1). The parameter representing MutT activity (parameter C1, reaction 4, Fig. 1A), found to be relevant in previous work on DAMP (3), was also in this group of parameters controlling mutation rate and so was also considered as candidate processes for further exploration. We hypothesise that the additional processes required to reproduce DAMP as observed in the lab are associated with these reactions (numbered 1-4 in Fig. 1A). We systematically tested each of these processes using structural variants to the model, explicitly modifying density dependence in biologically plausible ways. We thus use these models as a method of hypothesis generation, to determine which mechanisms may plausibly cause DAMP, with a view to testing these candidate mechanisms in the lab.

The slight increase in mutation rates seen as density increases in model A (a reversal of the negative association seen in the DAMP phenotype, therefore referred to as ‘reverse DAMP’) is the result of increased external glucose leading to increased internal glucose concentrations (Fig. S2), creating an increased rate of ***ROS*** production and therefore higher mutation rates. It is therefore plausible that if glucose importer proteins are more expressed under low external glucose conditions, increasing the rate of reaction 1 (Fig. 1A) at low glucose concentrations may increase mutation rates at low density. Introducing this model variant (model B, using Eq. 1_B_) does indeed remove model A’s positive association between mutation rate and density but does not give the negative association observed *in vivo* (model B slope = -0.01 ± 0.06 (95% CI), Fig. 2, Regression 1 (SI)).

In model A, ***ROS*** are produced only by cellular metabolism, however lab media also accumulates significant concentrations of H_2_O_2_ through photochemistry (22). This is represented in model C by replacing reaction 2 (Fig. 1A) with a constant ***ROS*** concentration in the system (using Eq. 8_C_ rather than Eq. 8) and in model D by a constant rate of ***ROS*** production (using Eq. 7_D_ rather than Eq. 7). Both models abolish model A’s positive slope. However, while model C removes DAMP (slope = -0.001 ± 0.06 (95% CI), Regression 1 (SI)), model D introduces a strong negative slope similar to the laboratory data (slope = -0.91 ± 0.06 (95% CI), Regression 1 (SI)).

Decreasing mutation rates at higher population densities could also be the result of changes in cellular ROS degradation rates (reaction 3). We therefore created models where degradation is determined by the internal glucose concentration (model E) and by the population density (model F), replacing Eq. 7 with Eq. 7_E_ and 7_F_ respectively. Of these two, the first had very little effect (model E, slope = 0.09 ± 0.06 (95% CI), Regression 1 (SI)) whereas the second had a large effect, giving a strong slope similar to the laboratory data (model F, slope = -0.89 ± 0.06 (95% CI), Regression 1 (SI)).

Given that previous work has shown the action of MutT in degrading ROS-damaged dGTP (***odGTP***, Fig. 1A) to be essential to DAMP (3), we explored models in which the rate of ***odGTP*** degradation by MutT (reaction 4, Fig. 1A) is determined by the internal glucose (model G, using Eq. 8_G_ rather than Eq. 8), ***odGTP*** (model H, using Eq. 8_H_ rather than Eq. 8) or ***ROS*** concentration (model I, using Eq. 8_I_ rather than Eq. 8). None of these models consistently resulted in DAMP (Fig. 2B): making MutT activity dependent on ***odGTP*** had very little effect at all (model H, slope = 0.08 ± 0.06 (95% CI), Regression 1 (SI)) whereas making MutT activity directly responsive to internal glucose or ROS concentration did reproduce some degree of DAMP slope (models G and I slopes -0.28 ± 0.06 and - 0.86 ± 0.06 respectively (95% CI), Regression 1 (SI)). However, the DAMP slopes of models G and I are highly parameter dependent with the majority of parameter combinations in the global sensitivity analysis giving very little slope at all (Fig. 2B).

Finally, we replaced model A’s mass action dynamics with saturating Michaelis Menten kinetics for MutT activity (reaction 4, model J using Eq. 7_J_ rather than Eq. 7 (19)) and enzymatic degradation of H_2_O_2_ (reaction 3, model K using Eq.7_KA_ and Eq.7_KB_ rather than Eq. 7, (23)). Neither of these modifications greatly affected the mutation rate response of the model to population density (slope = 0.104 ± 0.06 and 0.096 ± 0.06 respectively, (95% CI), Regression 1 (SI), Fig. 2). Thus, across 11 biologically plausible model structures only two, D and F, affecting reactions 2 and 3 respectively in specific ways, produced DAMP comparable to that observed in the laboratory (Fig. 2A) and robust to parameter variations (Fig. 2B).

We can use these model findings for hypothesis generation: Model A (without DAMP) only describes ROS production from metabolism, whereas Model D (with DAMP) modifies the initial model to have a constant rate of ROS generation, independent of the cell density. Model D is consistent with ROS production in the system being dominated by environmental sources at a constant rate. If DAMP is a result of such environmental ROS production, we would expect this phenotype to be absent under anaerobic conditions where external H_2_O_2_ production is negligible (22).

Model F, which gains DAMP relative to model A, describes an increased rate of ROS detoxification dependent on the population density. This reflects a system in which ROS detoxification is primarily occurring within cells. Here ROS diffusion into cells from the environment is significant and therefore the environment is more efficiently detoxified by larger populations. If DAMP is a result of an increased environmental detoxification capacity in dense populations in this way, we expect strains deficient in ROS degradation not to show DAMP. We would further expect dense populations to show greater removal of environmental ROS than low-density populations.

We therefore go on to test these predictions *in vivo* using fluctuation assays to estimate the mutation rate in batch cultures of *E. coli*.

### Environmental oxygen is necessary for DAMP *in vivo*

To test the hypothesis (from model D) that DAMP is dependent on external oxygen (from model D) we estimated mutation rates of *E. coli* under anaerobiosis across a range of nutrient-determined final population densities, analysing the results using a linear mixed effects model (Regression 4 (SI)). We find that anaerobic growth results in a loss of the negative relationship between density and mutation rate, indeed mutation rates significantly increased with density (slope = 0.6 ± 0.42 (95% CI), Fig. 3B, statistical tests in Table S1, Regression 4 (SI)). We further test this relationship using a second wild-type strain (*E. coli* MG1655). Again, we see a loss of DAMP under anaerobiosis (slope = 0.12 ± 0.7 (95% CI), cf. slope = -0.43 ± 0.25 (95%CI), anaerobic and aerobic respectively, Regression 4 (SI), Fig. S3). This supports the hypothesis arising from model D that, when external ROS production is substantial (model D / aerobiosis) mutation rates fall with increasing final population size, whilst when external ROS production is not included (model A / anaerobiosis) mutation rates remain similar or increase slightly with higher cell densities.

**Figure 3:**
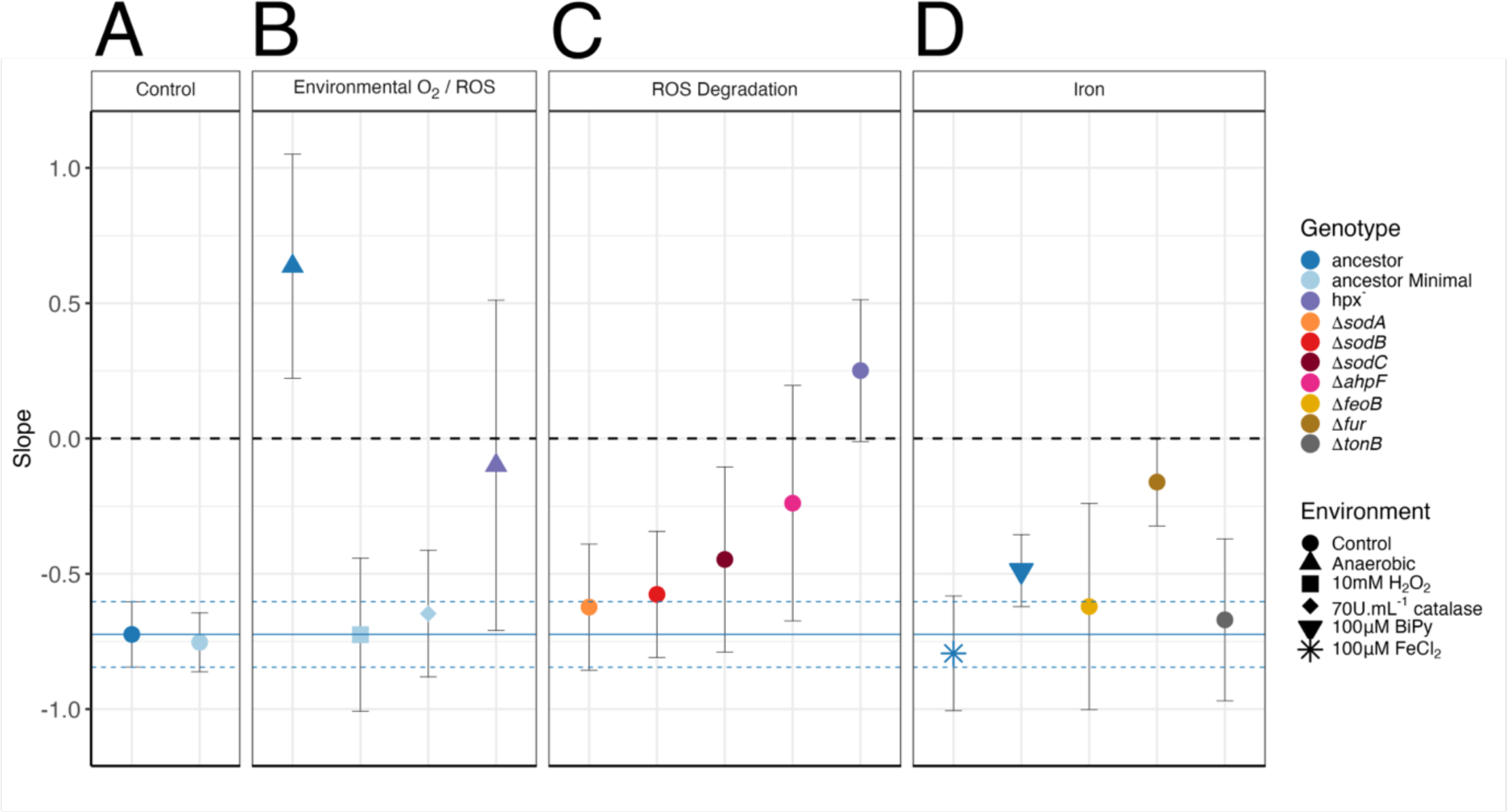
Mutation rate responses to population density in vivo under environmental and genetic manipulations. Points show the slope of a log-log relationship between final population size and mutation rate (raw data shown in Fig. S4, Regression 4 (SI)), error bars show 95% CI on slope. Treatments shown are BW25113 ancestor (1106 parallel cultures (pc) across 69 fluctuation assays (fa)); ancestor minimal media (942 pc, 59 fa); ΔahpF (266 pc, 17 fa); hpx_-_ (402 pc, 26 fa); ancestor anaerobic (168 pc, 11 fa); ancestor 10mM H_2_O_2_ (179 pc, 12 fa); ancestor 70U mL^-1^ catalase (167 pc, 11 fa); hpx^-^ anaerobic (105 pc, 7 fa); ancestor + chelator 2,2,Bipyridyl 100µM (382 pc, 24 fa); ancestor + FeCl_2_ 100µM (210 pc, 13 fa); ΔfeoB (192 pc, 12 fa); Δfur (504 pc, 31 fa); ΔtonB (113 pc, 7 fa). Dashed line shows a slope of 0 (no DAMP); solid blue line shows the slope of BW25113 ancestor in rich media with dashed blue lines showing 95% CI on this estimate (Regression 4 (SI)). All experiments were conducted in dilute LB media unless stated ‘Minimal’, in which case glucose minimal media was used.

### Endogenous ROS degradation is necessary for DAMP *in vivo*

The second ODE model able to reproduce DAMP (model F) introduces increased rates of ROS degradation with increasing population density. If DAMP is the result of active cellular ROS degradation, we would expect strains deficient in this trait to lack DAMP. The two alkyl hydroperoxide reductase subunits AhpC and AhpF are together responsible for the majority of H_2_O_2_ scavenging in aerobically growing *E. coli* (24). The remaining H_2_O_2_ is degraded by the catalase enzymes HPI (*katG*) and HPII (*katE*) (25). The role of catalases in H_2_O_2_ scavenging is much more significant at high H_2_O_2_ concentrations due to the higher Michaelis constants of these enzymes, whereas AhpCF is saturated at ∼20µM (25). We therefore estimated DAMP in a version of the *E. coli* MG1655 strain lacking *ahpC*, *ahpF*, *katG* and *katE* (hpx^-^, (26)). This quadruple deletion results in a complete loss of DAMP with no significant change in the mutation rate across densities (slope = 0.25 ± 0.26 (95% CI), Fig. 3C, Regression 4 (SI)). Enzymatic degradation of H_2_O_2_ is thus essential to the DAMP phenotype, consistent with model F. Deleting only *ahpF* gives an intermediate DAMP phenotype (slope = -0.24 ± 0.44 (95% CI), Fig. 3C, Regression 4 (SI)) with significantly weaker DAMP than the wild-type (LR = 4.9, *P* = 0.028, Regression 4 (SI)), but still retaining stronger DAMP than hpx^-^ (LR = 4.1, *P* = 0.043, Regression 4 (SI)), indicating that DAMP requires both catalase and alkyl-hydroperoxide reductase activity. In contrast, individual knockouts affecting superoxide rather than H_2_O_2_ (the superoxide dismutase genes *sodA, sodB and sodC,* slope = -0.62 ± 0.23, -0.58 ± 0.23, and - 0.45 ± 0.34, respectively (95% CI), Fig. 3C, Regression 4 (SI)), or adding environmental H_2_O_2_ or catalase (slope = -0.73 ± 0.28, -0.65 ± 0.23, respectively (95% CI), Fig.3B, Regression 4 (SI)) do not significantly disrupt the wild-type negative relationship between population density and mutation rate (Table S1).

If the DAMP reproduced by model F is biologically realistic in this way, it requires that high-density populations, exhibiting reduced mutation rates, show greater efficiency at removing H_2_O_2_ from their environment than low-density populations. We measured external H_2_O_2_ in cultures after 24 hours of growth in rich or minimal media and found high-density populations to achieve significantly lower H_2_O_2_ concentrations (*F*_28_ = 24.3, *P*=3.3×10^-5^, Regression 7B (SI), Fig. S6); there was no significant effect of rich versus minimal media (*F*_26_ = 0.77, *P=*0.39, Regression 7A (SI)). The reverse pattern is seen in sterile media where increasing nutrient provision leads to increased H_2_O_2_ concentration (*F*_46_ = 9.8, *P* = 3×10^-3^, Regression 6 (SI)). This supports the hypothesis that, as required by model F, high-density populations detoxify external H_2_O_2_ better than low-density populations. However, while they are necessary, it does not require that all external H_2_O_2_ present in sterile media is degraded by alkyl-hydroperoxidase and catalase – other molecules, notably pyruvate, are excreted by *E. coli* with a substantial capacity for H_2_O_2_ degradation (27–29).

### Cellular iron regulation is required for DAMP

Our model-guided hypothesis testing has shown that DAMP requires H_2_O_2_. Our models involve the direct effect of ROS on DNA, however, it’s the reaction of free Fe(II) with H_2_O_2_ to produce mutagenic OH. radicals, Fenton chemistry, which is a major source of oxidative stress in *E. coli* (30, 31). These radicals are far more reactive and damaging to DNA than H_2_O_2_ itself, making iron critical to determining the amount of damage H_2_O_2_ causes (32). If DAMP’s dependence on H_2_O_2,_ is the result of variable oxidative damage to DNA and nucleotides, we would expect this mutation rate plasticity to be perturbed by changes in cellular iron homeostasis. We first tested this using environmental manipulations of iron. However, the provision of FeCl_2_ or starving cells of iron with a chelator (2,2-bipyridyl), has little effect on DAMP (Fig. 3D). Nonetheless we find that a deletant of *fur*, the master regulator of intracellular iron, results in an almost constant mutation rate across cell densities, with a significant reduction in DAMP compared to the BW25113 wild-type (slope = -0.16 ± 0.16 (95% CI); wt slope comparison: *LR* = 30.2, *P =* 3.9×10^-8^, Regression 4 (SI)). Fur is a negative regulator of multiple iron importers (33), therefore in Δ*fur* strains the internal redox-active iron pool is elevated (34) leading to increased oxidative stress and DNA damage (35). Knockouts of the iron importer genes *feoB* and *tonB*, which, if anything, reduce intracellular iron (36, 37), do not lead to any change in mutation rate plasticity (Fig. 3D, Table S1), likely because regulators such as Fur are able to maintain iron homeostasis in the absence of these individual importers.

The critical contribution of iron to H_2_O_2_ stress is further demonstrated through whole genome sequencing of the hpx^-^ and strain used here. We find a 190bp loss-of-function mutation in the iron importer *fecD* (all mutations listed in Table. S2). This may have allowed this hpx^-^ strain to escape the positive feedback cycle that hpx^-^ cells experience, in which higher H_2_O_2_ concentrations prevent Fur from effectively limiting iron uptake, more intracellular free iron then further exacerbates the damage done by the excess H_2_O_2_ (38, 39). It is likely that this loss-of-function mutation is an adaptation, during laboratory culture, to the loss of Fur functionality caused by the oxidation of intracellular iron.

### Wild-type cells restore DAMP in cells deficient in peroxide degradation

We have identified DAMP as requiring environmental H_2_O_2_ and iron regulation, where, with wild-type iron regulation, less H_2_O_2_ leads to lower mutation rates at higher final cell densities. This understanding leads us to predict that the presence of wild-type cells should restore DAMP in the peroxidase and catalase deficient hpx^-^ strain. It has previously been shown that a wild-type population can provide protection against environmental H_2_O_2_ to cocultured hpx^-^ cells, or similarly H_2_O_2_ sensitive Δ*oxyR* cells, through decreasing the peroxide concentration of the external environment (24, 40). To better distinguish hpx^-^ and wild-type strains in a coculture, two nalidixic acid (Nal) resistant strains of hpx^-^ were independently created with the resistance conferred by point mutations in *gyrA* (D87G & D87Y). Coculturing these hpx^-^ _nalR_ strains with wild-type BW25113 cells, the loss of DAMP via the hpx^-^ mutation (Fig. 3C) is phenotypically complemented by the wild-type cells. That is, hpx^-^ _nalR_ mutation rate is significantly decreased in coculture with increasing population density either of the hpx^-^ strain (Fig. 4, Fig. S7, slope = -0.93 ± 0.54 (95% CI), Χ² _(*DF*=1)_= 10.9, *P =* 9.8×10^-4^, Regression 4 (SI)), or total population density (Fig. S8, Slope = -1.4 ± 0.98 (95% CI), t_29_ = -2.8, *P* = 9×10^-3^, Regression 8 (SI)). This supports the hypothesis that DAMP is the result of reduced environmental H_2_O_2_ concentrations achieved by the local wild-type population.

**Figure 4:**
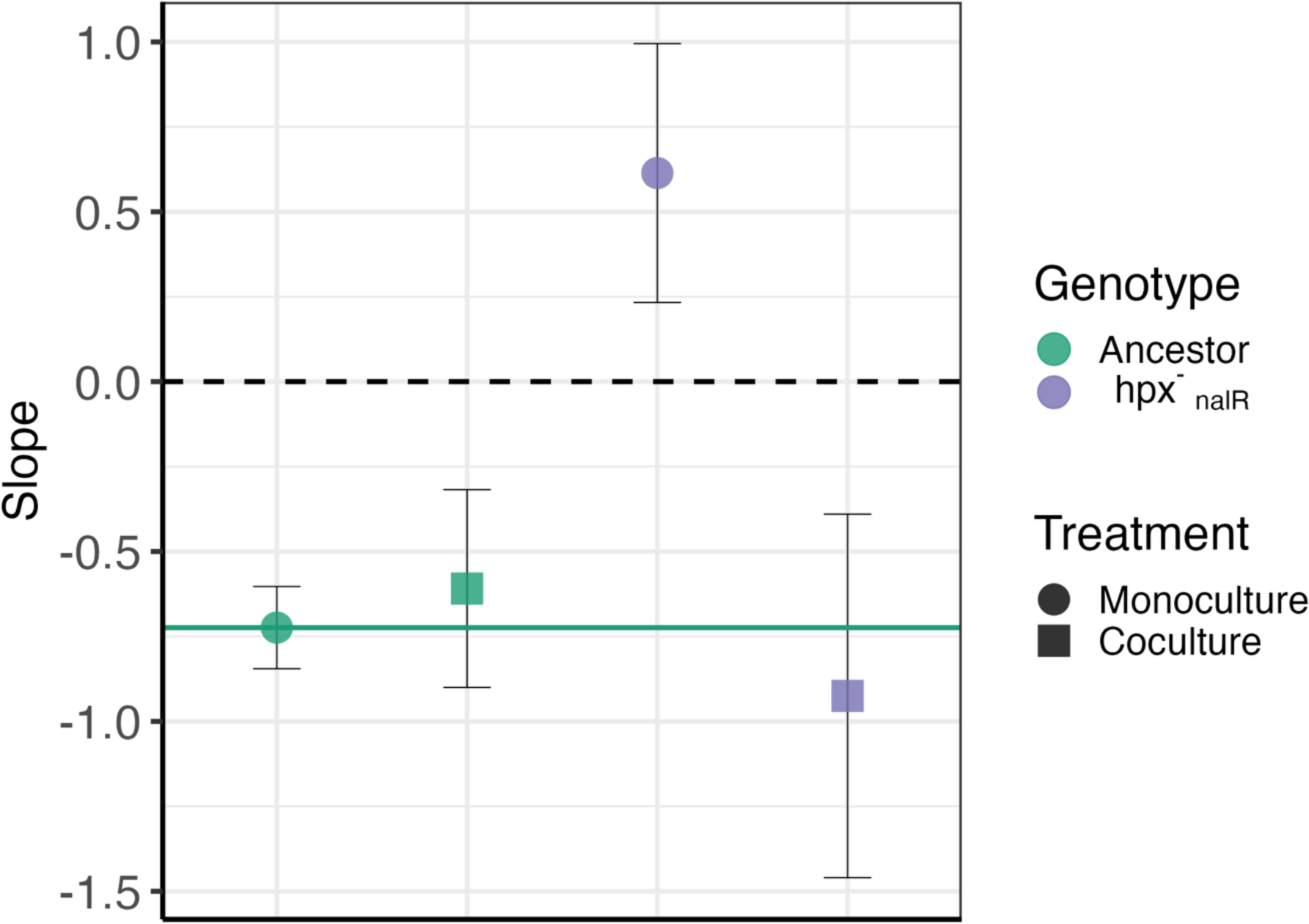
Coculture with wild-type cells restores DAMP in cells deficient in peroxide degradation: points show log-log relationship between final population density of the focal strain and mutation rate fitted by regression 4 (SI) (raw data shown in Fig. S7), error bars show 95% CI on slope. We found no significant differences between independent hpx^-^_nalR_ strains, therefore hpx^-^ strains D87Y & D87G are combined in hpx^-^_nalR_. Treatments shown are: BW25113 ancestor (1106 pc, 69 fa); BW25113 in coculture with hpx^-^ (498 pc, 31 fa); hpx^-^_nalR_ (388 pc, 24 fa); hpx^-^_nalR_ in coculture with BW25113 (319 pc, 20 fa).

Such mutation rate estimates in coculture could potentially be confounded by differential survival of rifampicin-resistant (RifR) mutants of hpx^-^_nalR_ when plated in a monoculture or a coculture. In order to test for any differences in mutant survival we conducted a ‘reconstruction test’ (as in Fig. S13 of (21)); plating a predetermined number of hpx^-^_nalR&rifR_ cells with a population of rifampicin susceptible hpx^-^ or wild-type cells on the selective rifampicin agar. No significant difference in plating efficiency was seen between plating with hpx^-^ vs. low, medium or high density of wild-type cells (Fig. S7; LR_7_ = 0.3, *P* = 0.96, Regression 9 (SI)). Some difference in plating efficiency between the two hpx^-^ _nalR&rifR_ strains was observed (Fig. S9), this is likely due to the pleiotropic effects of RifR resistance mutations in the *rpoB* gene (41, 42).

Although the wild-type strain reintroduces a negative dependence of mutation rates on population density in hpx^-^_nalR_ we also observe an increase in total hpx^-^_nalR_ mutation rates (Fig. S8). This is potentially the result of out-competition by the wild-type strain leading hpx^-^_nalR_ growth to stop earlier in the culture cycle where, consistent with previous fluctuation results ((43), Chapter 5.4.3), our modelling leads us to expect higher mutation rates (Fig. 1D).

## Discussion

Using ODE modelling (Fig. 1-2) to guide *in vivo* experiments in *E. coli* we have been able to predict and demonstrate the mechanisms behind the widespread phenomenon of reduced mutation rates at high microbial population densities (density-associated mutation rate plasticity, DAMP (3)). Genetic and environmental manipulations show that DAMP results from the improved degradation of H_2_O_2_ as the population density is increased (Fig. 3). The reintroduction of DAMP in catalase/peroxidase deficient cells by coculture with wild-type cells (Fig. 4) demonstrates the importance not only of a microbe’s own population density in determining the mutation rate but also the density and genotype of coexisting populations. Our results demonstrate that mutation rates can be context dependent, through the degradation capacity of a community for mutagens including, but perhaps not limited to, H_2_O_2_. This collective protection from harmful molecules mirrors studies such as (44), demonstrating the importance of population structure in microbial ecology.

Increased population density provides protection against high levels of external H_2_O_2_ stress (40, 45, 46). However, the concentrations of 100µM-1mM applied in such studies is far beyond the range of known environmental concentrations, which is typically up to only 4μM (47). Here we show that without any external input of H_2_O_2_ higher density populations detoxify environmental H_2_O_2_ more effectively over 24 hours than low-density populations (Fig. S6). As well as improving survival under extreme H_2_O_2_ stress, previous work also finds mutation rates to decrease in cells protected by a higher density of neighbours able to detoxify the environment (40). Here we find that this mutation protection holds in the absence of external H_2_O_2_ application with the presence of higher density wild-type rescuers able to modify mutation rates in catalase/peroxidase deficient cells (Fig. 4). This interaction between two *E. coli* strains raises the question of whether similar effects will be seen in mixed species communities such as human microbiomes where mutations can be critical for medically important traits, such as antimicrobial resistance (48, 49). This study, and previous work on DAMP (3, 14, 21), considers *E. coli* batch culture in which there is no renewal of media, meaning that peroxide detoxification is permanent. As media inflow and outflow increase in a system, the ability of individual cells to detoxify ROS is decreased (50), it therefore seems possible that, increasing flow will be similar to transitioning from model D (fixed supply of environmental ROS, resulting in DAMP, Fig. 2) to model C (fixed level of environmental ROS, resulting in no DAMP, Fig. 2). That would mean that the spatial structuring and resulting fluid dynamics of flow, which can be critical for bacterial competition (44, 51), are also critical for mutation supply. Such factors vary greatly among natural environments, meaning that the effect of DAMP could be very different in low versus high through-flow environments (e.g. soil rather than water or lung rather than bladder). Tracing mutagenesis in single cells of spatially structured populations (52) has the potential to define the spatial scales and through flow conditions under which benefits from mutagen degradation are shared.

Our finding that oxygen is key to mutation rate plasticity is supported by mutation accumulation experiments showing that increased oxygen uptake is correlated with increased mutation rates (4). However, existing literature is not agreed on this point – anaerobic fluctuation assay-like experiments also report reduced mutation frequencies for resistance to multiple antibiotics (53), while anaerobic mutation accumulation experiments instead report increased mutation rates (54, 55). Work assessing mutation rate by the accumulation of resistance mutants in chemostats also shows oxygen limitation to reduce mutation rates relative to carbon limitation (10). This discrepancy is likely due to the change in mutational spectra caused by anaerobiosis: although overall mutation rates increase, base pair substitutions (BPS) fall in frequency by 6.4 times (54) and it is such BPS which we modelled computationally and are often responsible for antibiotic resistance (56–58), particularly to rifampicin, the drug we used for our mutation rate estimates (57). In line with our finding that iron and oxygen disruption are similarly able to abolish DAMP, iron and oxygen limitation produce similar mutational spectra (10). The loss of DAMP in the Δ*fur* strain is perhaps due to higher intracellular iron levels producing a greater rate of H_2_O_2_ breakdown into DNA-damaging radicals before it can be detoxified, reflective of ODE model C in which a constant ROS burden is applied and no DAMP seen.

Mutation supply is a key evolutionary hurdle often limiting the adaptation of populations (1, 59–61). As mutation supply depends on population size, one might expect the supply of mutations, for instance to AMR, to be severely limited in small populations, such as the small number of cells forming an infectious propagule of *E. coli* (17). Even when population size is sufficient to enable adaptation, mutation supply may have more subtle effects on the course of evolution, as demonstrated by the pervasive effects of mutational biases (62–64). However, due to the action of DAMP in elevating mutation rates at low density, small populations can experience a very similar supply of mutations to large populations (as demonstrated in our data, Fig. S4). For *E. coli* at least, there is a limit to this effect as beyond intermediate densities (∼7×10^8^ CFU ml^-1^) the action of stress-induced mutagenesis causes mutation rates to rise, rather than fall, with increased density (14).

The dependence of DAMP on active cellular control of H_2_O_2_ concentrations, uncovered here, helps explain it’s highly conserved nature. The evolution of cellular systems in an anaerobic world for ∼1 billion years (65) means that all branches of life are similarly vulnerable to damage by ROS, leading to parallel effects of ROS damage across life (66), potentially including its population level control in DAMP. Although DAMP is highly conserved, it is notably not seen in *Pseudomonas aeruginosa* (3), despite this species being a close relative of *E. coli.* How DAMP is lost between close evolutionary relatives remains an interesting question and is perhaps linked to the formation of multicellular aggregates by *P. aeruginosa* (67). Population associations with mutation rate are widespread, including a significant negative relationship between the effective population size and mutation rates across vertebrates (68) as well as microbes (69). Such patterns seem likely to be driven by the increased efficiency of natural selection against the deleterious effects of mutation in large populations (the drift barrier hypothesis, (70)), rather than any adaptive benefit. The broad reach of such non-adaptive explanations and the fact that the evolutionary effects of DAMP are yet to be explored means that any adaptive explanations should be approached with great caution. Nonetheless, in strains with DAMP, the mutation rate decreases as the absolute fitness increases, providing the greatest mutation supply to the most poorly performing populations (21). Mutation supply also rises in the most nutrient rich environments (14), perhaps providing greater evolutionary potential where competition is most intense. Such plausible evolutionary benefits of DAMP could exist, even if the ultimate origins of its conserved mechanism lie not in selection for its indirect effects via mutation, but in the legacy, across domains of life, of the chemistry of the Great Oxidation Event (71).

## Materials & Methods

### Ordinary differential equations: Model A

All variables (Fig. 1A, Table. 1) are measured in molar concentration within the cytoplasm, aside from the volume of that cytoplasm (***cytVol***), measured in mL, and external glucose (***eGlc***) and number of growing cells (***wtCell*** & ***mCell***) which are measured as molar concentrations within the 1ml batch culture. It is possible to convert between cytoplasmic and total metabolite concentrations through scaling by the cytoplasmic volume; this is calculated as the number of cells multiplied by a volume of 1.03 × 10^-9^ μl per cell (72). The reaction of ***dGTP*** with ***ROS*** creates oxidised dGTP (***odGTP***) which is then incorporated into DNA, creating AT > CG base pair substitution mutations (20). Mutations caused by ***odGTP*** may be avoided or repaired by the action of MutT, MutY and MutS enzymes (73). By dividing the number of mutant cells (***mCell***) by the total cell number (***mCell*** + ***WtCell***) at any point during the simulation, a mutation rate (bp^-1^ generation^-1^) across the simulation up to that point, can be calculated.

The uptake of glucose is described by saturating Michaelis Menten kinetics whilst the oxidation of ***dGTP*** is described as a bimolecular reaction dependent on the cytoplasmic concentrations of ***dGTP*** and ***ROS***. All other steps are described by 1st order mass action kinetics in which the rate equals the concentration of the reactant multiplied by a rate constant (Eq. 1-10). The model is parameterised from published enzymatic and culture data alongside our own wet lab data (Table. 1).

R code to recreate all models and analysis is available as a supplementary file. All models were simulated in R (V4.3.1) (74) using package deSolve (V1.36) (75), logarithmic sequences were produced with emdbook (V1.3.13) (76), data handling and plotting was done using the tidyverse (V2.0.0) (77) and magrittr (V2.0.3) and parallel computing was done using parallel (V4.3.0), doParallel (V1.0.17) and foreach (V1.5.2). Linear mixed models were fitted to lab data with nlme (V3.1-162) (78), and plots formatted and coloured using cowplot (V1.1.1), gridExtra (V2.3), ggeffects (V1.3.1) (79) and RColorBrewer (V1.1-3). The flow diagram (Fig. 1A) was made using R package DiagrammeR (V1.0.10).

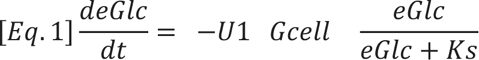

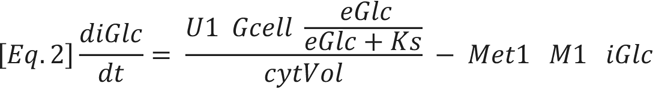

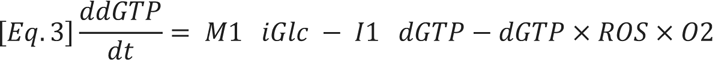

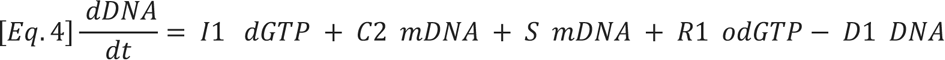

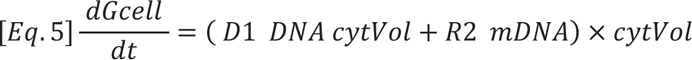

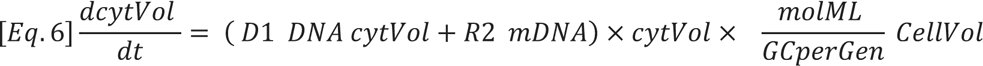

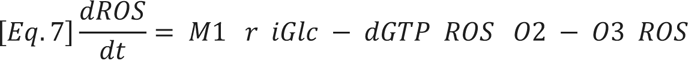

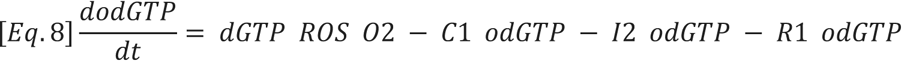

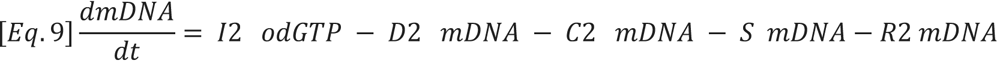

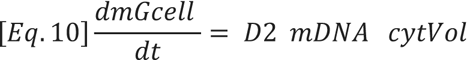

*Equations 1-10: ODE equations for initial model (A)*

### Model Variants

#### Model B - Glucose uptake increases at low eGlc

Original Michaelis Menten kinetics are removed as this reverses the intended effect.

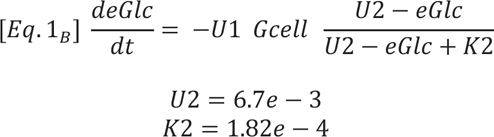

Both U2 and K2 are in Molar units.

6.7×10^-3^ is chosen as a value slightly higher than maximum eGlc so that the value of 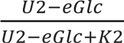 can cover almost a full range of 0 to 1. This means that glucose uptake rate will increase from almost 0 to 100% of the measured uptake rate as the external glucose concentration falls. K2 is given as 1.82×10^-4^ as this value produces the most negative DAMP slope achievable within the structure; values were tested from 1.82×10^-6^ to 1.82×10^-2^ (SI code file).

#### Model C - Constant ROS concentration uncoupled from metabolism

By decoupling ROS concentration from metabolism there is no extra production of ROS in cells grown to higher density, we expect this to prevent a positive DAMP slope.

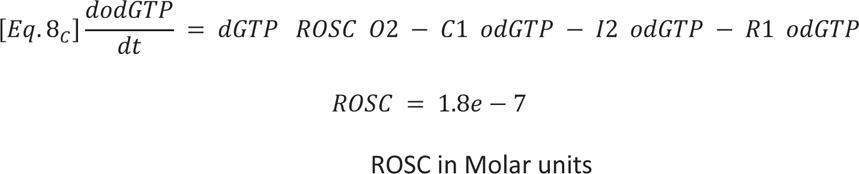

ROS is removed as a variable and replaced with a constant concentration of 1.8×10^-7^, this is within the known internal ROS concentration of 1.3×10^-7^ - 2.5×10^-7^ M (87) and produces mutation rate of 1.93×10^-10^ based on lab data and (20).

#### Model D - Constant ROS production regardless of population density

By creating a situation in which ROS is produced in the media at a constant rate (e.g. (22)) and then taken up among all present cells, higher density populations will expose each individual cell to less ROS. We expect this to create negative a DAMP slope as more mutagenic ROS exists inside the cells of low-density populations. This structure may be reflective of a situation in which iron is limited and so higher density populations have less iron per cell. Because of the role of iron in the Fenton reaction it may be expected that less iron leads to less ROS damage in the cells, as observed by (88).

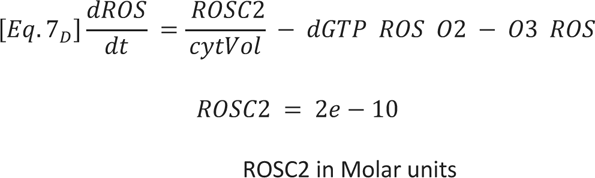

ROSC2 defines the number of millimoles of hydrogen peroxide produced in the system each second, this is split between the cytoplasm of all cells. The chosen value of 2×10^-10^ creates a H_2_O_2_ production rate at 76% of that expected from (24) and a mutation rate 96.8% of that expected from (20).

#### Model E - ROS removal dependent on internal glucose

We expect greater rates of ROS removal to lead to lower rates of GTP oxidation, and therefore lower mutation rates. If ROS is more able to be degraded when resources are abundant this may produce DAMP.

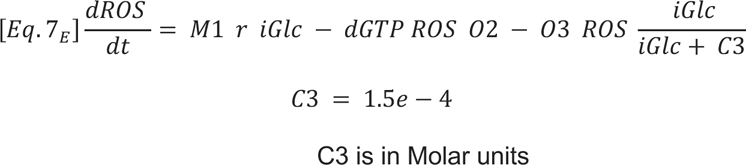

C3 is adjusted to produce known mutation rate of 1.98×10^-10^ base pair substitutions per nucleotide in 0.2% glucose minimal media (20).

#### Model F - ROS removal dependent on population density

We expect direct control of ROS degradation by population density to allow cells in higher density populations to avoid mutations more efficiently.

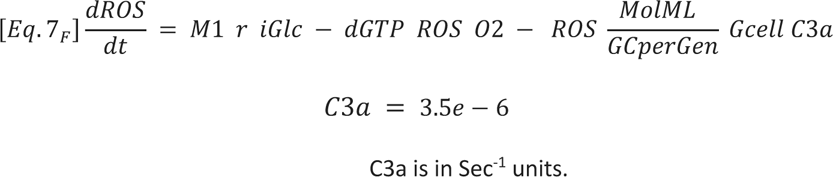

C3a of 3.5×10^-6^ is chosen to reproduce the mutation rate of 2.05×10^-10^ base pair substitutions per nucleotide.

#### Model G - MutT activity upregulated by internal glucose

MutT activity is known to be essential to DAMP and so density dependent MutT activity is a candidate DAMP mechanism. iGlc accumulates at higher levels in cells growing to high density, we expect high MutT activity in these cells to lead to a reduced mutation rate due to MutT cleaning of odGTP.

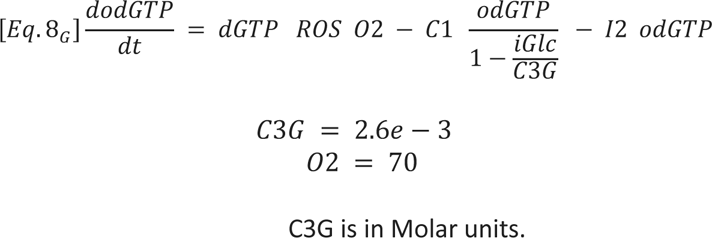

2.6×10^-3^ is selected as a number slightly higher than the maximum iGlc achieved (∼0.0023), this prevents MutT activity levels from falling below 0. O2 is refitted to 70 to restore desired mutation rate.

#### Model H - MutT activity upregulated by odGTP

If MutT activity is actively upregulated to degrade odGTP at a higher rate upon exposure to higher odGTP concentrations, then we expect cells grown in higher glucose, with higher internal metabolite concentrations, to have a greater ability to evade mutations caused by odGTP.

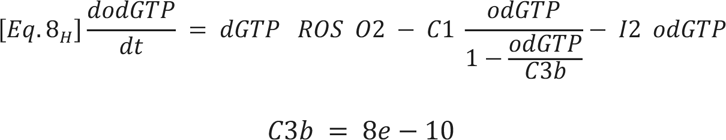

C3b is in Molar units.

8×10^-10^ is selected as a number slightly higher than the maximum odGTP achieved, this prevents MutT activity levels from falling below 0.

#### Model I - MutT activity upregulated by ROS

Reasoning and value selection as in Models G/H

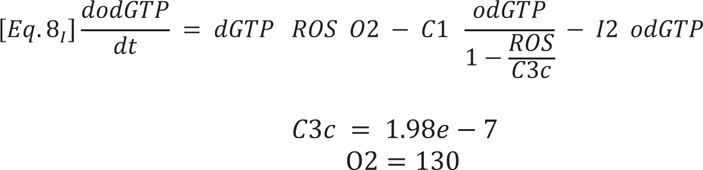

C3c is in Molar units.

#### Model J - Michaelis Menten MutT kinetics

Michaelis Menten kinetics describe saturating, enzyme catalysed reactions. In this situation reaction rates proceed slower at low substrate concentrations rising to an asymptote at maximum reaction rate. As with models G/H/I we expect this to reduce mutation rates by increasing MutT activity in high density populations with greater internal metabolite concentrations.

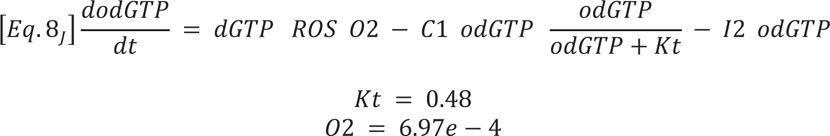

Kt is in Molar units.

Kt value given by (19), O2 is then titrated to restore mutation rate as in (20).

#### Model K – Separated activity of ahpCF and katEG genes + limited diffusion of ROS across the plasma membrane

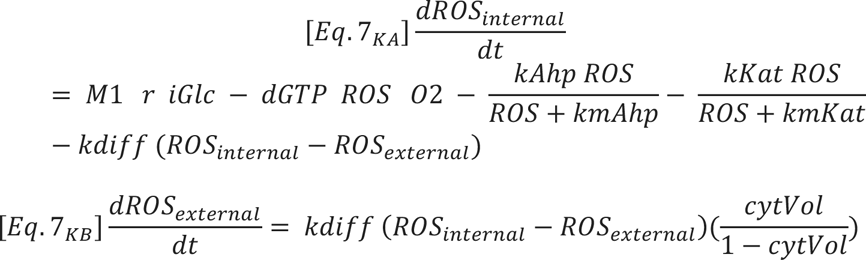

As in (23) the activity of alkylhyrdoperoxidase and catalase proteins are separated to allow for their specialisations to low and high H_2_O_2_ concentrations respectively. Michaelis Menten constants are as follows:

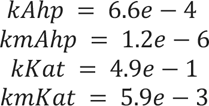

All in Molar units.

Permeability coefficient, diffusion coefficient and cell surface area are taken from (23) to calculate the diffusion coefficient as follows:

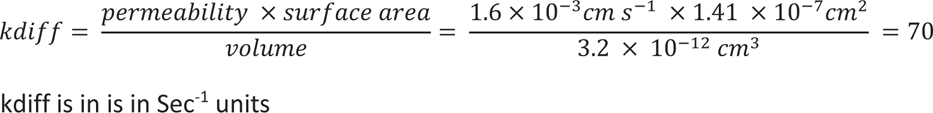

H2O2 production rate and standing concentration are restored to expected values by altering the value of r:

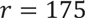

### Global Sensitivity Analysis

For each parameter within each model, 50,000 values between 10% to 1000% of the baseline value (Table. 2), spaced evenly along a log scale were tested. The set of values for each individual parameter were then independently shuffled so that no parameters were corelated with one another; allowing for substantial exploration of the available parameter space. Of these 50,000 parameter sets some encountered fatal errors in the ODE solver and so did not produce a DAMP slope estimate, the number of parameter sets run without fatal error is shown in Table 3 as ‘complete’. Results were filtered for the following criteria: 1) Stationary phase is reached in all glucose conditions (defined as an average increase of less than 1 cell per 10 second time step across the last 1000 time steps (2.7 hours) of the simulation), 2) final population size >1×10^7^ & <1×10^10^ at every glucose condition, 3) final population size increases with each increase in glucose concentration, 4) mutation rate >2×10^-12^ & <2×10^-8^ at all glucose conditions and 5) log-log relationship between mutation rate and final population size is substantially linear (defined by R-squared > 0.5). After this filtering the following number of parameter sets were retained for each model (Table 3).

**Table 3:** Counts of completed and filtered simulations from 50,000 parameter sets produced for global sensitivity analysis for each model variant. ‘Complete’ column lists the number of these parameter sets that were able to be simulated without fatal error from the ODE solver. ‘Filtered’ column lists how many parameter sets remained after filtering as described above.

| Model | Complete | Filtered |
| --- | --- | --- |
| A | 49971 | 12261 |
| **B** | 33274 | 5444 |
| **C** | 49988 | 13496 |
| **D** | 49990 | 7128 |
| **E** | 49555 | 10267 |
| **F** | 49982 | 10115 |
| **G** | 23311 | 2730 |
| **H** | 49900 | 2210 |
| **I** | 22832 | 3334 |
| **J** | 49994 | 11741 |
| **K** | 45558 | 7365 |

### Strains used in this study

The parent of the Keio collection is *E. coli* strain BW25113 (F-, Δ(*araD*-araB)567, Δ*lacZ*4787 (::*rrnB*-3), λ-, *rph*-1, Δ(*rhaD*-*rhaB*)568, *hsdR*514). *E. coli* hpx^-^ LC106 mutant is Δ*ahpCF*’ kan:: Δ*ahpF* Δ (*katG*17::Tn10)1 Δ (*katE*12::Tn10)1 (26). *E. coli* single gene knockouts Δ*fur*, Δ*feoB*, Δ*tonB* and Δ*ahpF* are sourced from the Keio collection (89). *E. coli* K-12 strain MG1655 is from Karina B. Xavier. Nalidixic acid resistant strains hpx^-^ (*gyrA* D87Y) & hpx^-^ (*gyrA* D87G) were isolated from independent fluctuation assays of the original hpx^-^ strains on 30 mg L^-1^ nalidixic acid selective plating.

Strains Δ*fur*, Δ*feoB*, Δ*tonB*, Δ*ahpF*, hpx^-^, hpx^-^_nalR_(D87Y) and hpx^-^_nalR_(D87G) were sequenced to 30x depth by MicrobesNG to verify gene deletions. Lack of KatE activity in hpx^-^ was verified by covering a colony on TA agar with 30% H_2_O_2_ with no bubbles of oxygen observed (as in (25)), the MG1655 wild-type was used as a positive control. Mutations were identified using breseq version 0.36.0 (90, 91) with bowtie2 version 2.4.1 and R version 4.2.0 and are listed in Table. S2. For hpx^-^ strains the reference genome used was the *E. coli* K-12 MG1655 genome [(92), NCBI accession U00096.3]. For Keio knockout strains the reference genome used was the *E. coli* K-12 BW25113 genome [(93), NCBI accession CP009273.1], with additional annotations for insertion (IS) element regions to improve the calling of mutations related to IS insertion (modified Genbank format file as file S1 in (1)).

### Media

We used Milli-Q water for all media, all chemicals are supplied by Sigma-Aldrich unless stated otherwise. LB medium contained: 10 g of NaCl (Thermo Fisher Scientific), 5 g of yeast extract (Thermo Fisher Scientific) and 10 g of tryptone (Thermo Fisher Scientific) per litre. DM medium contained 0.5 g of C_6_H_5_Na_3_O_7_ ·2H_2_O, 1 g of (NH_4_)2SO_4_ (Thermo Fisher Scientific), 2 g of H_2_KO_4_P and 7 g of HK_2_O_4_P· 3H_2_O per litre; 100 mg L^-1^ MgSO_4_ ·7H_2_O (406 µmol) and 4.4 μg L^-1^ thiamine hydrochloride were added to DM after autoclaving. Selective tetrazolium arabinose agar (TA) medium contained 10 g of tryptone, 1 g of yeast extract, 3.75 g of NaCl and 15 g bacto agar per litre; after autoclaving 3 g of arabinose and 0.05 g of 2,3,5-triphenyl-tetrazolium chloride were added per litre, this was supplemented with freshly prepared rifampicin (50 µg ml^−1^) or nalidixic acid (30 µg ml^−1^) dissolved in 1mL of methanol or 1M NaOH respectively when required. For all cell dilutions sterile saline (8.5 g L^-1^ NaCl) was used.

### Fluctuation Assays

Fluctuation tests were conducted as described in (94). Briefly; initial growth of glycerol stocks in LB was carried out for 4 hrs for all strains aside from hpx^-^ which was grown for 7 hrs due to its reduced growth rate. A dilution factor of 1000x was then used for transfer to overnight cultures. Overnight acclimatisation was carried out in DM supplemented with 3.5% LB or 250mg L^-1^ glucose with nutrient type matching that of the fluctuation assay. The density achieved in the assay was manipulated by growth in varying nutrient conditions, either 2 - 5% LB diluted in DM or 80 - 1000 mg glucose L^-1^. Selective plates were prepared 48 hours before use and stored for 24 hours at room temperature followed by 24 hours at 4०C. All strains were plated on rifampicin selective media.

Anaerobic conditions were produced by incubating the 96 deep well plates in an airtight 2.6L container with one Anaerogen 2.5L sachet (Thermo Scientific). The Anaerogen sachet rapidly absorbs oxygen and releases CO_2_ creating anaerobic conditions. Aerobic plates of matching design were grown in an identical container ventilated with 8 × 4mm diameter holes without Anaerogen sachet. In these plates 2-4 wells in each 96 well plate contained DM supplemented with 2.5% LB, resazurin and *E. coli* MG1655, leaving space for fluctuation assays of 15-16 parallel cultures. On removing the 96 well plates from incubation the resazurin absorbance at 670 nm was measured; this quantifies the change from pink resorufin (aerobic cell growth) to clear dihydro resorufin (anaerobic cell growth) thus providing an objective measure of anaerobiosis (Fig. S12).

During coculture fluctuation assays between BW25113 wild-type and hpx^-^ both strains were grown up in LB, diluted LB overnight and diluted into cultures of ∼1×10^3^ CFU mL^-1^ as above. Some combination of these 2 initial cultures was then mixed in each parallel culture ranging from an hpx^-^ :wild-type ratio of 1:1 to 124:1 (recorded in the supplementary data file as ‘Mut_to_WT_ratio’). Plating of these cultures on TA or TA+Rif agar enabled the Ara+ (white) hpx^-^ colonies and the Ara-(red) wind-type colonies to be distinguished. For assays using NalR hpx^-^ strains, selective plating was done on TA+Rif+Nal plates and so only hpx^-^ mutants and not wild-type mutants were counted, Nt was determined for both strains using plating on both TA+Nal and TA. Due to amino acid synthesis defects, hpx^-^ cells cannot be cultured in glucose minimal media and so all cocultures were conducted in dilute LB media (30).

In order to test for any differences in survival of hpx^-^ (NalR) grown in monoculture VS with differing densities of BW25113 wild-type cells we conducted a reconstruction test (Fig. S9). A known quantity of hpx^-^ (NalR + RifR) cells were plated with one of the following treatments: wild-type supernatant (3.5%LB overnight growth), sterile DM, 80U mL^-1^ catalase, hpx^-^ (5%LB overnight growth), wild-type (2.5%LB overnight growth), wild-type (3.5%LB overnight growth), wild-type (5% overnight growth). Raw data from the reconstruction test is available in the supplementary R code file.

Though fluctuation assays allow for high-throughput and low-cost estimates of mutation rate, they classically come with some important assumptions to consider (95). For example, the assumption that resistance markers will be selectively neutral is not reasonable in practice (42). Fortunately, this can be accounted for with the estimation of fitness cost which can then be accounted for in the estimation of mutational events using R package flan (V0.9). We find estimations of DAMP with the co-estimation of the individual fitness cost in each assay or with the application of the median mutant:wild-type fitness ratio estimation (median = 0.57, Fig. S3) to have no effect on our conclusions. In regression 4 (SI), re-running the model using either of these approaches to account for genotypes having different competitive fitness makes no difference to whether DAMP is inferred (i.e. categorising each treatment as DAMP, no DAMP or reverse DAMP, Table S1) for all treatments. In this study we allow flan to co-estimate fitness along with mutational events (*m*) for each assay. Occasionally this model failed to converge on estimates, in these cases average fitness effects were estimated from a model fitted to all successful estimates (Regression 3 (SI)) and then used to estimate *m* from the data with this pre-determined fitness effect of mutation. It is also possible to avoid issues of mutant fitness effects by using the *p*_0_ method of estimation (95, 96) in which parallel cultures are simply divided into those with or without any viable mutants. However, this method is more restrictive as only assays in which parallel cultures both with and without growth have been observed can the method be applied, it is also subject to more error on estimates than maximum-likelihood methods. Reanalysing our data with the *p*_0_ method shows DAMP to exist in the same set of treatments as in the original analysis (Fig. S11), discounting any effect of mutant fitness costs on our conclusions. Another potentially unrealistic assumption of the fluctuation assay is that there will be no death, this too is possible to account for using the tools provided in ‘flan’. We find that death rates as high as 10%, well beyond what would be expected under our conditions which lack added stressors, cause no changes in DAMP category (Fig. S13).

### Hydrogen Peroxide Measurement

External hydrogen peroxide is measured using the Amplex UltraRed (AUR)/Peroxidase assay as described in (22). All reagents were dissolved in 50 mM dibasic potassium phosphate. Diethylenetetraaminepentaacetic acid (DTPA) and AUR solutions were corrected to pH 7.8 with HCl or NaOH. Reactions containing 660 μL 1mM DTPA, 80 uL filter sterilised sample solution and 20 μL 0.25 mM AUR were mixed by vortexing before transferring 141 uL to three wells of a clear bottomed black 96-well plate. Fluorescence was measured at 580 nm excitation, 610 nm emission before and after the injection of 7.5 µL horseradish peroxidase (0.25 mg mL^-1^) to each well, net fluorescence was calculated as initial fluorescence subtracted from final fluorescence. H_2_O_2_ concentration was estimated by calibration to standard solutions of 5 & 20 μM H_2_O_2_ (Regression 5 (SI)). Because of background levels of fluorescence, some predicted concentrations were negative, this was accounted for by taking the absolute value of the lowest prediction and adding this to all predictions. The range of H_2_O_2_ concentrations we observed is in good agreement with similar measurements in the literature (e.g. Fig. 6B in (22)).

### Statistical Analysis

All statistical analysis was executed in R (V4.3.1) (74) using the nlme (V3.1-162) package for linear mixed effects modelling (78). This enabled the inclusion within the same regression of experimental factors (fixed effects), blocking effects (random effects) and factors affecting variance (giving heteroscedasticity). R package car (V3.1.2) (97) was used to carry out Chi-squared tests comparing slope to a null-hypothesis of 1 (Table. S1). In all cases log_2_ mutation rates were used. Details of all regression models are given supplementary statistical methods along with diagnostic plots and ANOVA tables for each model. The code and data to reproduce the main text figures are given in the accompanying R script, and supplementary data files, respectively. The content of the supplementary data files is explained in Table. S3. Standard deviation on estimates of *m* is calculated as in (98). The same R packages were used for parallel computing, data handling and plotting as for the ODE modelling, with the addition of plyr (V1.8.8), ggbeeswarm (V0.7.2) and gridExtra (V2.3).

## Supporting information

Supplementary Statistics

## Acknowledgements

Thanks to Danna Gifford for many insightful discussions during this study and for the improved BW25113 reference genome. Thanks also to The University of Manchester Research IT for their assistance and the use of the Computational Shared Facility. Thanks to MicrobesNG (http://www.microbesng.com) for their genome sequencing services. Thanks to Simon C. Andrews for providing the hpx^-^ strain and to Karina B. Xavier for providing the MG1655 wild-type strain.

## Supporting Information

**Figure S1.**
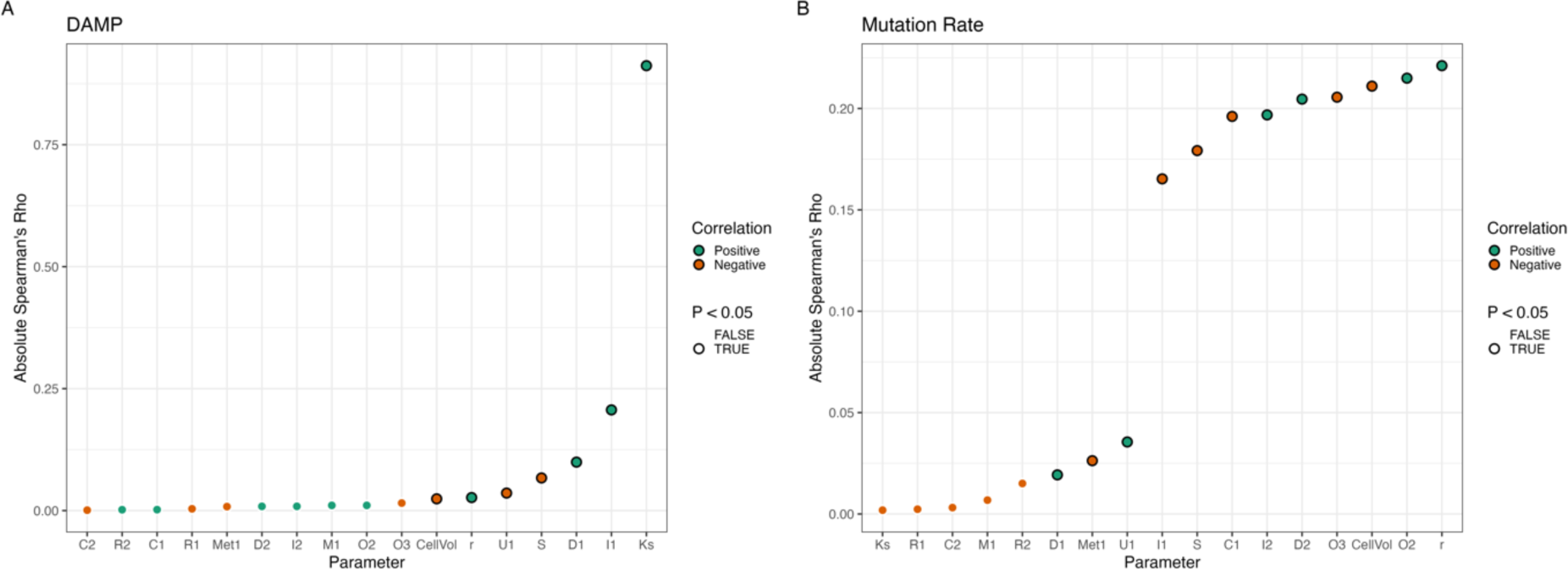
Global Sensitivity Analysis of Model A: Left Hand Side (A) shows the rank correlation, as quantified by Spearman’s Rank Correlation Coefficient, between each parameter and the slope of DAMP, parameters are ordered from least to most correlation from left to right. Right Hand Side (B) shows the equivalent information for the correlation between parameter values and mutation rate (at 250 mg L_-1_). Positive correlations are shown in green whilst negatively correlated parameters are shown in orange. Black borders show significant rank correlation (P < 0.05). Note the different y axis limits and × axis order on the left VS right hand side.

**Figure S2.**
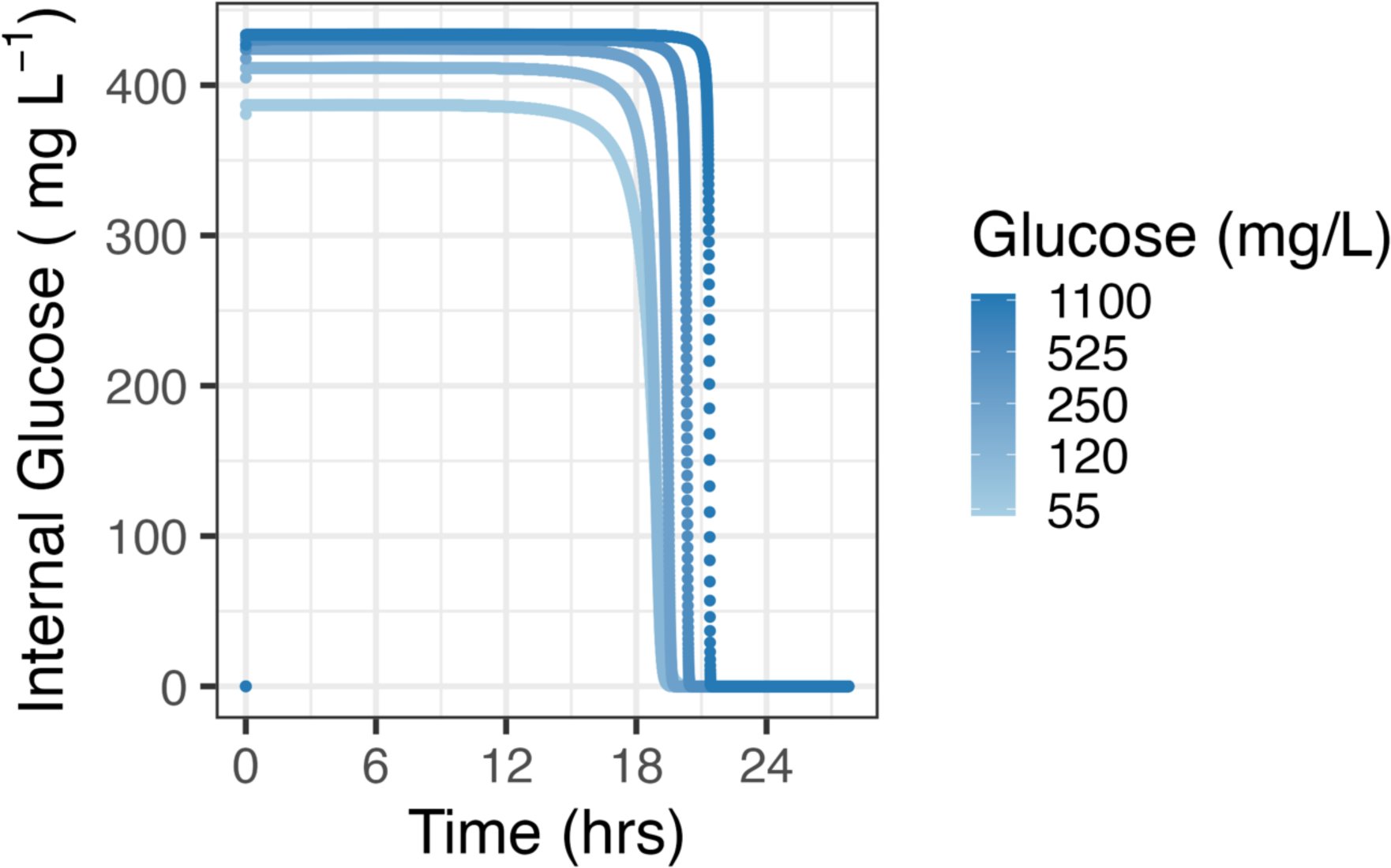
Dynamics of internal glucose over time in model A simulated at 5 log-spaced glucose concentrations from 55 to 1100 mg L_-1_. Higher levels of initial external glucose provision (point colour) lead to higher levels of internal glucose (y-axis).

**Figure S3.**
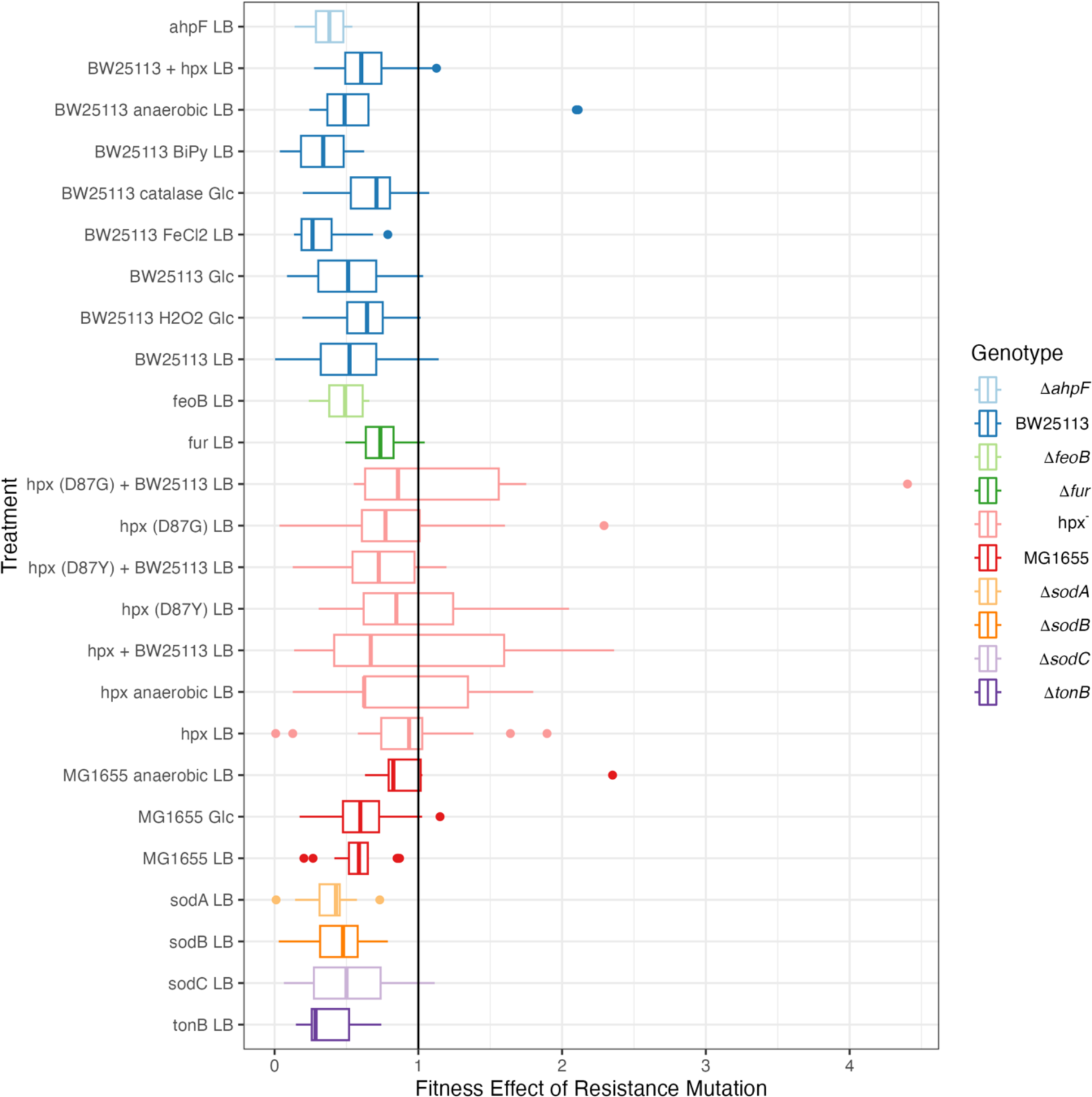
Fitness effects of resistance mutations where fitness is coestimated with mutational events. Boxplots shown for each treatment with colour representing genotype. Vertical lines inside boxes represent the median for that treatment, with the boxes depicting the interquartile range. The black vertical line at a fitness effect size of 1 represents neutral fitness effects.

**Figure S4.**
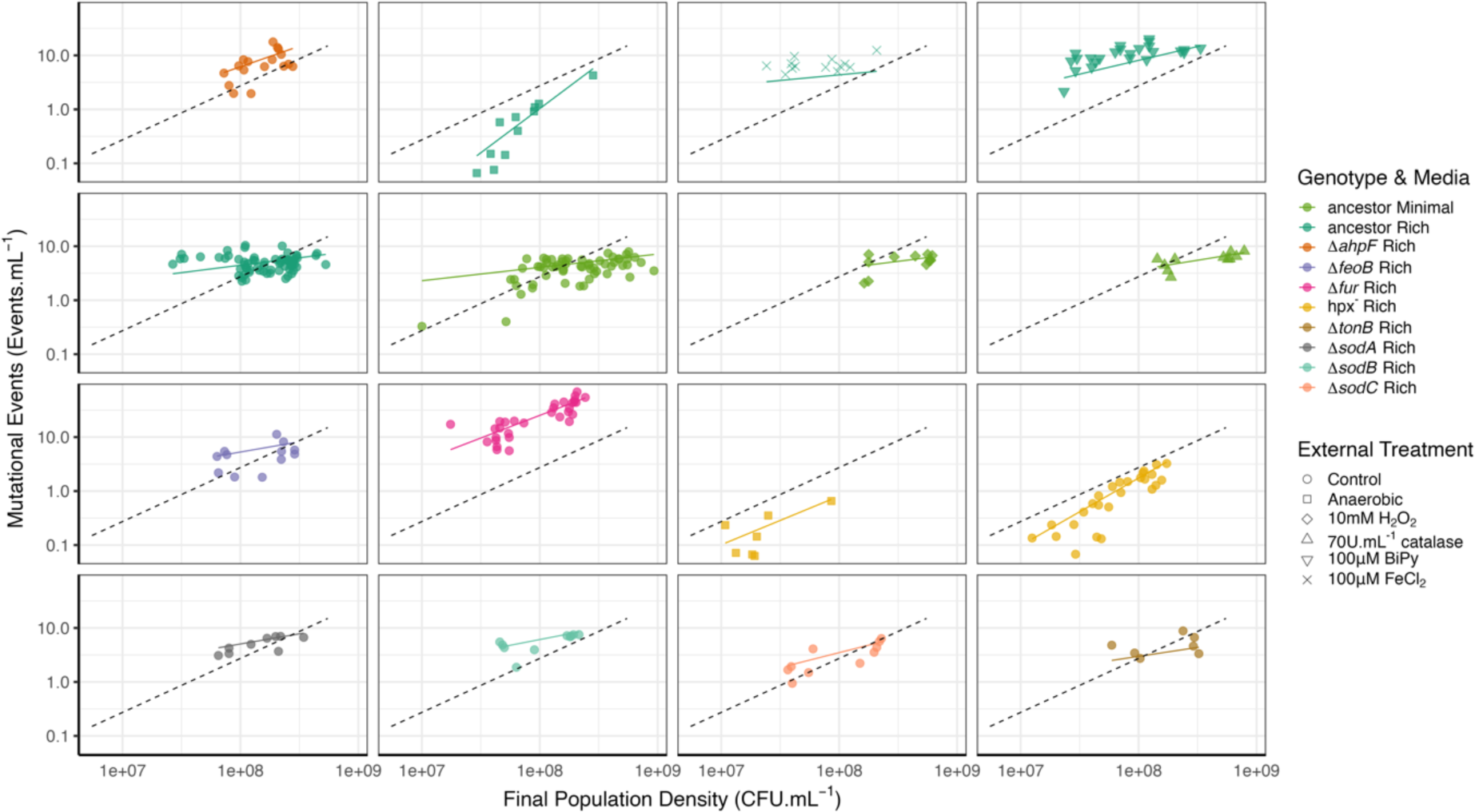
Raw data used in figure 3: Final population density is plotted against mutational events per mL on a log-log scale. Dashed lines show the null expectation of a constant mutation rate (i.e. slope=1) y intercept for dashed line is arbitrary. Coloured lines are fitted slopes from mod3 (Supplementary Statistics file), line gradients with 95% CI shown in figure 3. Treatments shown are BW25113 ancestor (1106 parallel cultures (pc) across 69 fluctuation assays (fa)); ancestor minimal media (942 pc, 59 fa); ΔahpF (266 pc, 17 fa); hpx_-_ (402 pc, 26 fa); ancestor anaerobic (168 pc, 11 fa); ancestor 10mM H_2_O_2_ (179 pc, 12 fa); ancestor 70U mL^-1^ catalase (167 pc, 11 fa); hpx^-^ anaerobic (105 pc, 7 fa); ancestor + chelator 2,2,Bipyridyl 100µM (382 pc, 24 fa); ancestor + FeCl_2_ 100µM (210 pc, 13 fa); ΔfeoB (192 pc, 12 fa); Δfur (504 pc, 31 fa); ΔtonB (113 pc, 7 fa).

**Figure S5.**
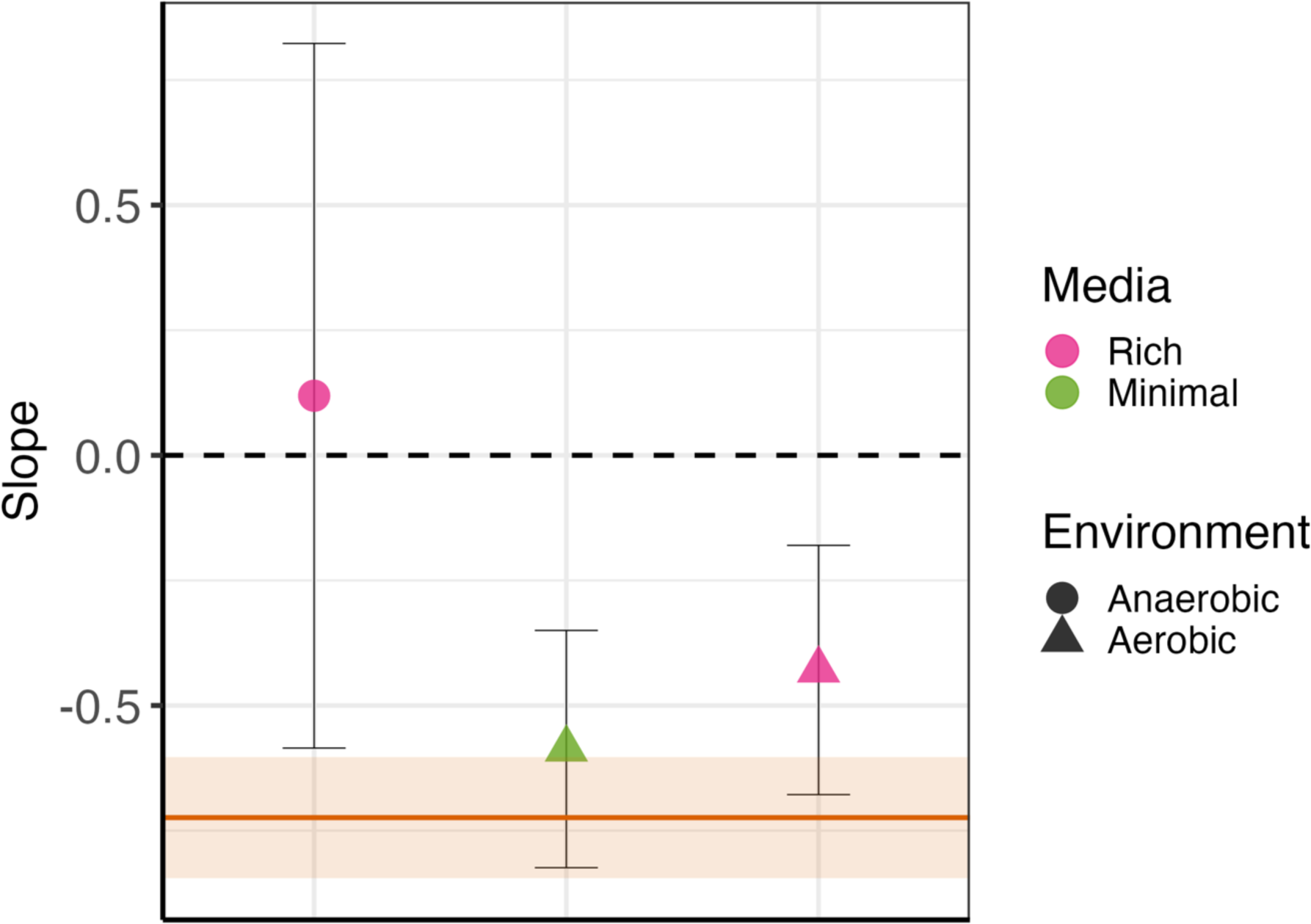
log-log relationship between population density (CFU mL_-1_) and mutational events (mL_-1_) in wild-type strain MG1655 under aerobic and anaerobic conditions. Pink circle = MG1655 rich media anaerobic (173 pc, 11 fa); Green triangle = MG1655 minimal media aerobic (273 pc, 17 fa); Pink Triangle = MG1655 rich media aerobic (285 pc, 18 fa). Orange line and shaded area shows DAMP for BW25113 in rich media as in Fig. 3 with 95% CI.

**Figure S6.**
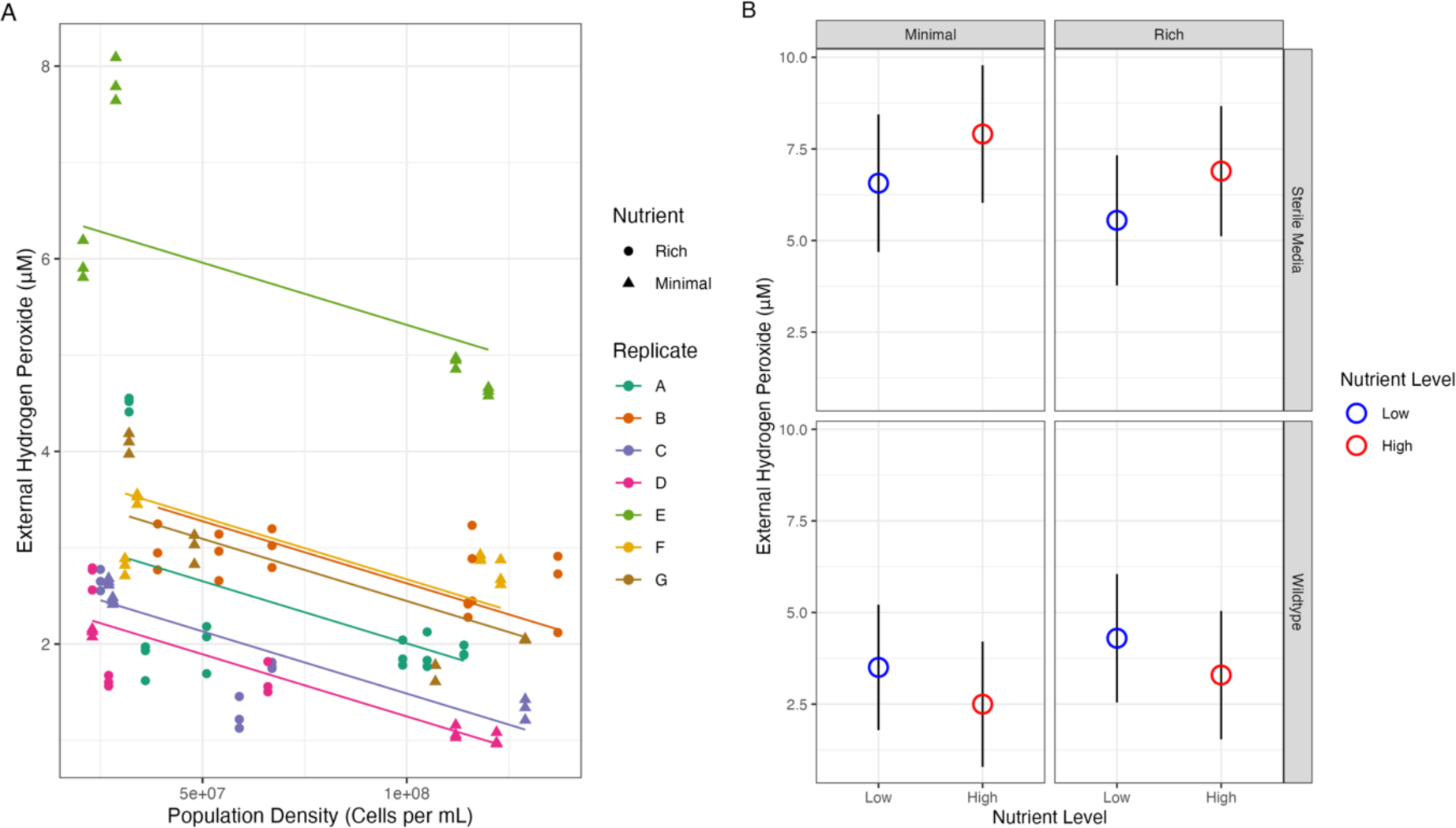
Effects of Population Density and Nutrient Level on H_2_O_2_. Left Hand Side (A) shows the relationship between population density and external H_2_O_2_ in cultures of MG1655 after 24 hours of incubation. Rich media is 2/5% LB diluted in DM, minimal media is 80/1000 mg L_-1_ glucose in DM. Population density is estimated from the optical density given the assumption that OD 1 = 1 × 10^9^ cells mL^-1^. Lines are from regression 7B (SI). Right Hand Side (B) shows the H_2_O_2_ concentration after 24 hours incubation in rich or minimal media; sterile or with wild-type MG1655, Regression 6 (SI); error bars show 95% CI. The interaction effect between nutrient level (Low versus High) and presence of a culture (Sterile Media versus Wild-type), where external peroxide decreases with nutrients increases in the presence of a culture but increases without one, is highly significant (F _DF=46_ = 9.8, P = 3×10^-3^, Regression 6 (SI)).

**Figure S7.**
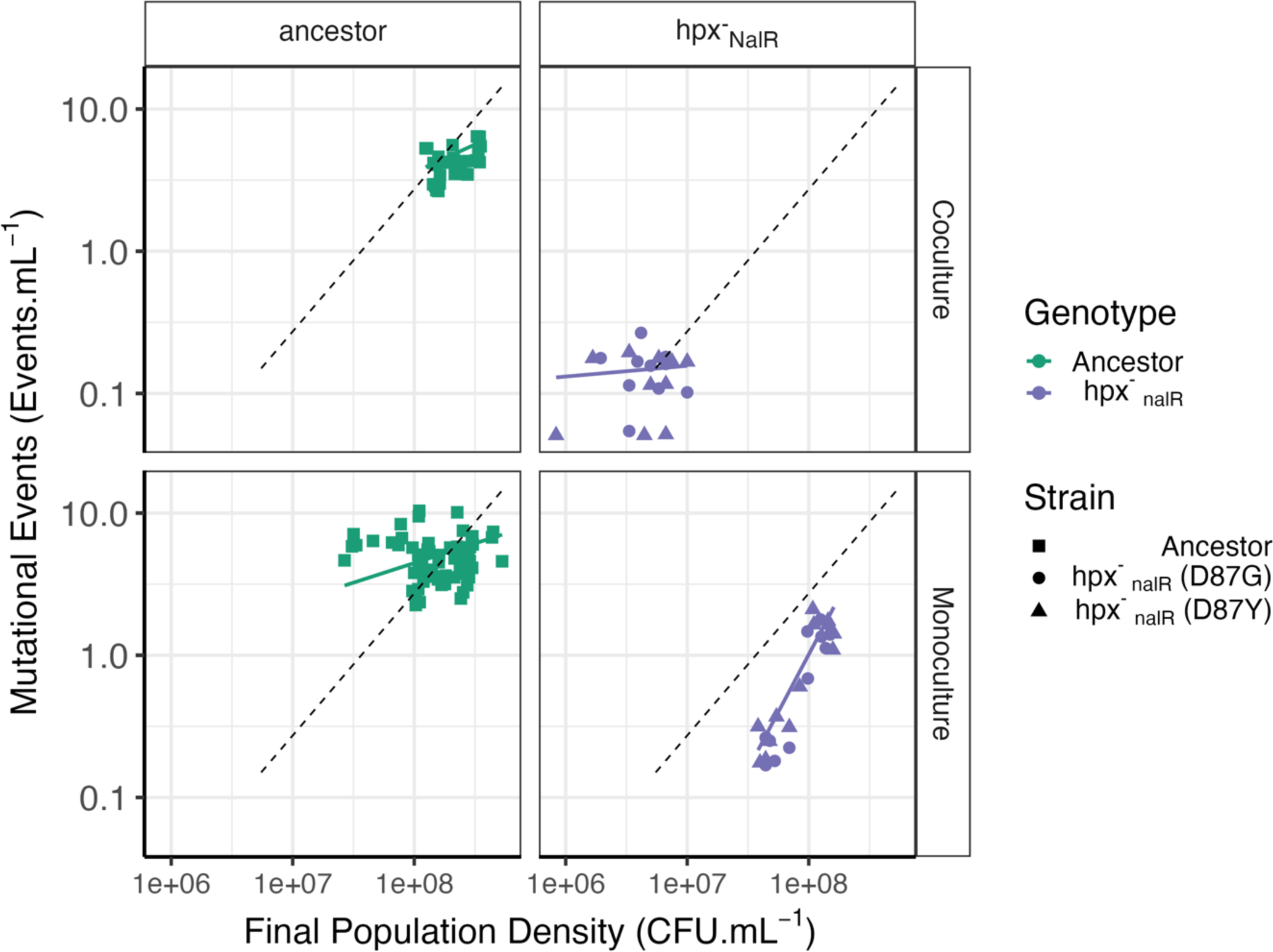
Raw data used in figure 4: Final population density of the focal strain is plotted against mutational events per mL on a log-log scale. Dashed lines show the null expectation of a constant mutation rate with a slope of 1. Ancestor coculture measurements are taken in coculture with hpx-, hpx D87Y & D87G are cocultured with ancestor BW25113. Lines are fitted slopes shown in fig. 4. BW25113 ancestor (1106 pc, 69 fa); BW25113 in coculture with hpx_-_ (498 pc, 31 fa); hpx^-^_nalR_ (388 pc, 24 fa); hpx^-^_nalR_ in coculture with BW25113 (319 pc, 20 fa).

**Figure S8.**
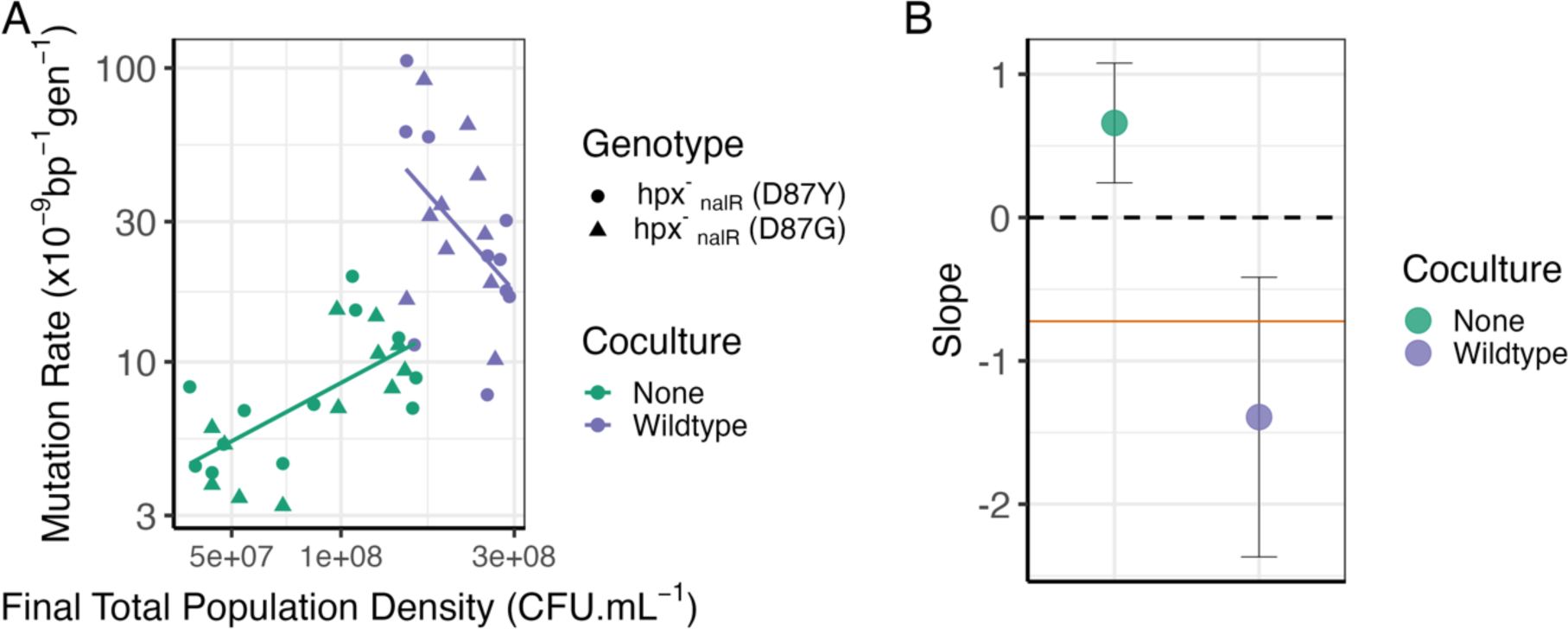
Relationship between total population density and mutation rate in hpx-with cocultured wild-type BW25113. A: Final population density (focal + coculture strain where relevant) is plotted against mutation rate on a log-log scale. hpx^-^_nalR_ monoculture (388 pc, 24 fa); hpx^-^_nalR_ in coculture with BW25113 (319 pc, 20 fa). Lines are fitted slopes shown from Regression 8 (SI). B: Slope and 95% CI on the lines shown in LHS graph. Horizontal orange line shows the slope of the BW25113 ancestor in rich media (Regression 4 (SI), Fig. 3). In monoculture hpx_-_ mutation rates increase with total population density whilst in coculture the wild-type restores a negative association between density and mutation rates (DAMP).

**Figure S9.**
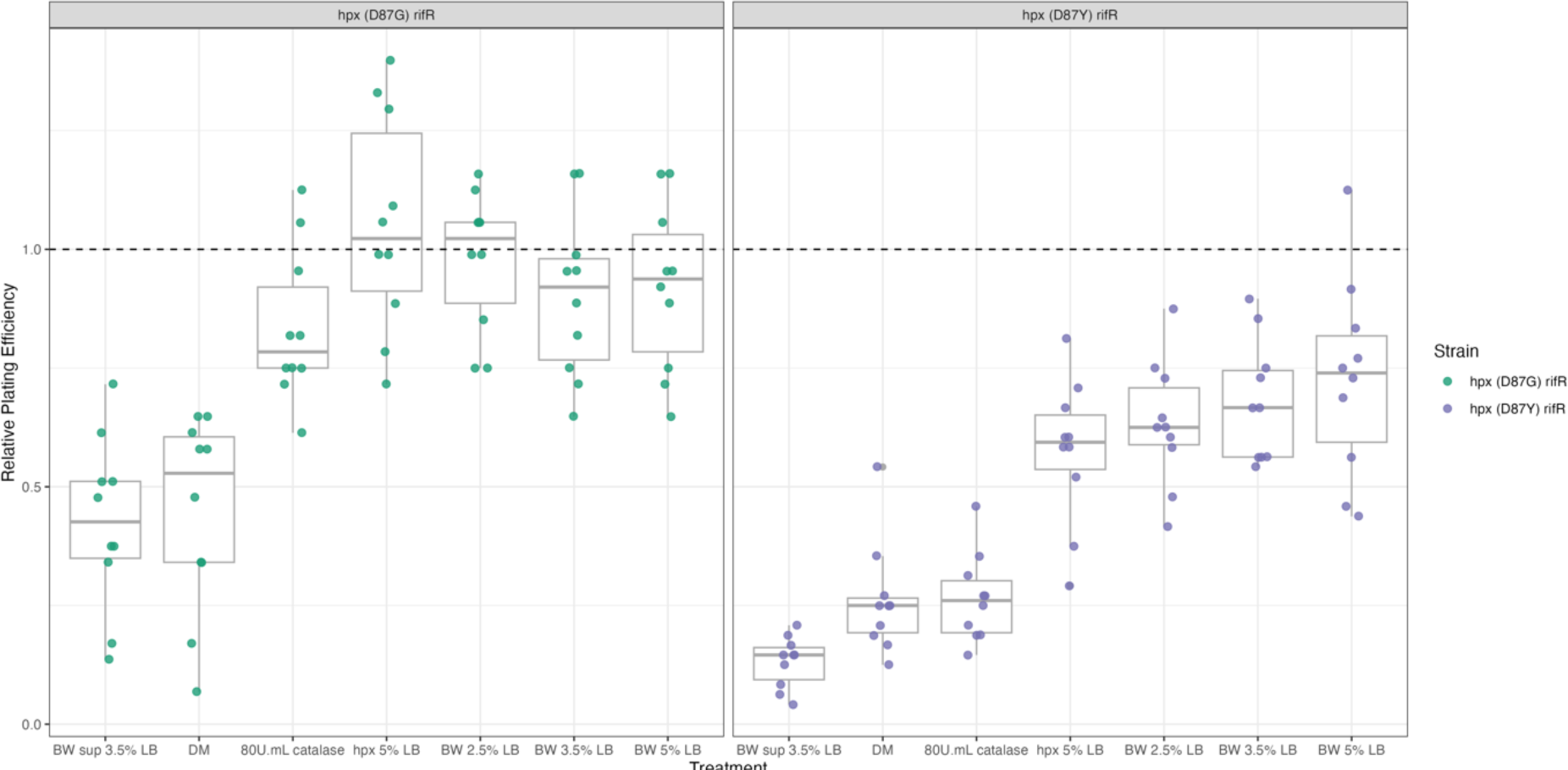
Reconstruction test showing the plating efficiency of rifampicin resistant hpx_-_ (D87Y) and hpx_-_ (D87G) when combined and plated in 1.25mL with: supernatant from mid-density BW25113, sterile DM media, DM media with 800U mL_-1_ catalase, hpx_-_ low-density, BW25113 low-density, BW25113 mid-density, BW25113 high-density. Plating efficiency is calculated as the number of colonies counted divided by the number of colonies counted on non-selective TA agar plates without any additional treatment.

**Figure S10.**
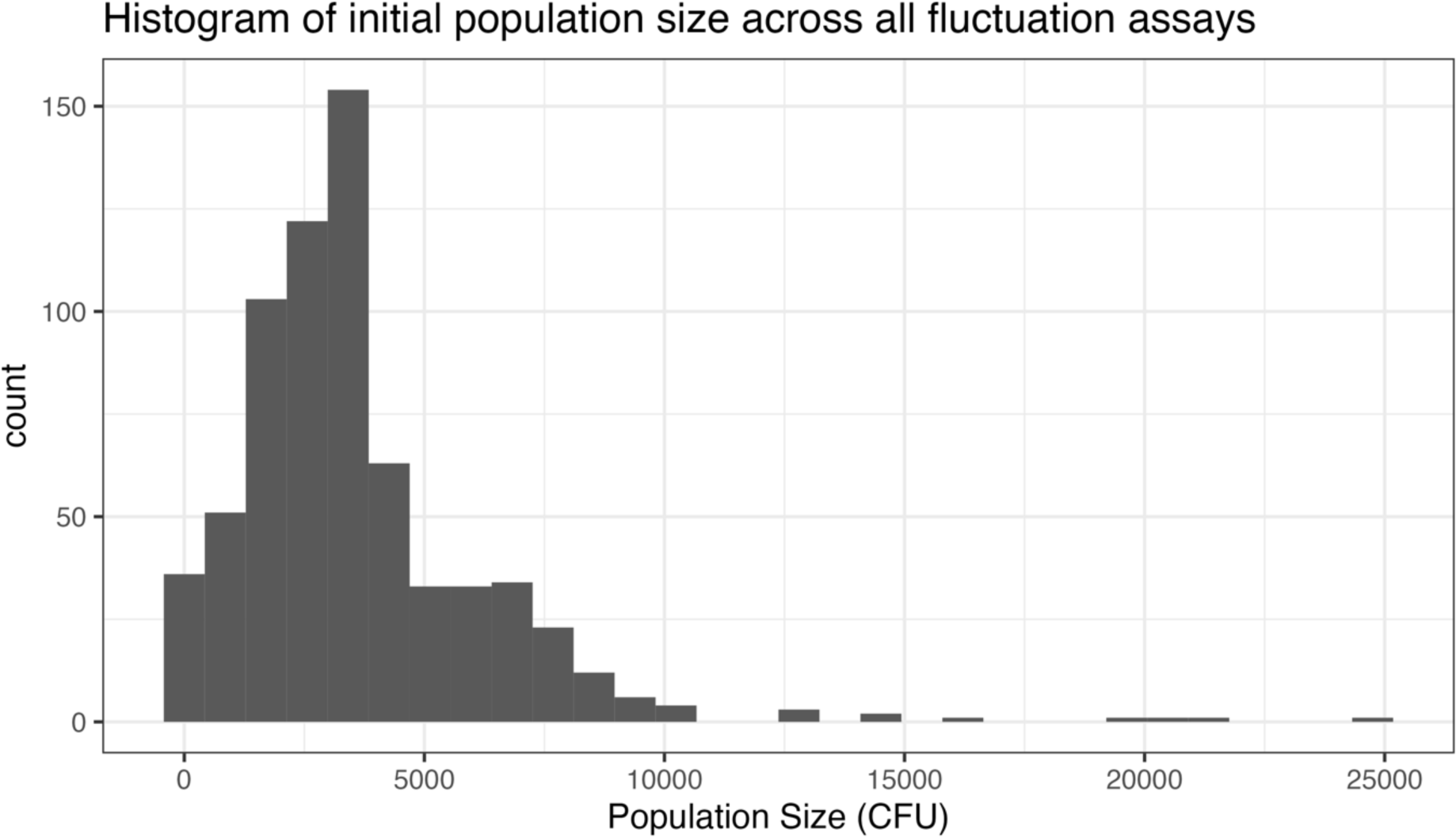
Distribution of initial population size across all fluctuation assays. Mean = 3397, median = 3000. Low population size is desirable in order to reduce the chances of resistant mutants being present in the starting population (‘jackpot cultures’).

**Figure S11.**
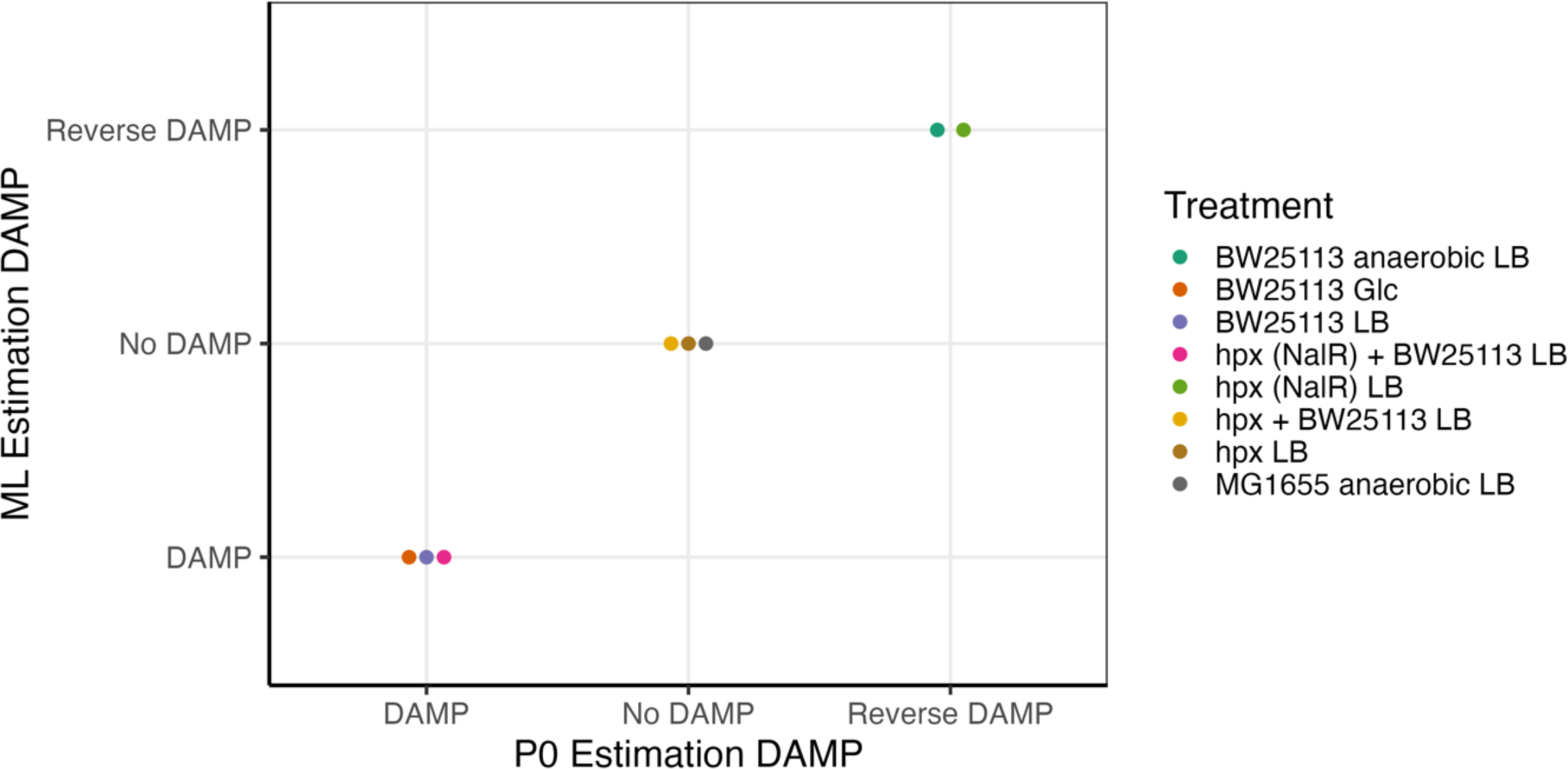
DAMP is seen in the same set of treatments using Maximum Likelihood or p_0_ estimation methods. Treatments in which 8< fluctuation assays can be analysed by the p_0_ method are shown. Colour indicates treatment identity. Treatments without DAMP in both methods (No DAMP:No DAMP) are: hpx_-_ + BW25113, hpx_-_ and MG1655 anaerobic. Treatments with DAMP in both methods (DAMP:DAMP) are: BW25113 glucose, BW25113 LB and hpx^-^_nalR_ + BW25113. Treatments with reverse DAMP (a significantly positive association between mutation rate and population density; Reverse DAMP:Reverse DAMP) are BW25113 anaerobic LB and hpx^-^_nalR_ LB. No treatments show a change in conclusions between methods.

**Figure S12.**
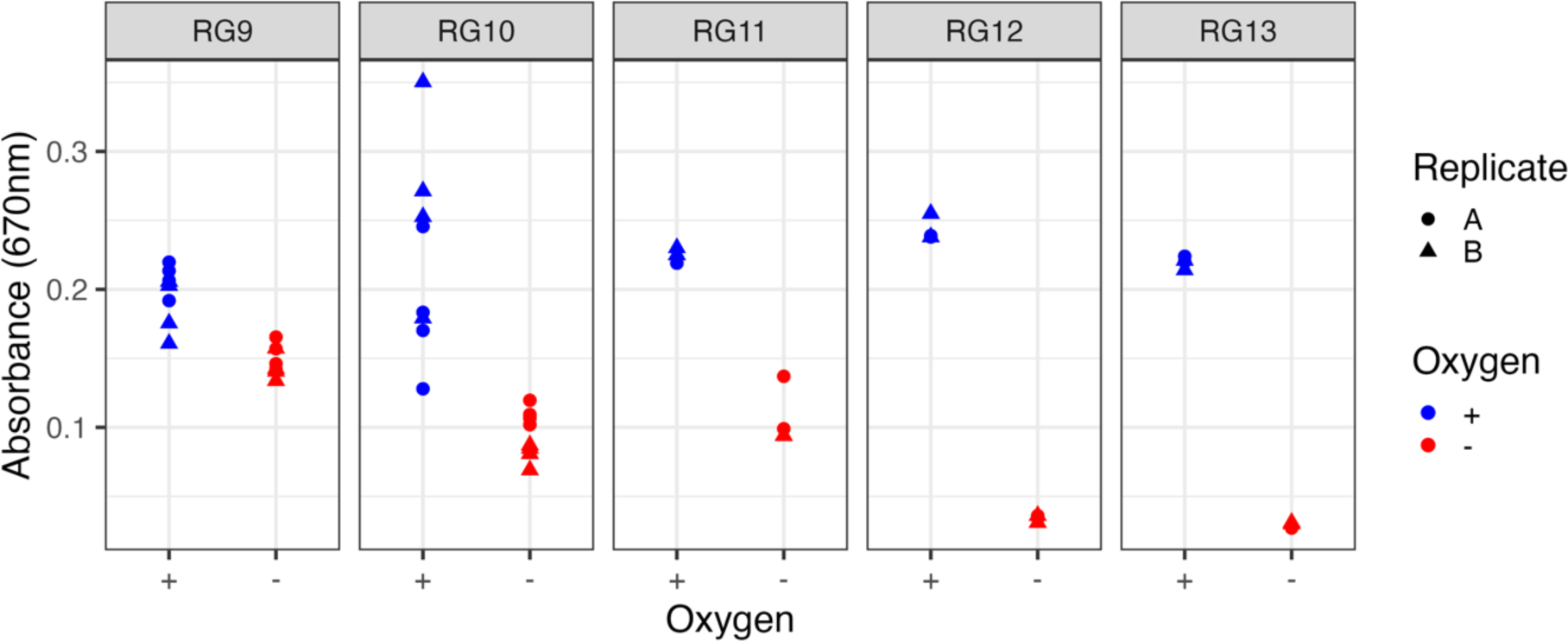
Reduction of resorufin to dihydro resorufin by anaerobic respiration. results in reductions in absorbance at 670nm verifying the anaerobic conditions during anaerobic fluctuation assays. Each of 5 blocks is shown as a separate facet; within each block 2 sets of paired fluctuation assays (A&B) were conducted in aerobic and anaerobic conditions, for each of these sets 2-4 measurements of resorufin/dihydro resorufin absorbance were taken after 24 hours of growth.

**Figure S13.**
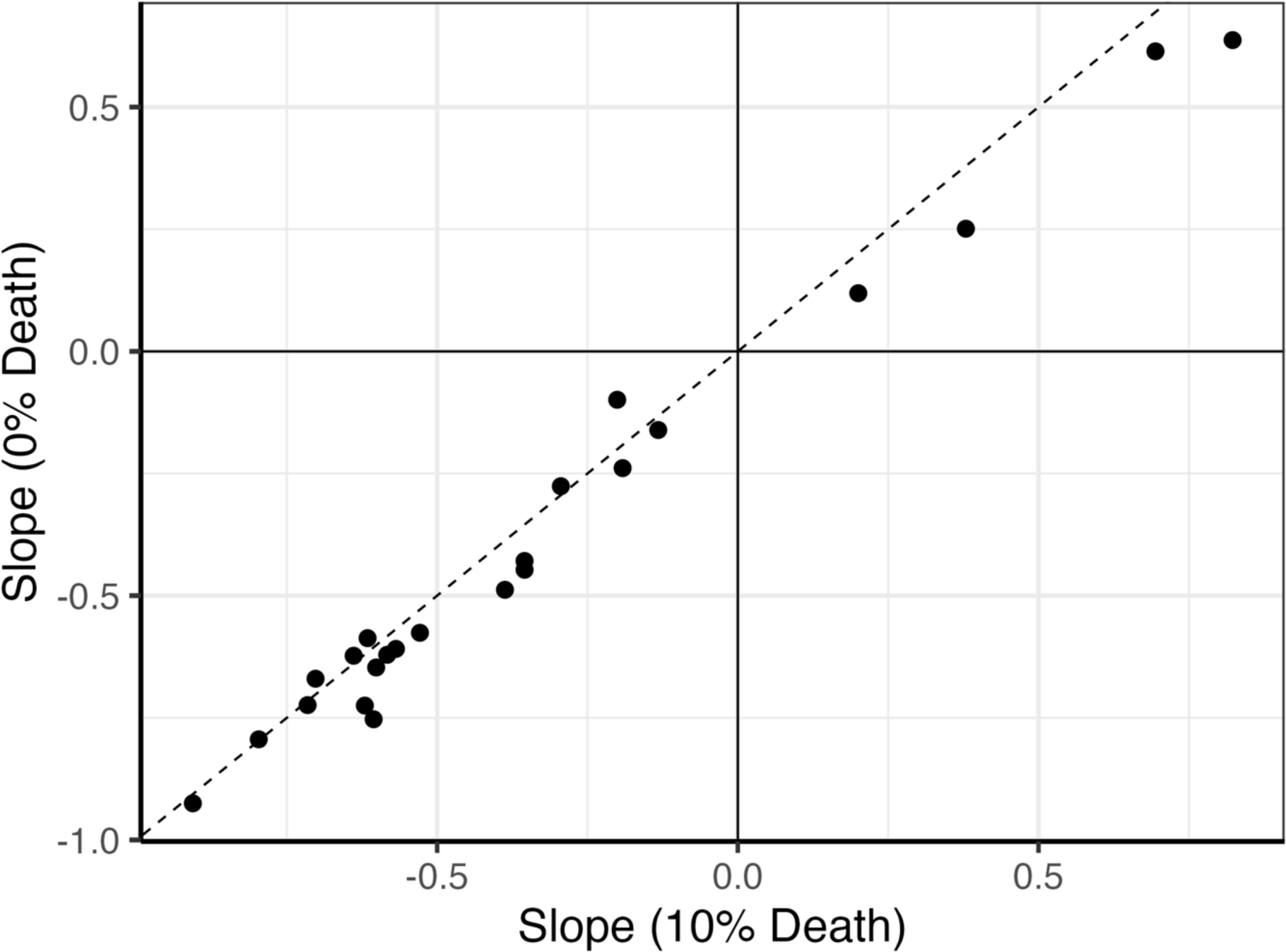
An assumption of 10% death has little effect on the estimation of DAMP slope. Solid lines indicate a slope of 0 (no DAMP), dashed line shows identical slope values for both estimates. All treatments remain in the same category (DAMP, no DAMP or reverse DAMP).

**Figure S14.**
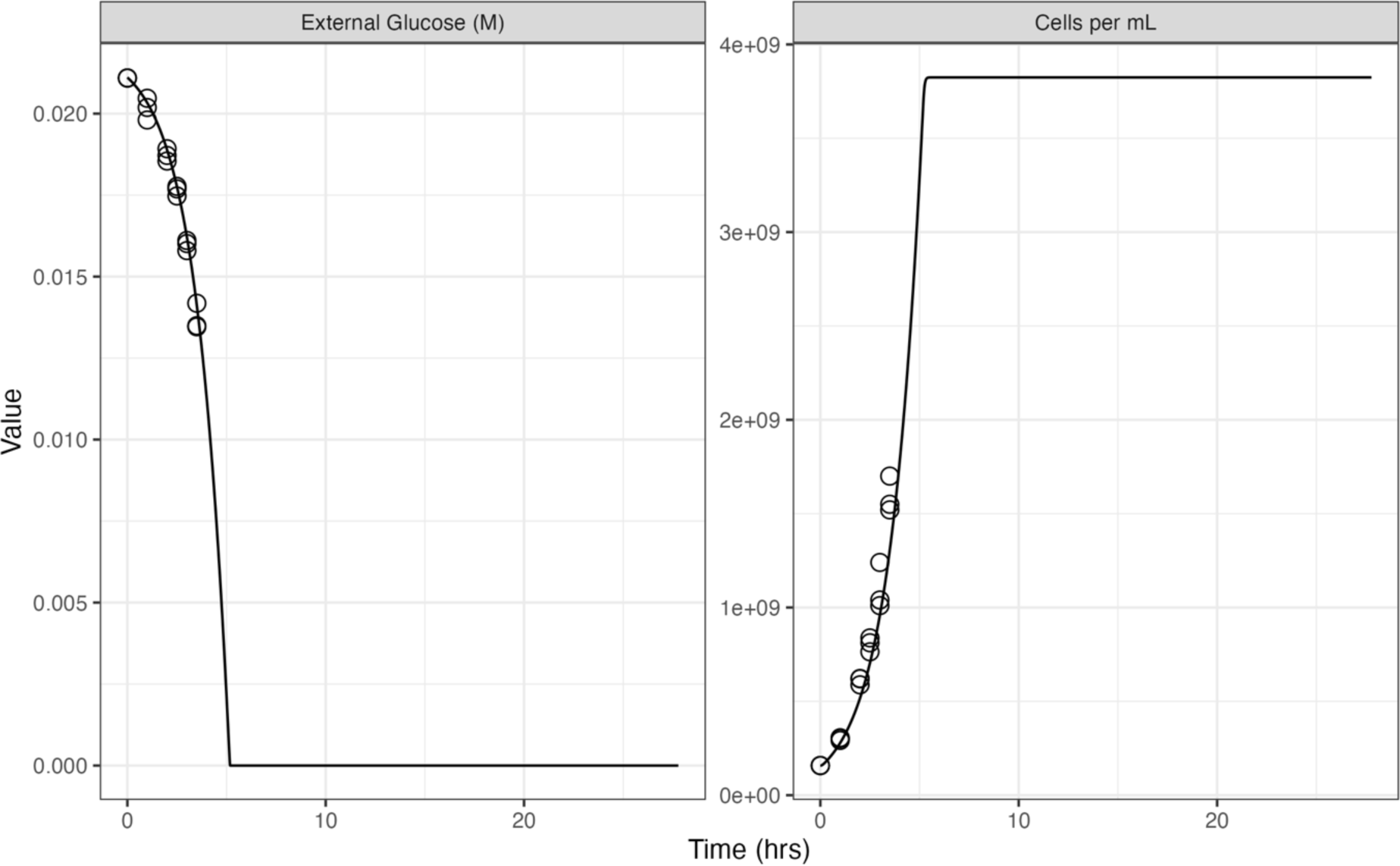
Fit of model variant A to published data. Lines show results of ODE model A and circles show data from (81) used to fit parameters U1 and M1. Left-hand panel shows the molar concentration of external glucose over time and right-hand panel shows E. coli cells per mL over time.

**Figure S15.**
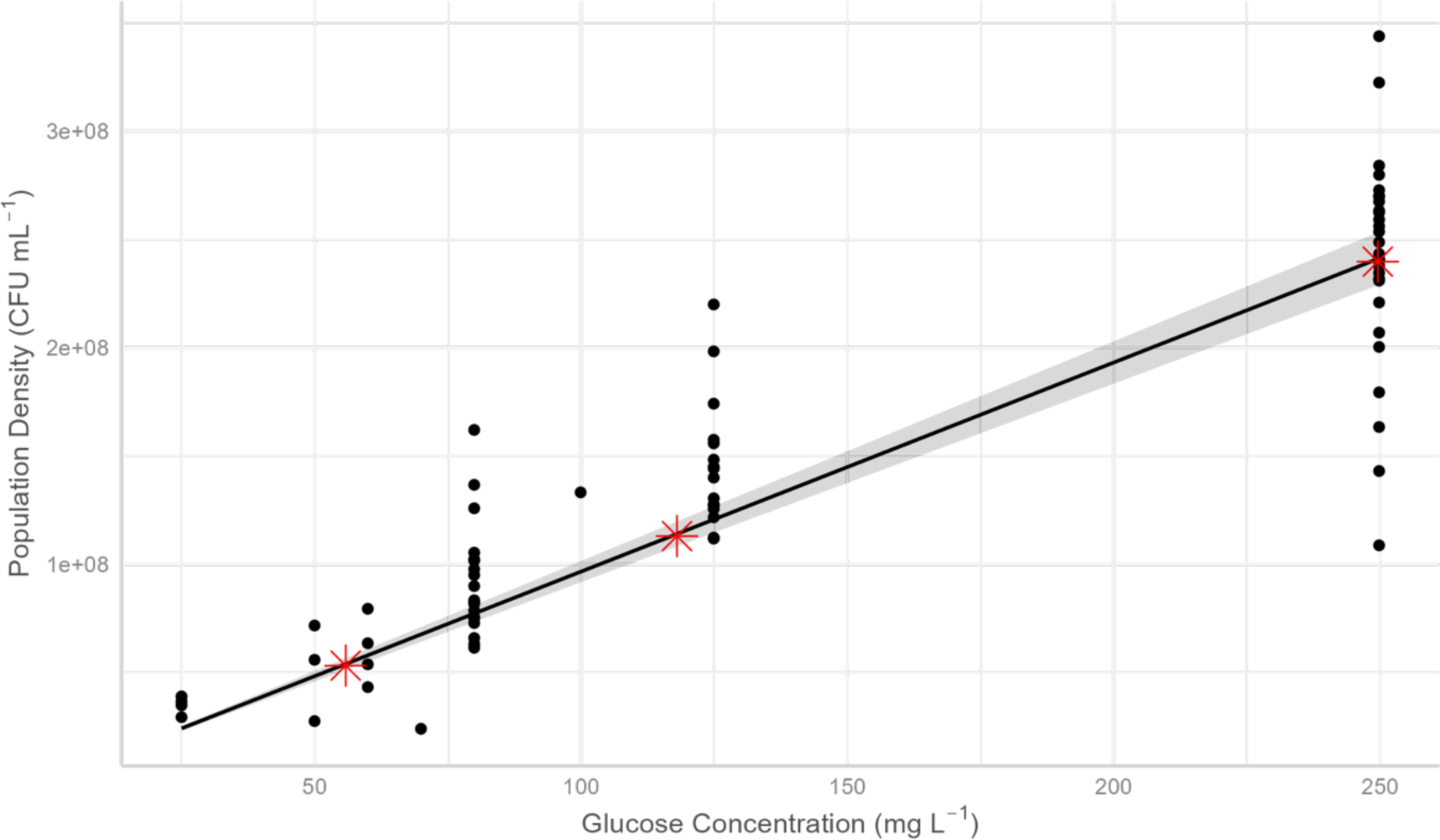
Fit of linear mixed effects model relating final population density to initial glucose concentration. Used to fit parameter Met1 in ODE models. Black points show published lab data from (3) on population density and glucose provision in E. coli MG1655 used to fit this regression. Black line and shaded area show fitted relationship and 95% confidence interval respectively of a mixed effects model accounting for random effects of experimental block and plate. Red stars show output, in final population density, from initial ODE model A under differing initial glucose concentrations.

**Table S1:**
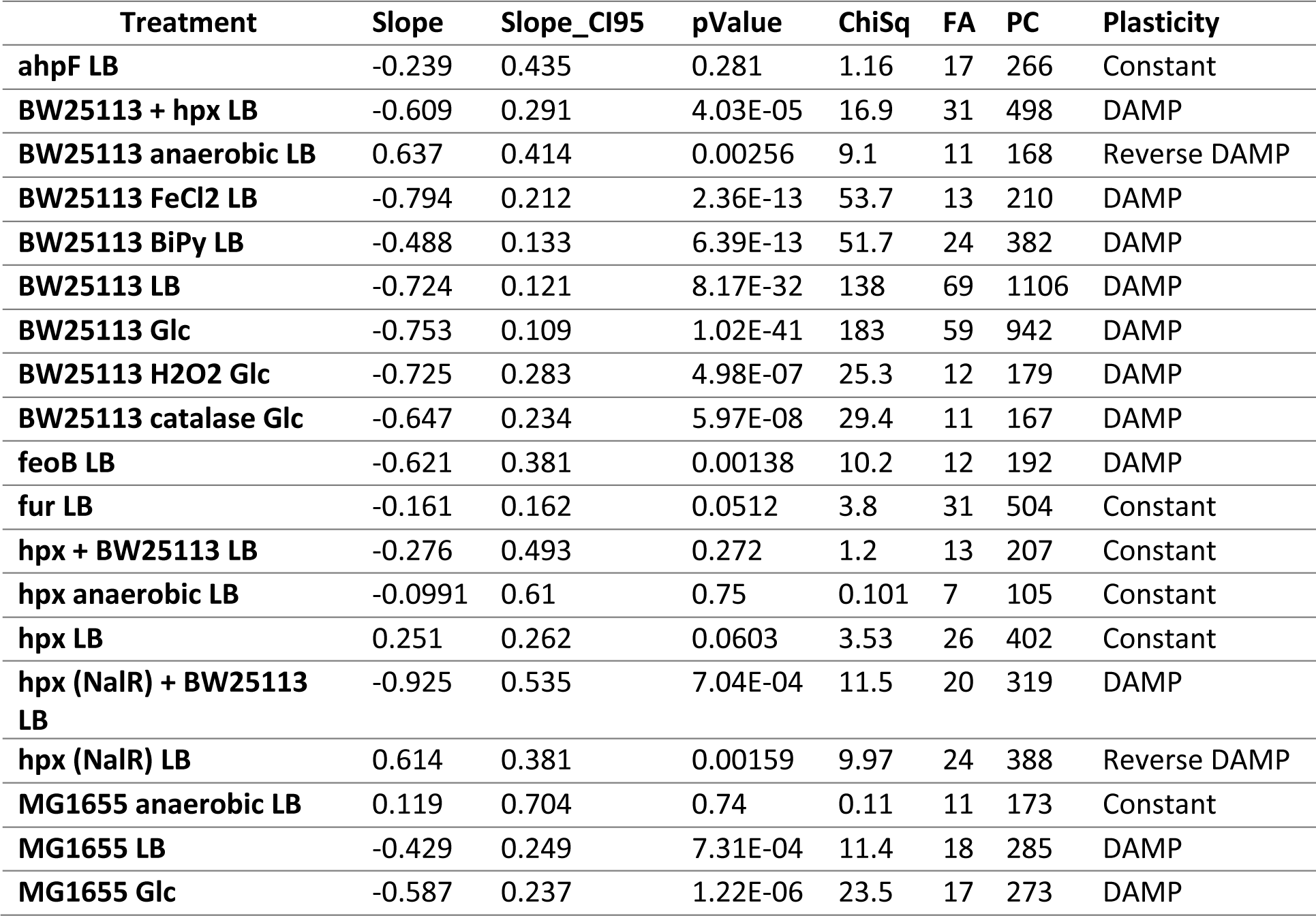
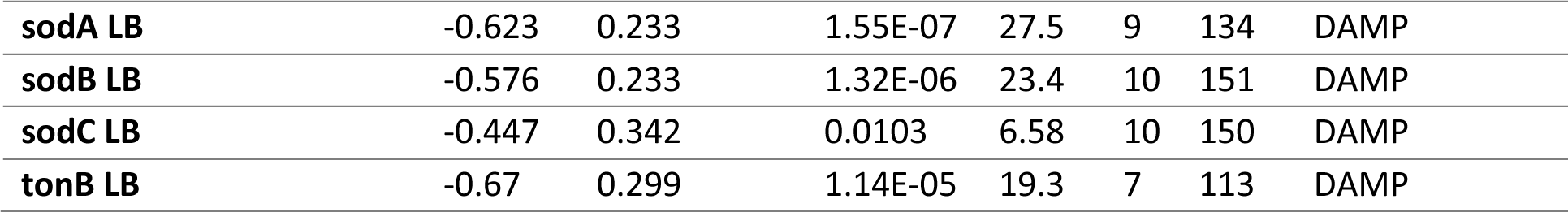
Slope estimates with associated Chi-Squared tests from Regression 4 (SI). Slope indicates the log-log relationship between population density and mutational events per mL minus 1 (1 is subtracted to make interpretation simpler as a constant mutation rate is now defined by a slope of 0 rather than a slope of 1). Slope_CI95 indicates that a 95% confidence interval on the slope estimate will be slope ± slope_CI95. pValue is calculated from a Chi-Squared test (DF=1) comparing the original slope value to the Null Hypothesis that the slope of the given treatment = 1 (slope = 1 when mutation rate is constant with respect to population density); therefore in treatments in which the slope significantly differs from 1 we have observed density associated mutation rate plasticity. FA and PC list the number of fluctuation assays and parallel cultures used in the analysis respectively. Plasticity shows if the treatment shows a constant mutation rate, DAMP (a significant inverse relationship between population density and mutation rate) or reverse DAMP (a significant direct relationship between population density and mutation rate).

**Table. S2:**
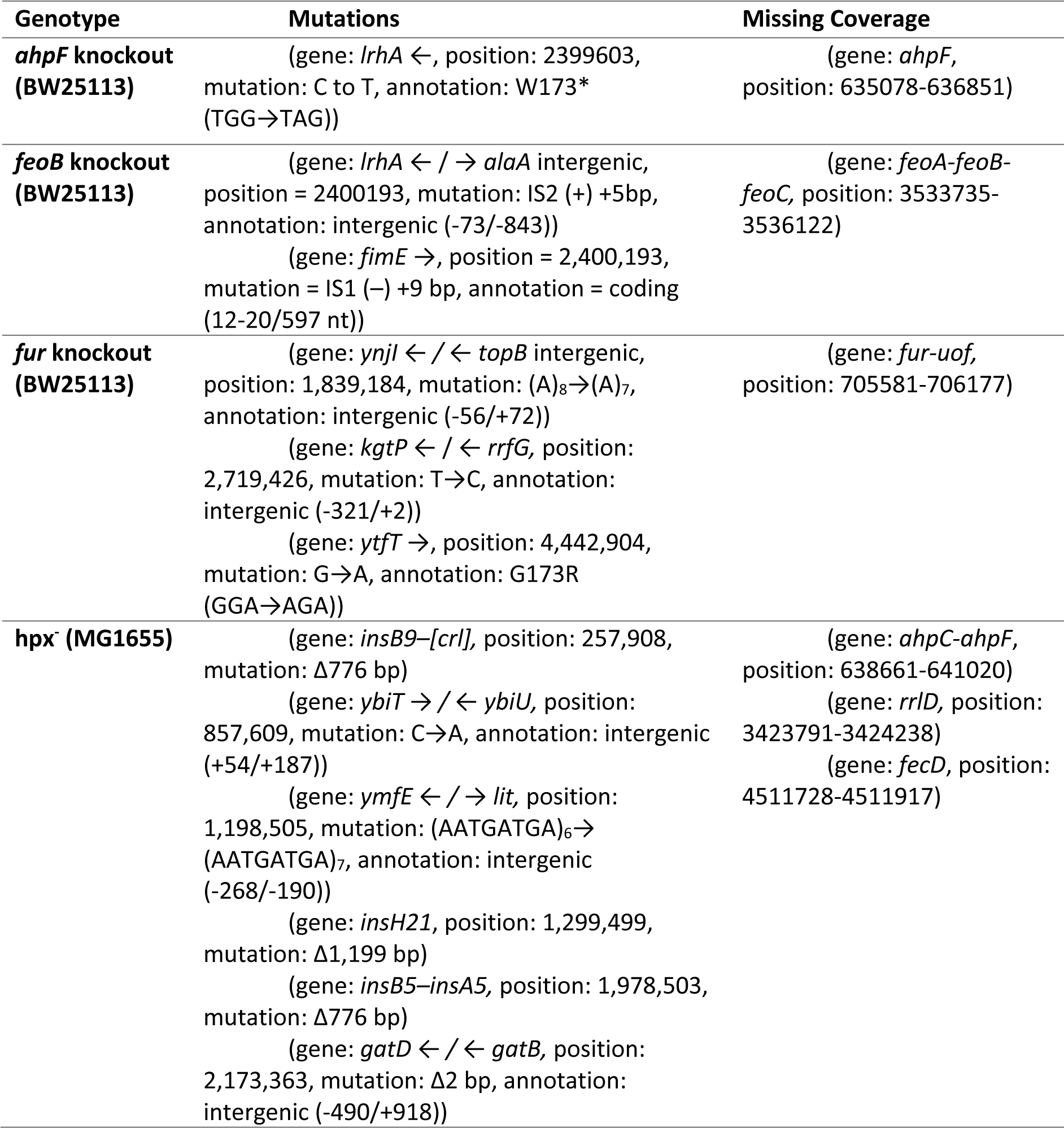
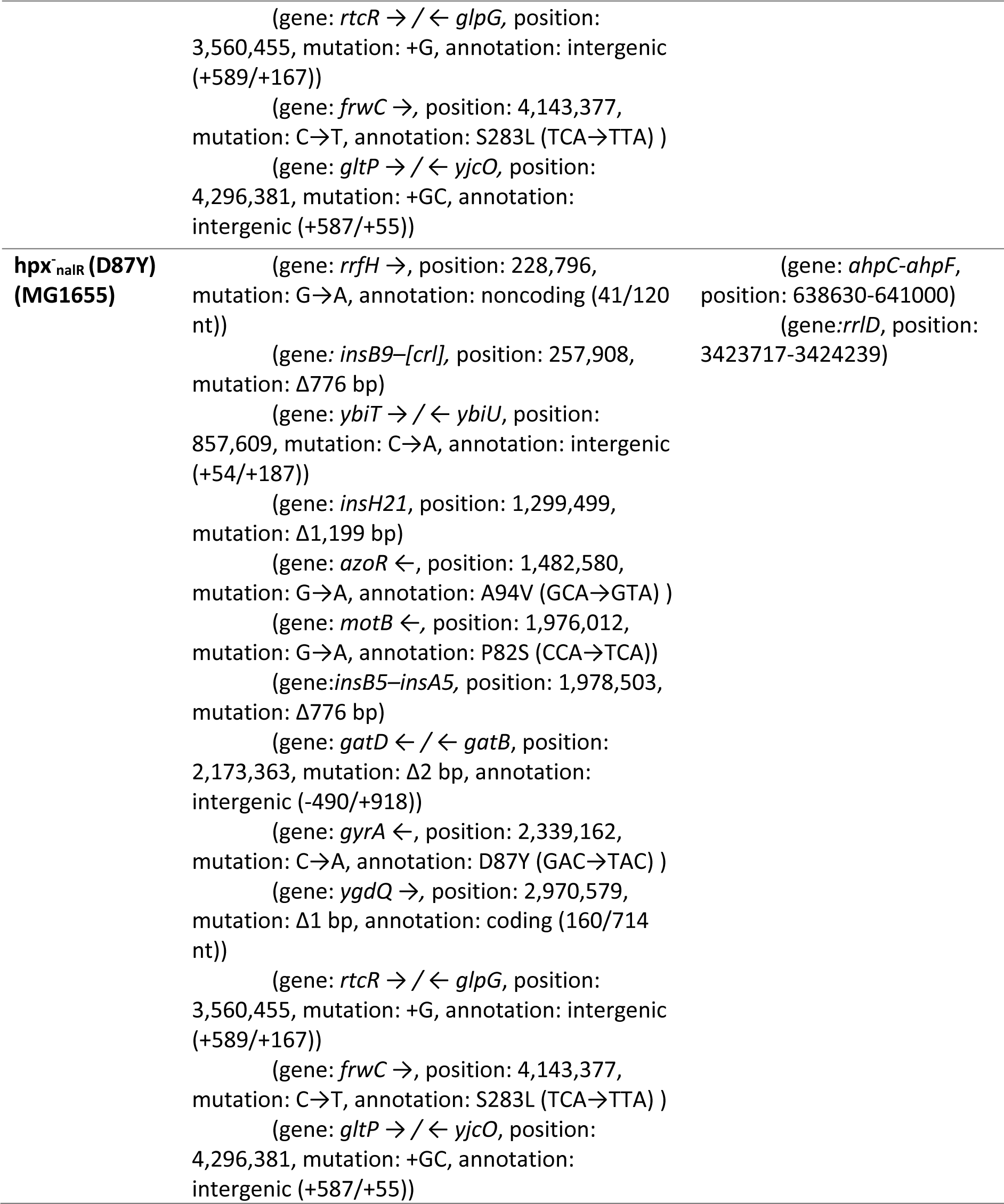
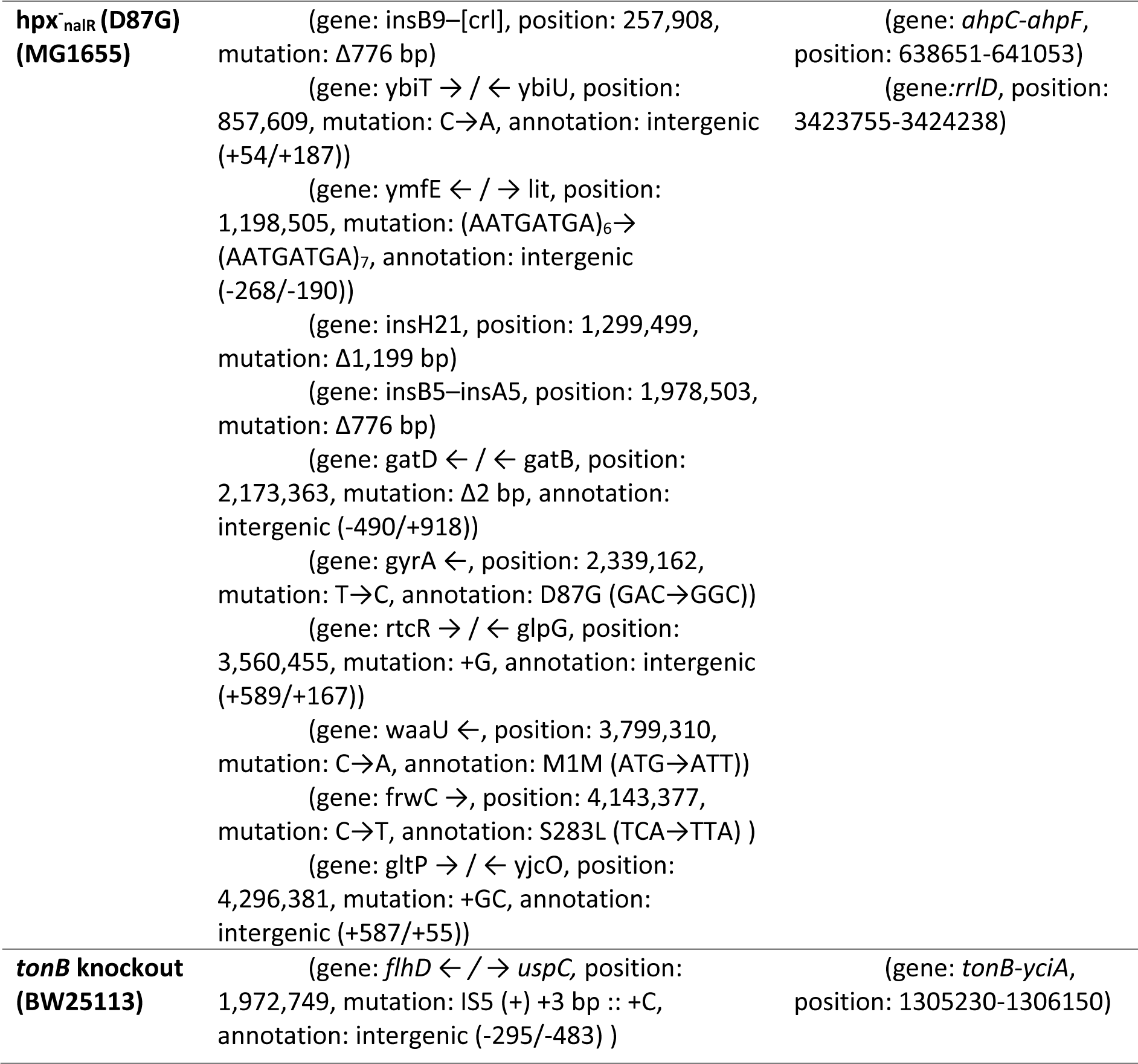
Mutations, missing coverage and new junction evidence for key strains in this study as predicted by variant calling with breseq.

**Table. S3:**
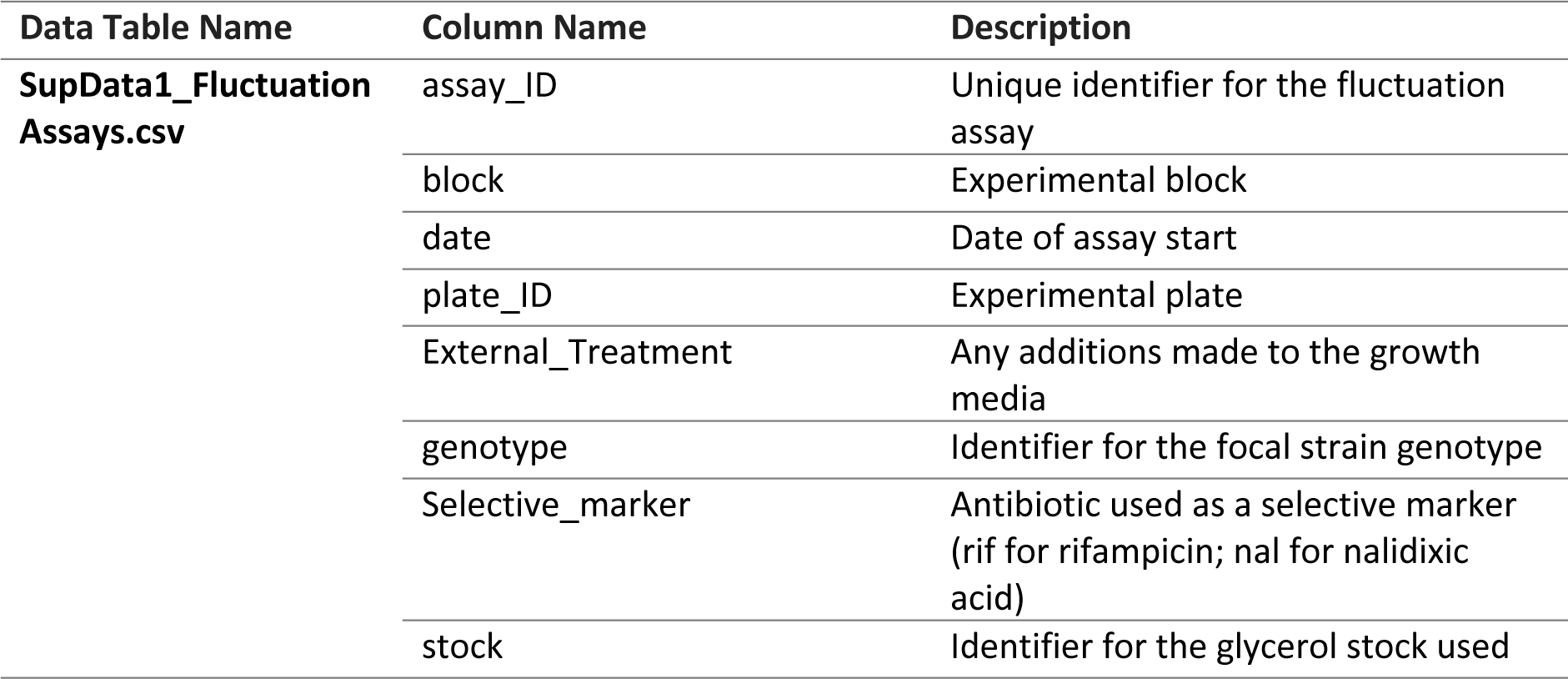
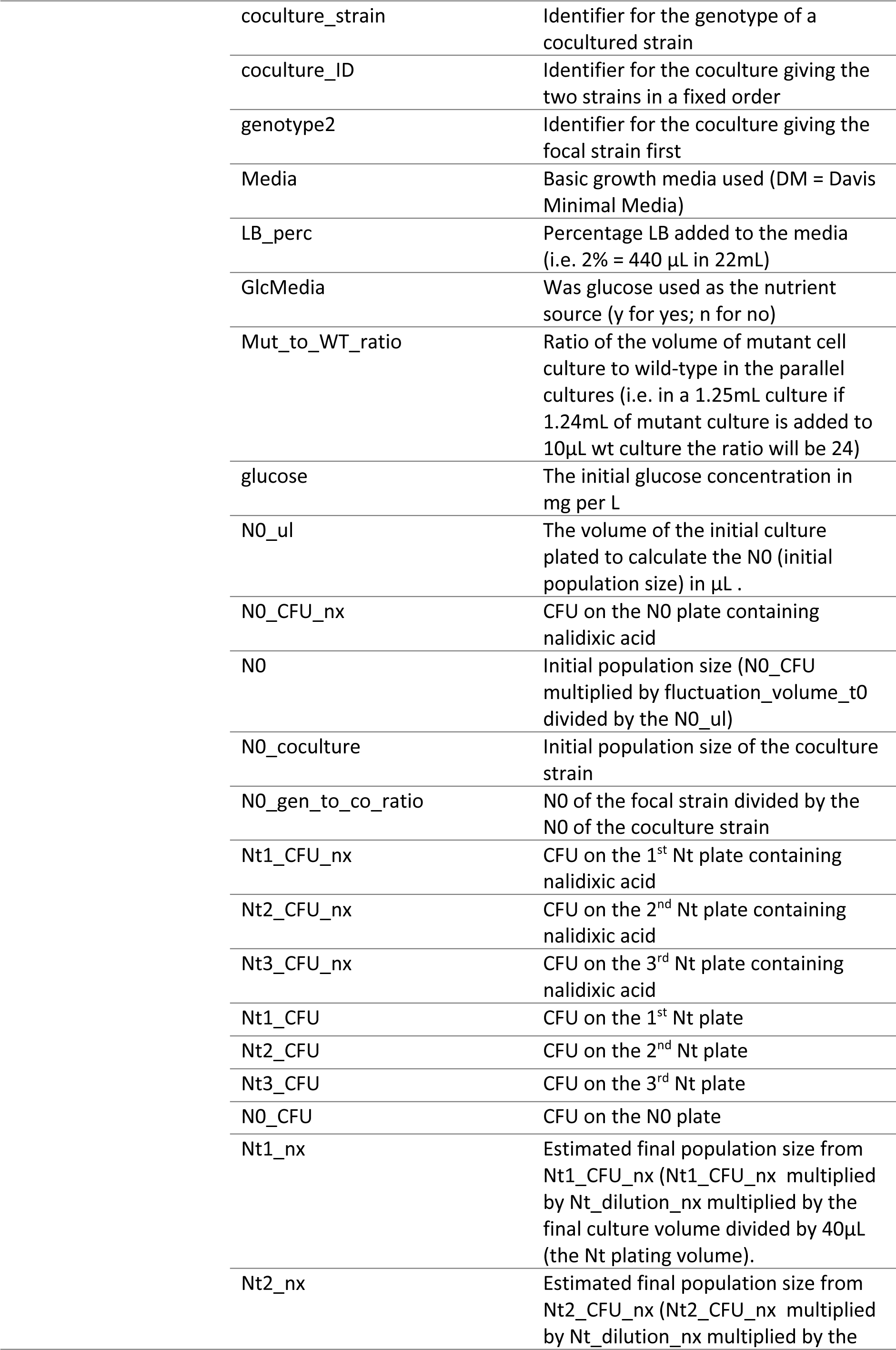
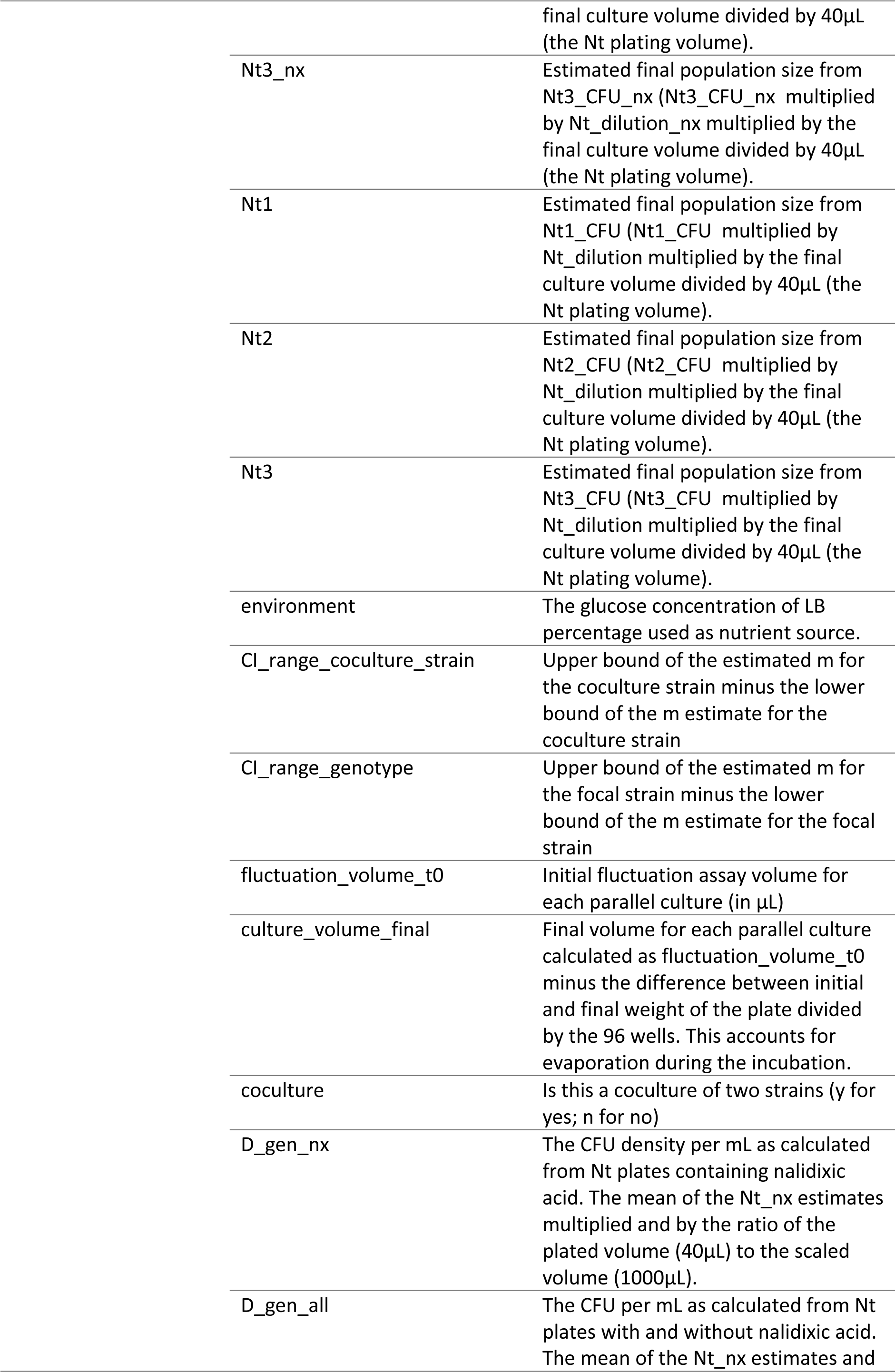
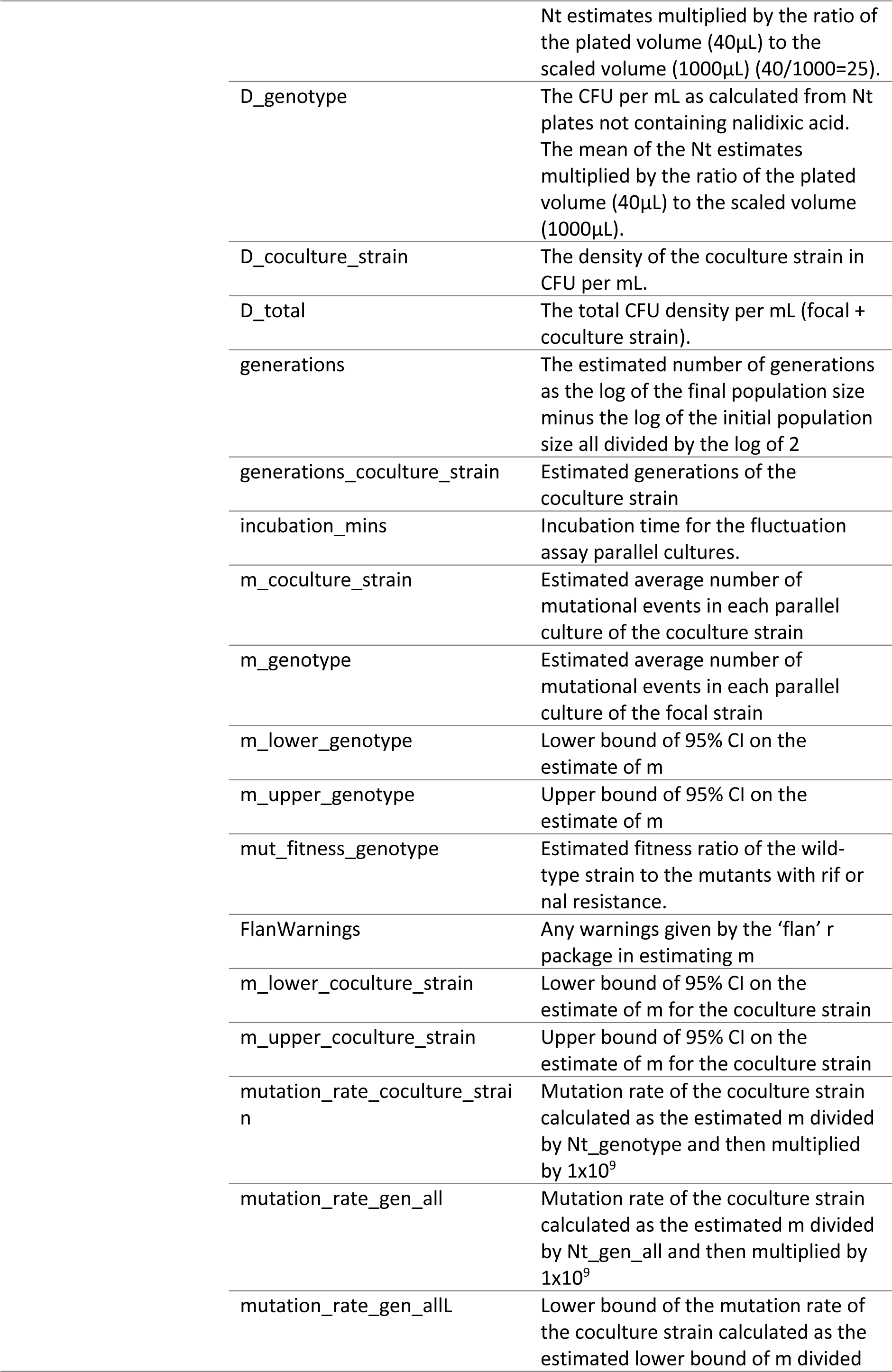
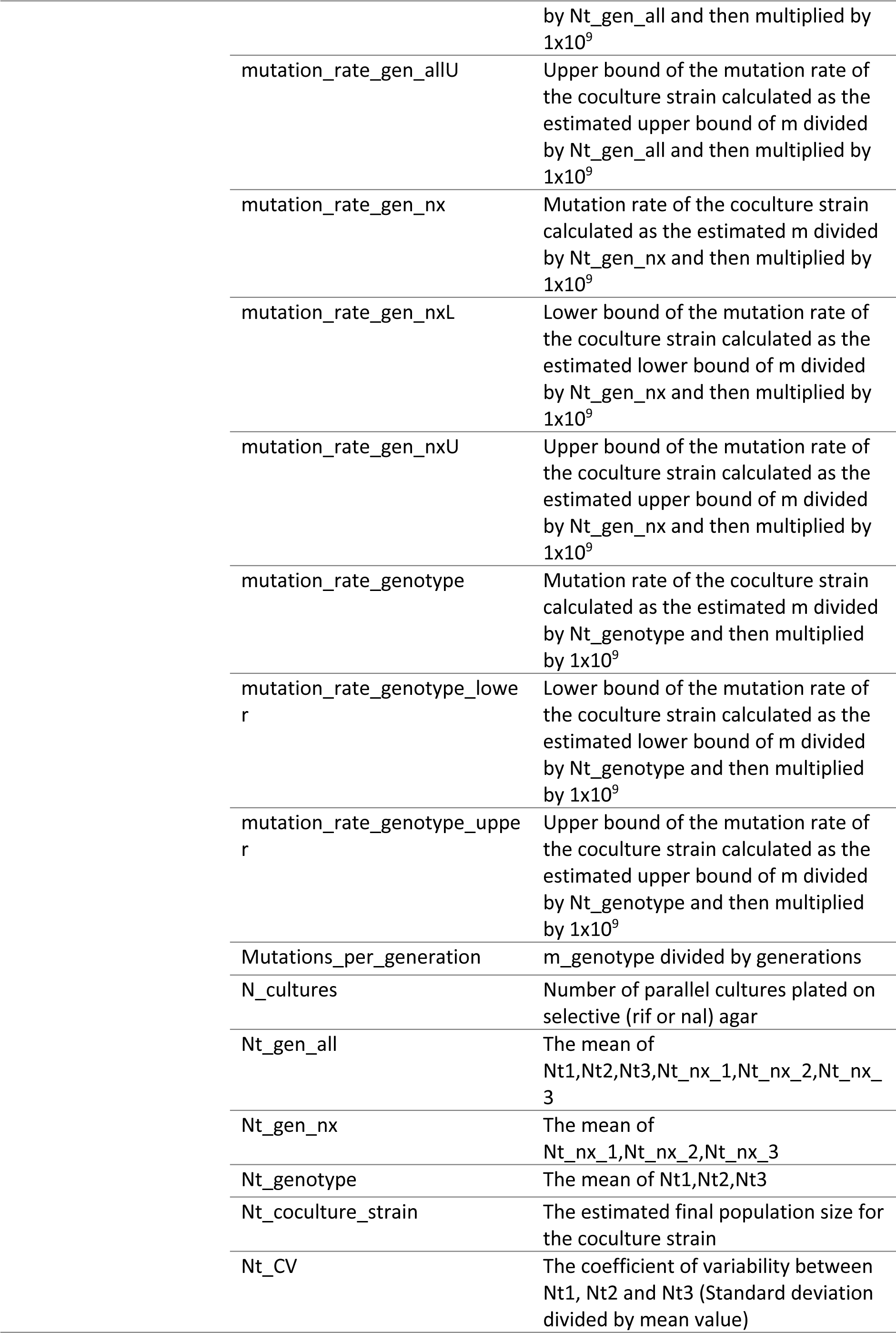
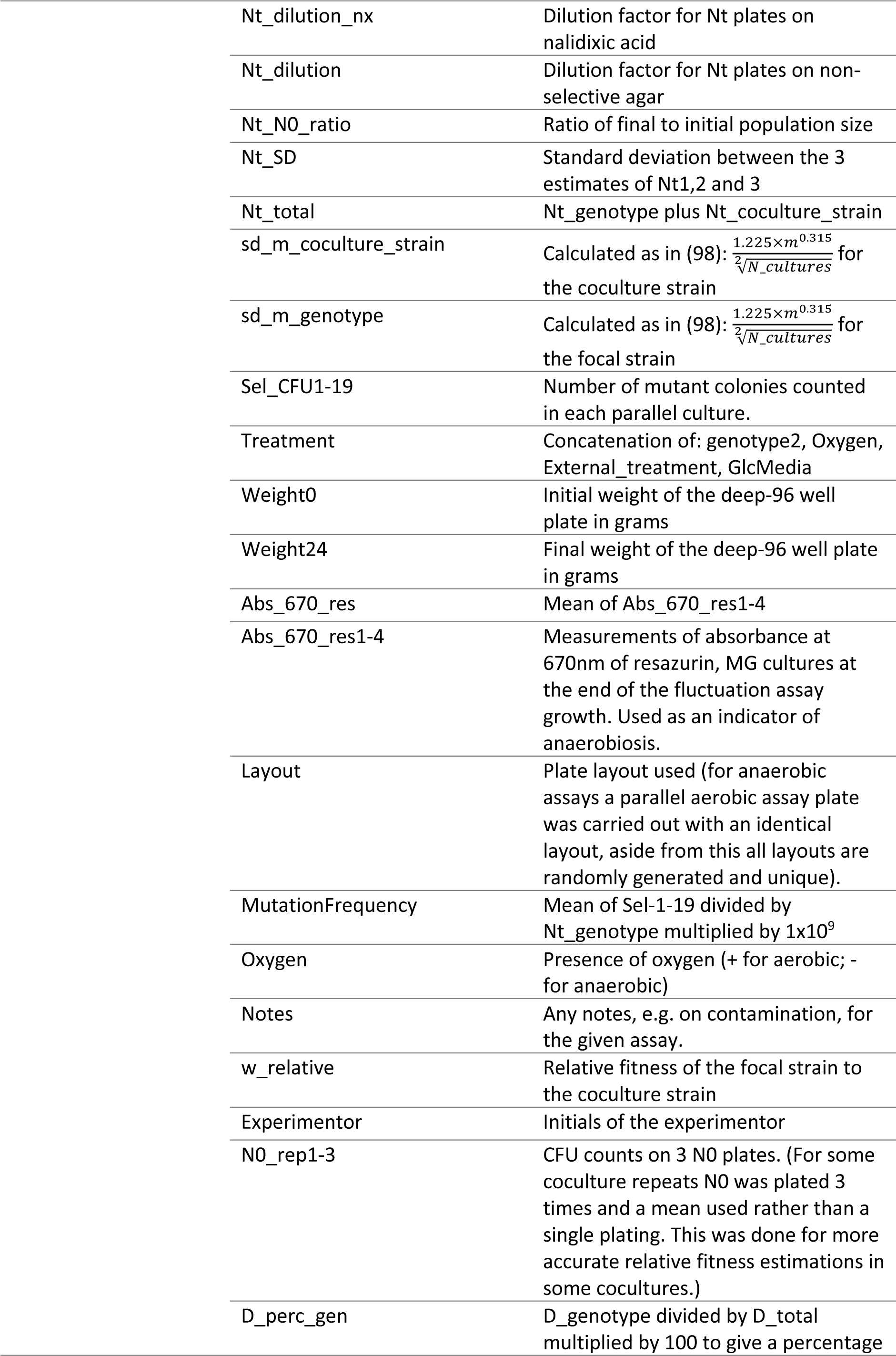
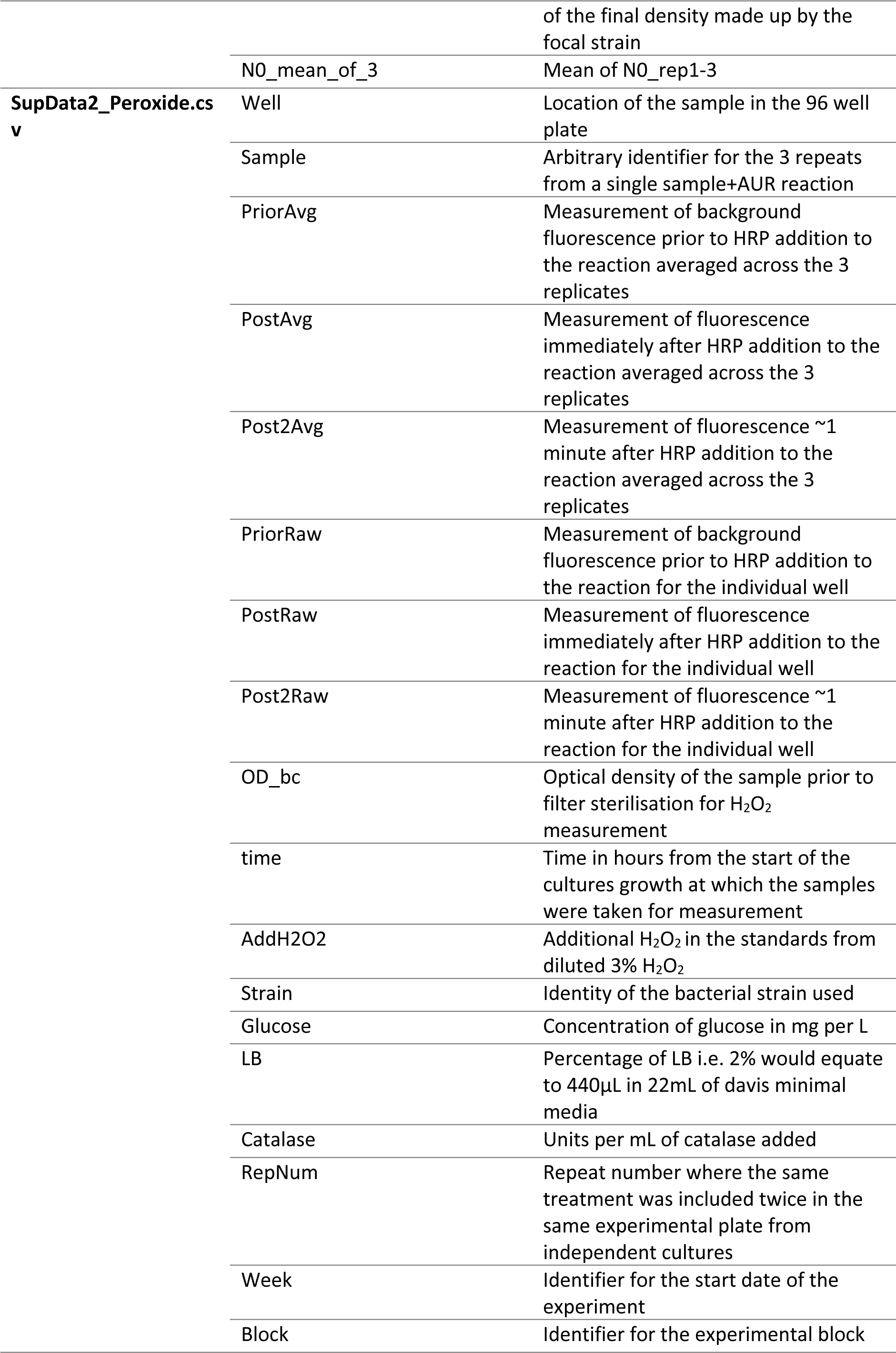
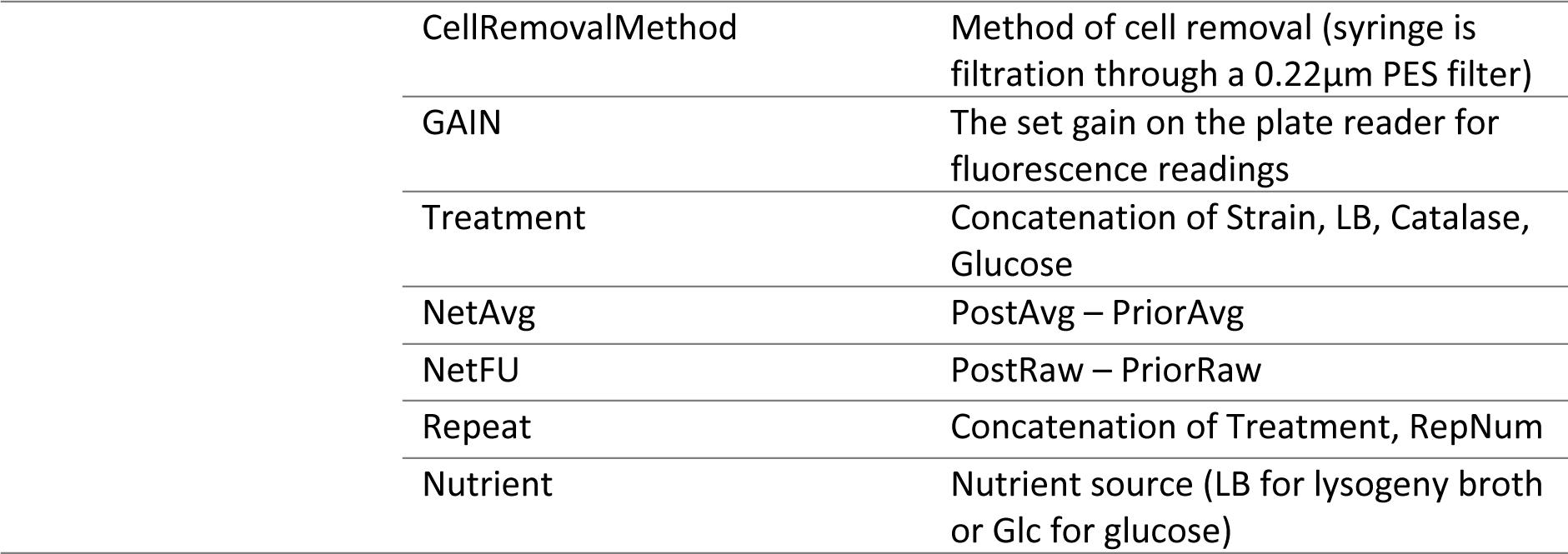
Descriptions for columns in supplementary data tables SupData2_Peroxide.csv and SupData1_FluctuationAssays.csv.

## Supplementary Files

**00_SupCode1.R:** R code necessary to recreate ODE modelling (Fig. 1, 2, S1, S2, S14, S15)

**00_SupCode2.R:** R code necessary to recreate lab work analysis (Fig. 3, 4, S3, S4, S5, S6, S7, S8, S9, S10, S11, S12, S13)

**SupplementaryStats.docx:** Details of statistical models used in this study.

**SupData1_FluctuationAssays.csv:** Data from fluctuation assays needed to run 00_SupCode2.R

**SupData2_Peroxide.csv:** Data from amplex ultra-red peroxide assays needed to run 00_SupCode2.R

**SupData3_Jain09.txt:** Data available from (81) used to fit U1 parameter in 00_SupCode1.R

**SupData4_Krasovec2017.txt:** Data available from (3) used to fit Met1 parameter in 00_SupCode1.R

