## Supplementary Statistics for "Working together to control mutation: how collective peroxide detoxification determines microbial mutation rate plasticity"

Supplementary Text - Statistical Models and Tests for Green et al., 2023

**Regression 1 (Fig. 2)**

Simulated mutation rates across 5 population densities in ODE models A-K (with log_2_ transformation) are fitted as a function of population density (with log_2_ transformation).

**ANOVA table for Regression 1**

|  | Df | Sum Sq | Mean Sq | F value | Pr(>F) |
| --- | --- | --- | --- | --- | --- |
| log2(wt) | 1 | 6.63 | 6.63 | 617 | 6.23E-23 |
| model | 10 | 1.22 | 0.122 | 1.13E+01 | 4.42E-08 |
| log2(wt):model | 10 | 22.5 | 2.25 | 209 | 7.41E-27 |
| Residuals | 33 | 0.355 | 0.0108 |  |  |

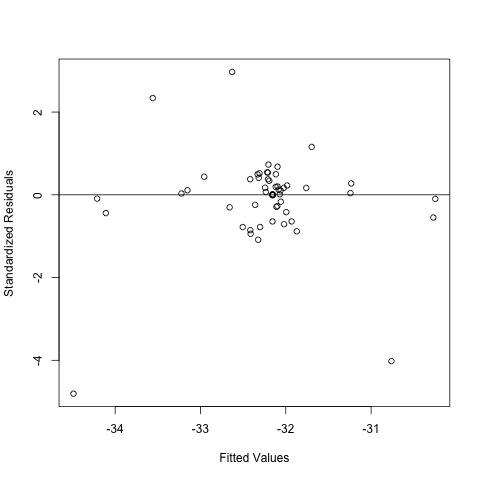

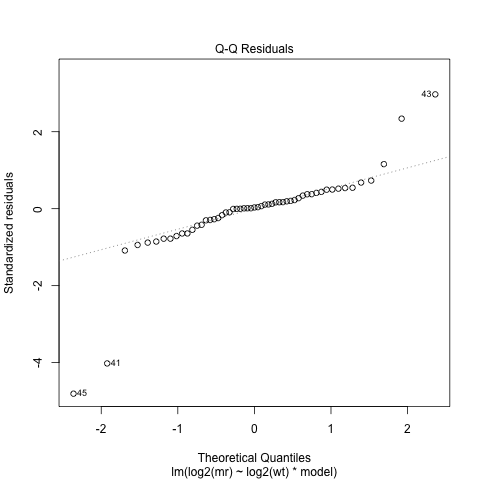

**Diagnostic plots for Regression 1.**

Standardised residuals by fitted values and normal quantile-quantile plot of standardised residuals.

**Model Regression 2 (Fig. 2)**

Mutation rates (with log_2_ transformation) of the wildtype BW25113 (ancestor) strain grown in minimal glucose media are fitted as a function of population density (also with log_2_ transformation). Random effects of experimental plate within experimental block on the intercept are included. This model uses a subset of the data used in Models S1-2.

**ANOVA table for Regression 2**

|  | numDF | denDF | F-value | p-value |
| --- | --- | --- | --- | --- |
| (Intercept) | 1 | 18 | 80491.69 | 0 |
| Log2(D_genotype) | 1 | 16 | 150.0677 | 1.52E-09 |

**Variance and Standard Deviations for Random Effects for Regression 2**

|  | Variance | StdDev |
| --- | --- | --- |
| block | 8.27E-02 | 2.88E-01 |
| plate_ID | 1.80E-09 | 4.24E-05 |
| Residual | 2.03E-01 | 4.50E-01 |

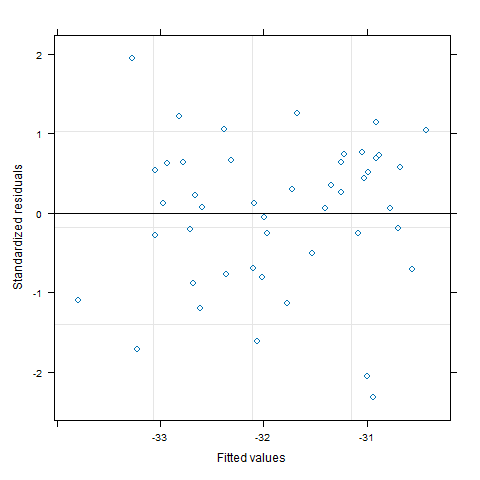

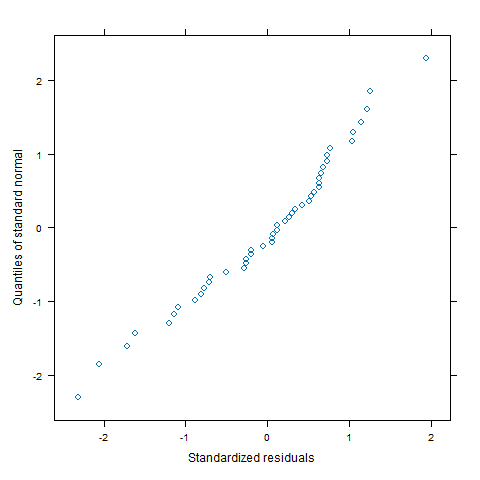

**Diagnostic plots for Regression 2.**

Standardised residuals by fitted values and normal quantile-quantile plot of standardised residuals.

**Regression 3 – (Fig S3)**

The model fitting the fitness effect of resistance during fluctuation assays is the log_2_ transformed predicted fitness effect as a function of the treatment (genotype, coculture strain and growth conditions; 47 levels). Random effects of experimental plate (159 levels) nested within experimental block (57 levels) nested within experimenter (5 levels), each affecting the intercept, are also fitted.

**ANOVA table for Regression 3**

|  | numDF | denDF | F-value | p-value |
| --- | --- | --- | --- | --- |
| (Intercept) | 1 | 485 | 36 | 3.78E-09 |
| Treatment | 46 | 485 | 3.19 | 1.56E-10 |

**Variance and Standard Deviations of Random Effects for Regression 3**

|  | Variance | StdDev |
| --- | --- | --- |
| Experimentor | 1.48E-01 | 3.84E-01 |
| block | 2.23E-09 | 4.72E-05 |
| plate_ID | 9.79E-10 | 3.13E-05 |
| Residual | 9.53E-01 | 9.76E-01 |

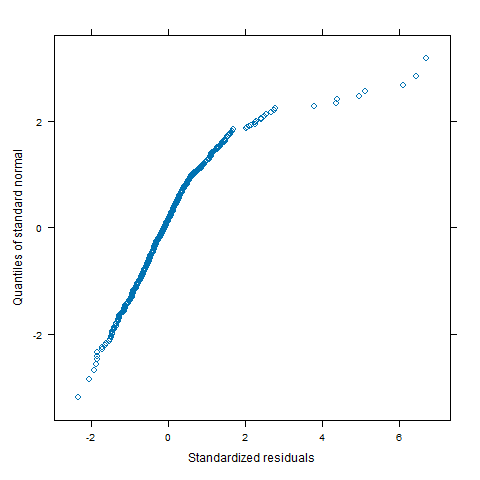

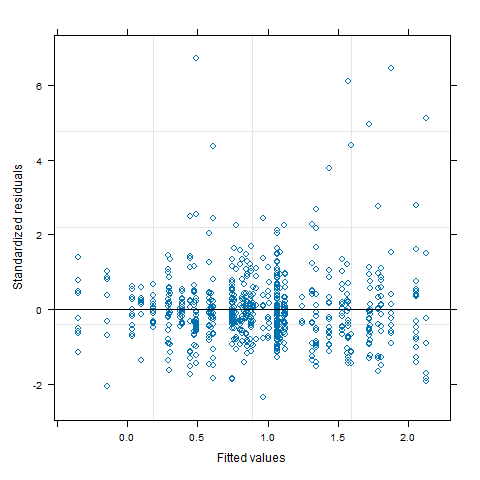

**Diagnostic plots for Regression 3.**

Standardised residuals by fitted values and normal quantile-quantile plot of standardised residuals.

**Regression 4 (Fig. 3-4)**

The model shown in fig. 3 and fig. 4 fits log_2_ mutational events per mL against mean-centred log_2_ density allowing for differences in intercept and slope among the 47 treatments. A series of variants of this model was constructed allowing differences in variance (i.e. heteroscedasticity) associated with one or two covariates.

Potential variance covariates considered were: experimental block, genotype, external treatment (e.g. anaerobiosis or chelator), date, experimental plate, selective marker (i.e. rifampicin or nalidixic acid), nutrient type, nutrient concentration and experimenter identity, all treated as discrete effects with a different variance at each level. Continuous variance covariates allowed variance to change as a power function of the covariate. Potential continuous variance covariates considered were: the fitted values of the response variable, the percentage of LB in the media, the concentration of glucose in the media, the initial population size (N0), the range between 95% confidence intervals on the mutation rate, the average final volume of the parallel cultures, density (estimated by colony forming units), the number of generations (log(final population)-log(initial population))/(log(2)), the estimated number of mutational events (m), the lower and upper bounds and standard deviation in that estimate, the estimated fitness ratio of cells with to without the selective marker, the mutation rate (m/final population size), the upper and lower bounds on the mutation rate, the number of parallel cultures, the final population size, the standard variation and coefficient of variation in that estimate, the ratio of final to initial population sizes, the initial and final weight of the experimental plate and the incubation time of the experimental plate.

The model variant with the lowest AIC was then chosen. In this case the model allowed variance to change with the upper bound of m and with genotype. The two slopes and intercepts of hpx^-^_nalR_ strains D87Y and D87G were very similar, and this model was therefore further simplified (improving AIC further) by combining these effects to estimate a single intercept and slope value.

**ANOVA table for Regression 4**

|  | numDF | denDF | F-value | p-value |
| --- | --- | --- | --- | --- |
| (Intercept) | 1 | 447 | 93.1 | 0 |
| Dc | 1 | 447 | 1680 | 0 |
| TreatmentHC | 44 | 447 | 41.7 | 0 |
| Dc:TreatmentHC | 44 | 447 | 6.29 | 0 |

**Variance and Standard Deviations of Random Effects for Regression 4**

|  | Variance | StdDev |
| --- | --- | --- |
| Experimentor | 0.33313575 | 0.5771791 |
| block | 0.06456309 | 0.2540927 |
| plate_ID | 0.01259386 | 0.1122224 |
| Residual | 0.61839495 | 0.7863809 |

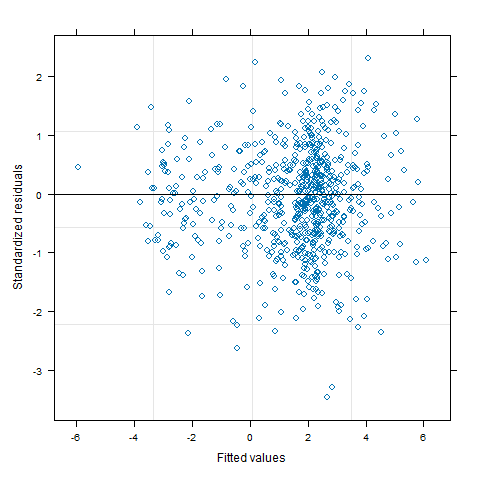

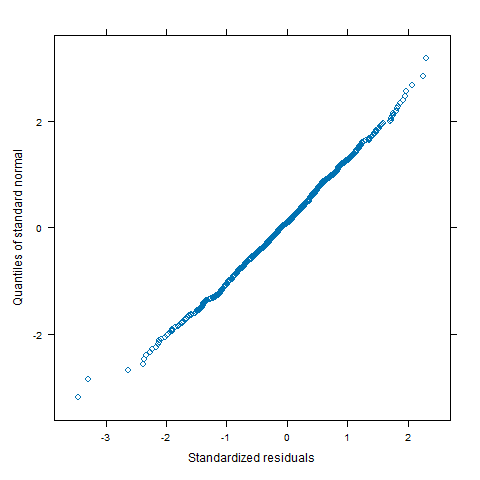

**Diagnostic plots for Regression 4.**

Standardised residuals by fitted values and normal quantile-quantile plot of standardised residuals.

**Regression** **5**

To estimate hydrogen peroxide concentration from arbitrary fluorescence units (AFU) in the amplex ultra-red assay, a standard curve was fitted with known hydrogen peroxide concentrations. H_2_O_2_ added in the standard is fitted as a function of net fluorescence (final fluorescence minus initial fluorescence or fluorescence of a 0-concentration standard). A random effect of block affecting the intercept and slope (NetFU) was also included.

**ANOVA table for Regression 5**

|  | numDF | denDF | F-value | p-value |
| --- | --- | --- | --- | --- |
| (Intercept) | 1 | 112 | 254 | 0 |
| NetFU | 1 | 112 | 90.3 | 4.44E-16 |

**Variance and Standard Deviations and Covariance for Random Effects for Regression 5**

|  | Variance | StdDev | Corr |
| --- | --- | --- | --- |
| (Intercept) Block | 1.45E+00 | 1.21E+00 | (Intr) |
| NetFU | 5.84E-09 | 7.64E-05 | 0.75 |
| Residual | 6.19E-01 | 7.87E-01 |  |

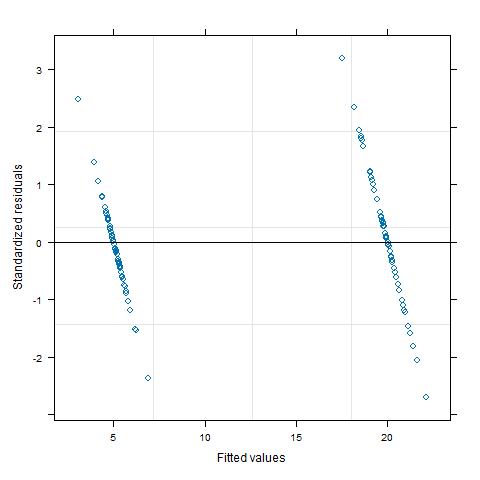

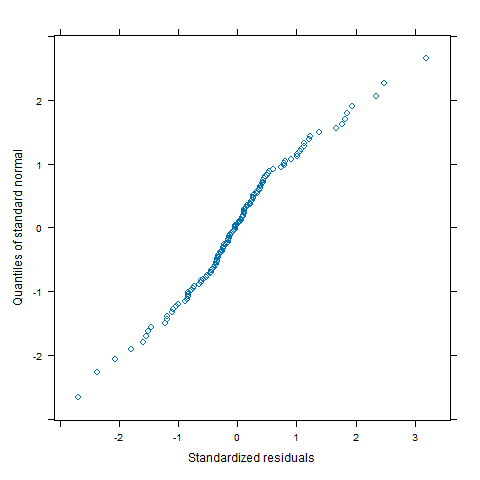

**Diagnostic plots for Regression 5.**

Standardised residuals by fitted values and normal quantile-quantile plot of standardised residuals.

**Regression 6 – Fig. S6B**

Predicted H_2_O_2_ (in μM units) using regression 5 is fitted as a function of nutrient type, nutrient level and presence or absence of cells. Initially all interactions between effects were included with the three-way interaction between all effects and the interaction between nutrient type and nutrient level then removed, improving the fit of the model. Random effects on the intercept of reaction tube nested within block nested within week are also included.

**ANOVA table for Regression 6**

|  | numDF | denDF | F-value | p-value |
| --- | --- | --- | --- | --- |
| (Intercept) | 1 | 120 | 41.80638 | 2.27E-09 |
| Nutrient | 1 | 46 | 0.868673 | 0.356189 |
| Strain | 1 | 46 | 63.42915 | 3.34E-10 |
| NutLev | 1 | 46 | 0.032108 | 0.858579 |
| Nutrient:Strain | 1 | 46 | 3.61737 | 0.063452 |
| Strain:NutLev | 1 | 46 | 9.801188 | 0.003028 |

**Variance and Standard Deviations for Random Effect for Regression 6**

|  | Variance | StdDev |
| --- | --- | --- |
| Week.. | 4.15E+00 | 2.037065416 |
| Block.. | 8.15E-08 | 0.0002855606 |
| Tube.. | 2.01E+00 | 1.42E+00 |
| Residual | 4.67E-02 | 2.16E-01 |

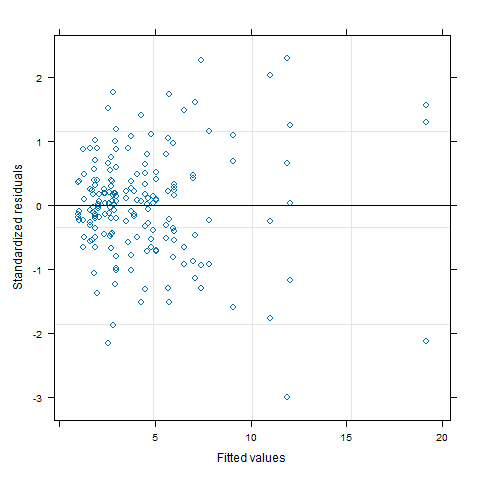

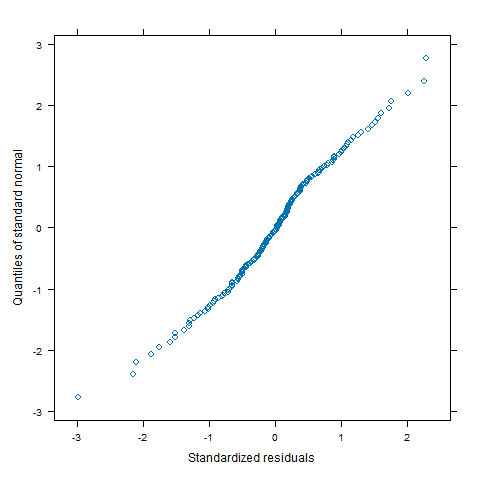

**Diagnostic plots for Regression 6.**

Standardised residuals by fitted values and normal quantile-quantile plot of standardised residuals.

**Regression** **7 – Fig. S6A**

Predicted H_2_O_2_ (in μM units) using regression 5 is fitted as a function of cell density (estimated by absorbance at 600nm). Initially interaction between the effects was included (reg 7A) this was then removed along with the effect of nutrient type, improving the fit of the model (reg 7B). Random effects on the intercept of reaction tube nested within block were included.

**ANOVA table for Regression 7A**

|  | numDF | denDF | F-value | p-value |
| --- | --- | --- | --- | --- |
| (Intercept) | 1 | 72 | 32.39773 | 2.54E-07 |
| OD_bc | 1 | 26 | 23.11608 | 5.58E-05 |
| Nutrient | 1 | 26 | 0.770591 | 0.388074 |
| OD_bc:Nutrient | 1 | 26 | 0.307902 | 0.583713 |

**ANOVA table for Regression 7B**

|  | numDF | denDF | F-value | p-value |
| --- | --- | --- | --- | --- |
| (Intercept) | 1 | 72 | 30.25295 | 5.48E-07 |
| OD_bc | 1 | 28 | 24.32518 | 3.34E-05 |

**Variance and Standard Deviations for Random Effects for Regression 7B**

|  | Variance | StdDev |
| --- | --- | --- |
| Block.. | 1.87371015 | 1.3688353 |
| Tube.. | 0.37484601 | 0.6122467 |
| Residual | 0.02630368 | 0.1621841 |

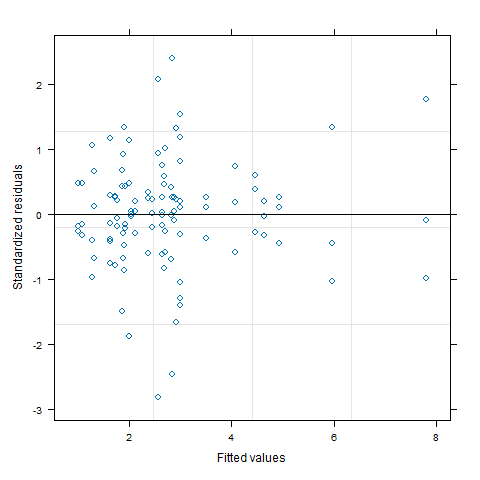

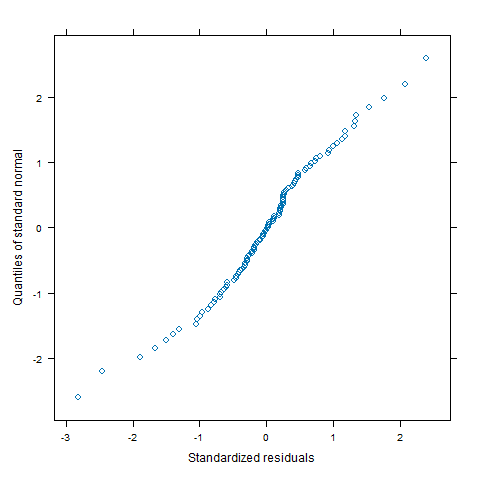

**Diagnostic plots for Regression 7B**

Standardised residuals by fitted values and normal quantile-quantile plot of standardised residuals.

**Regression** **8**

Regression 8 fits log2 mutation rate of the hpx^-^_nalR_ strains against mean-centred log2 total density (estimated via CFU) initially allowing for differences in intercept and slope among the four treatments (different strain/coculture combinations). Combining the two hpx^-^_nalR_ strains significantly improves the fit of the model (as with regression 4) and so only the effect of coculture on intercept and slope is used. Random effects of experimental plate nested within experimental block on the intercept are included.

**ANOVA table for Regression** **8**

|  | numDF | denDF | F-value | p-value |
| --- | --- | --- | --- | --- |
| (Intercept) | 1.00E+00 | 29 | 1090 | 0.00E+00 |
| log2(D_total) | 1.00E+00 | 29 | 57.2 | 2.43E-08 |
| coculture | 1.00E+00 | 2.90E+01 | 16.3 | 0.000363 |
| log2(D_total):coculture | 1.00E+00 | 2.90E+01 | 14.4 | 0.000704 |

**Variance and Standard Deviations for Random Effects for Regression** **8**

|  | Variance | StdDev |
| --- | --- | --- |
| Block | 1.81E-10 | 1.35E-05 |
| plate_ID | 1.41E-10 | 1.19E-05 |
| Residual | 5.68E-01 | 7.54E-01 |

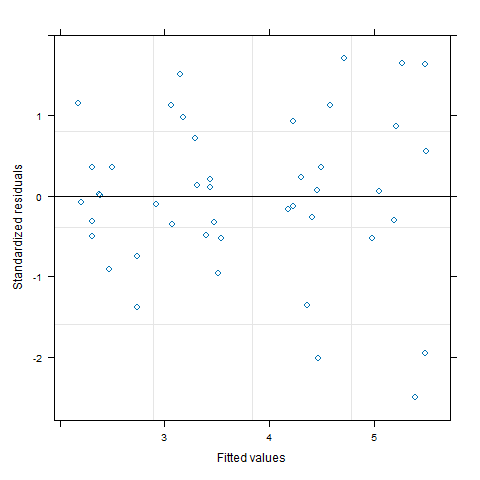

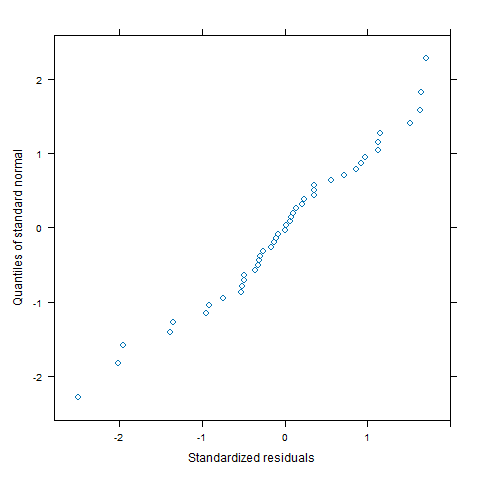

**Diagnostic plots for Regression** **8.**

Standardised residuals by fitted values and normal quantile-quantile plot of standardised residuals.

**Regression** **9A**

Regression 9 fits the plating efficiency of the hpx^-^_nalR&rifR_ strains used in a reconstruction test as a function of the plating environment and the strain identity, the random effect of the 6-well agar plate on which the cultures were plated is also included. Regression 9B is identical but the 4 cell treatments (hpx^-^ and wt low high and medium density) are combined to one level.

**ANOVA table for Regression 9A**

|  | numDF | denDF | F-value | p-value |
| --- | --- | --- | --- | --- |
| (Intercept) | 1 | 109 | 993 | 0 |
| Treatment | 6 | 109 | 45.2 | 0.00E+00 |
| Strain | 1 | 109 | 127 | 0 |

**Variance and standard deviations for random effects for Regression 9A**

|  | Variance | StdDev |
| --- | --- | --- |
| (Intercept) | 0.005355174 | 0.07317906 |
| Residual | 2.49E-02 | 0.15777969 |

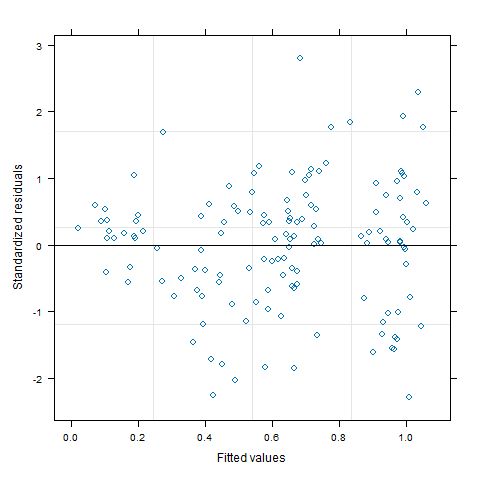

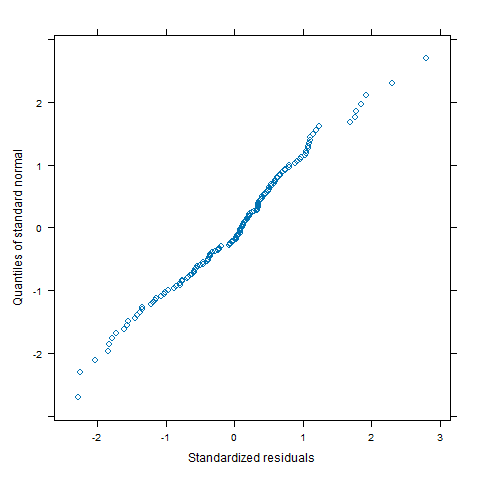

**Diagnostic plots for Regression 9A.**

Standardised residuals by fitted values and normal quantile-quantile plot of standardised residuals.

**ANOVA comparing Regression 9A to Regression 9B**

|  | call | df | AIC | BIC | Log  Lik | Test | L.Ratio | p-value |
| --- | --- | --- | --- | --- | --- | --- | --- | --- |
| Reg 9A | lme.formula(fixed = Efficiency ~ Treatment + Strain, data = ReconData, random = ~1 \| Plate, method = "ML") | 10 | -88.5 | -59.1 | 54.3 |  |  |  |
| Reg 9B | lme.formula(fixed = Efficiency ~ CombT + Strain, data = ReconData, random = ~1 \| Plate, method = "ML") | 7 | -94.2 | -73.6 | 54.1 | 1 vs 2 | 0.308 | 0.959 |
